# Circadian regulation of the transcriptome in a complex polyploid crop

**DOI:** 10.1101/2021.09.30.462369

**Authors:** Hannah Rees, Rachel Rusholme-Pilcher, Paul Bailey, Joshua Colmer, Benjamen White, Connor Reynolds, Sabrina Jaye Ward, Calum A. Graham, Luíza Lane de Barros Dantas, Antony N. Dodd, Anthony Hall

## Abstract

The circadian clock is a finely balanced time-keeping mechanism that coordinates programmes of gene expression. It is currently unknown how the clock regulates expression of homoeologous genes in polyploids. Here, we generate a high-resolution time-course dataset to investigate the circadian balance between sets of three homoeologous genes (triads) from hexaploid bread wheat. We find a large proportion of circadian triads exhibit imbalanced rhythmic expression patterns, with no specific sub-genome favoured. In wheat, period lengths of rhythmic transcripts are found to be longer and have a higher level of variance than in other plant species. Expression of transcripts associated with circadian controlled biological processes are largely conserved between wheat and *Arabidopsis*, however striking differences are seen in agriculturally critical processes such as starch metabolism. Together, this work highlights the ongoing selection for balance versus diversification in circadian homoeologs, and identifies clock-controlled pathways that might provide important targets for future wheat breeding.

## Introduction

Circadian clock homologs have been both inadvertently selected during crop domestication and identified as crop improvement targets^1–4^. Understanding circadian regulation of the transcriptome in crops such as bread wheat (*Triticum aestivum*) may provide useful insights for future crop improvement. Wheat also provides an excellent model system to explore how the circadian clock and its outputs are co-ordinated in a recently formed, complex allopolyploid. In *Arabidopsis*, circadian transcription factors act in a dose-dependent manner, with both knock-out and over-expression mutants resulting in altered function of the circadian oscillator^5–8^. It is not yet understood how rhythmic gene expression is balanced in species with multiple copies of the same gene. *T. aestivum* is a hexaploid (AABBDD) formed through interspecific hybridisation of three diploid ancestors around 10,000 years ago^9, 10^. 51.7% of high-confidence wheat genes still exist in triads; sets of three homoeologous genes present on each of the A, B and D genomes^11^. As these homoeologs evolved independently for several million years prior to hybridization, it is plausible that these independent species might have been subject to different selective pressures on their clocks (Fig. 1a). The circadian network in *Arabidopsis* comprises a series of interlocking negative transcriptional feedback loops connected by key activators^12^. Although monocots such as wheat diverged from their dicot relatives over 140 million years ago^13^, many circadian oscillator components seem to have been conserved, particularly those forming the core loop network. Orthologs of *TIMING OF CAB EXPRESSION 1 (TOC1*) and other *PSEUDO-RESPONSE REGULATOR (PRR)* genes have been identified in wheat, rice and barley, and several loci within these genes have been associated with altered flowering times, most notably *(ppd-1*) within *TaPRR3/7*^14–16^. Likewise, mutants of orthologs of *LATE ELONGATED HYPOCOTYL (LHY)*, *GIGANTEA* (*GI*), *EARLY FLOWERING 3 (ELF3),* and *LUX ARRYTHMO (LUX)* have been identified that alter heading dates, pathogen susceptibility, plant height or lower grain yields^17–21^.

**Fig. 1.**
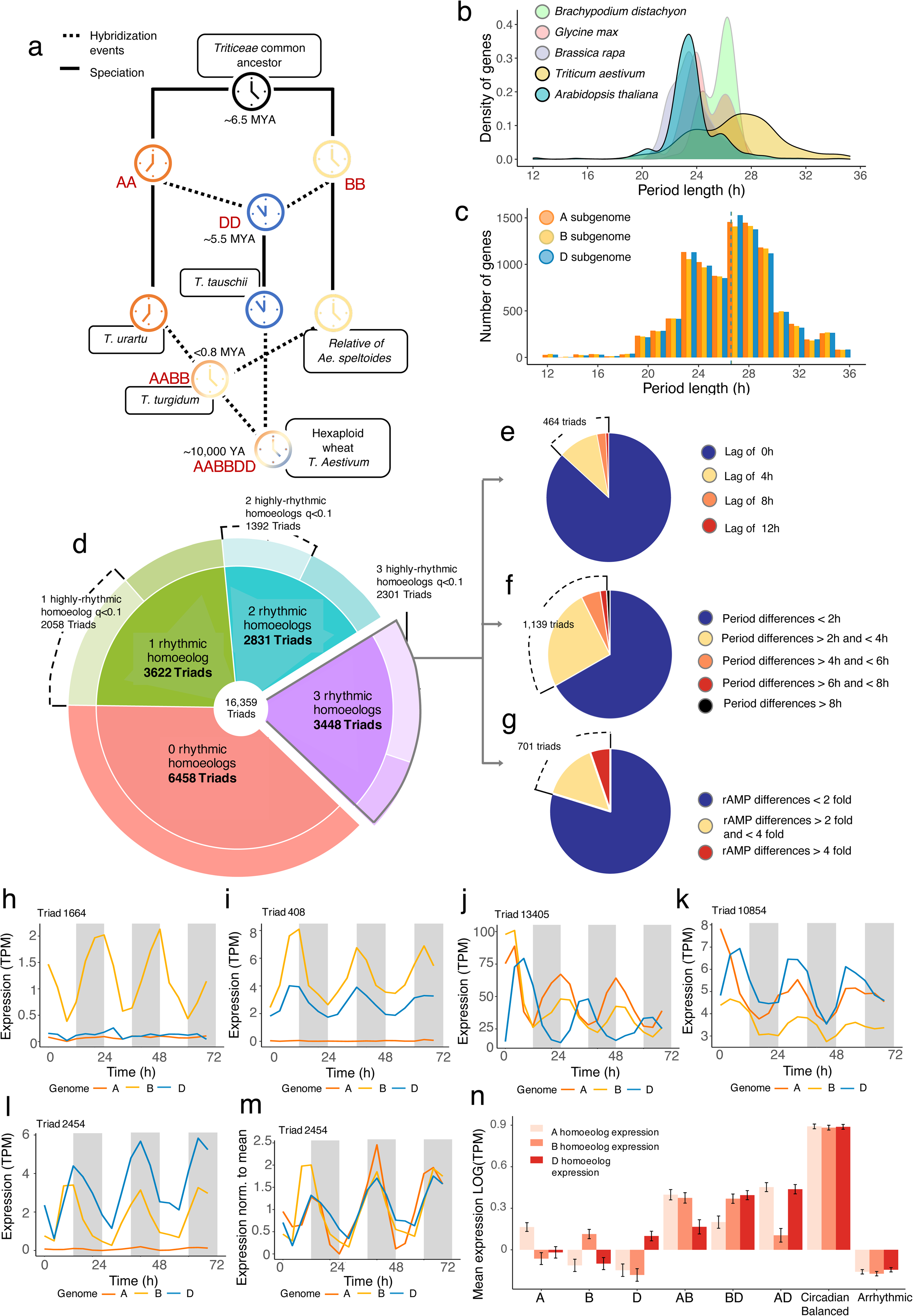
Circadian regulation of homoeolog expression of wheat triads. **a,** Schematic of the origins of hexaploid wheat, showing circadian clocks evolving independently in the ancestors of the A, B and D subgenomes following divergence from a common ancestor approximately 6.5 million years ago. Colours of clock icons represent theoretical differences in clock regulation integrated in the tetraploid and hexaploid hybrids either through circadian balance or through dominance of a particular homoeolog copy. Speciation and hybridisation event dates are based on estimates from^108^. **b,** Density plot showing the distribution of period lengths across rhythmic transcripts (BH *q* < 0.01) in *Arabidopsis*, *Brassica rapa*, *Brachypodium distachyon*, *Glycine max* (Soybean) and wheat based on meta2d estimates on 24-68h data following transfer to constant light. **c,** Histogram showing distribution of period lengths in wheat split between the A, B and D subgenomes. Dotted line indicates the mean period for the A, B and D subgenomes. **d,** Proportions of triads with either zero (red segment), one (green segment), two (blue segment) or three (purple segment) rhythmic gene(s) out of the 16,359 expressed triads in this dataset. Lighter shading in the outer segments represents cases where one/two homeolog(s) have high confidence rhythmicity (BH *q* < 0.01) alongside an arrhythmic homeolog (BH *q* >0.05). We term these genes “imbalanced rhythmicity” triads. Of the 3448 triads with three rhythmic genes (represented by the purple segment in d), we also looked for triads with circadian imbalance in: phase (**e**), period (**f**) or relative amplitude (**g**). 464 triads had homoeologs which peaked with an optimum lag of 4, 8 or 12h following cross-correlation analysis. 1,139 triads had homoeologs with period differences of more than 2h. 701 triads had homoeologs with a more than two-fold difference in relative amplitude. **h,i,** Example triads for imbalanced rhythmicity, where either one or two homoeologs are rhythmic respectively. **j,** Example triad where the D genome homeolog lags by 8h. **k,** Example of a triad where the A genome homoeolog has a period estimate 4h longer than the D genome homoeolog. **l,** Example triad where the relative amplitude of the D-genome homoeolog is more than four times that of the A-genome homoeolog. **m,** The rhythmicity of all three homoeologs in **l,** is evident when the expression is mean normalized. **n,** Mean expression of transcripts across all timepoints in the A, B and D subgenomes within imbalanced rhythmicity triads compared with circadian balanced and arrhythmic triads. Error bars represent standard error. Circadian statistics are meta2d estimates from data 0-68h after transfer to L:L. Data represent the mean of three biological replicates with transcript expression collapsed to gene level. Genes in example triads are: [Triad 1664: TraesCS3A02G177600, TraesCS3B02G207400, TraesCS3D02G183200], [Triad 408: TraesCS3A02G533700, TraesCS3B02G610500, TraesCS3D02G539000], [Triad 13405: TraesCS6A02G269100, TraesCS6B02G296400, TraesCS6D02G245800], [Triad 10854: TraesCS6A02G166500, TraesCS6B02G194000, TraesCS6D02G155100] and [Triad 2454: TraesCS2A02G333000, TraesCS2B02G348800, TraesCS2D02G329900].

Circadian control of carbon fixation and starch metabolism are thought to form part of the selective advantage conferred by the clock^22, 23^. This is apparent in the *lhy^-^/cca1^-^* short period double mutant, where night-time starch levels reach exhaustion earlier compared to wild-type, triggering early onset starvation responses that reduce plant productivity^23^. Similarly, genes encoding photosynthesis-related proteins are well-established targets of the circadian clock and include the *LIGHT HARVESTING CHLOROPHYLL A/B BINDING PROTEIN* genes (*LHCB* also known as *CAB* genes) and photosystem I and II reaction centre genes^24, 25^. Here, we investigate circadian balance within wheat triads to understand how circadian control is co-ordinated in a polyploid crop with three subgenomes. Second, we examine similarities and differences between the circadian transcriptome in wheat and its distant dicot relative *Arabidopsis,* at a global level and at the level of genes encoding key pathways such as primary metabolism and photosynthesis.

## Results

### Global analysis of the circadian transcriptome in wheat

We generated a circadian RNA-seq time-course and compared it with a recently published dataset from *Arabidopsis*^26^ over 24h - 68h following transfer to constant light. Rhythmicity was assessed using Metacycle Benjamini-Hochberg (BH) *q*-values. Of the 86,567 genes expressed in wheat, 33.0% were rhythmically expressed with a BH *q* < 0.05 and 21.5% with a BH *q* < 0.01 (Supplementary Note 1, Supplementary Table 1). This was significantly lower than the proportions of rhythmically expressed genes in the *Arabidopsis* dataset (50.7% BH *q* < 0.05, 39.1% BH *q* < 0.01) using the same criteria (X^2^ (1) = 2727.1, *p* < 0.001, one-tailed, two-proportions z-test). Circadian waveform characteristics of the rhythmically expressed genes (BH *q* < 0.01) in the wheat and *Arabidopsis* datasets were quantified using algorithms in Metacycle (JTK, ARSER, LS and meta2d) and Biodare2 (FFT-NLLS and MESA). Period, phase, and amplitude estimates from FFT-NLLS and meta2d were well-correlated for individual genes (Supplementary Fig. 1). All models reported that mean period length in wheat was approximately 3h longer than in *Arabidopsis* (wheat = 25.9 - 27.5h, *Arabidopsis* = 22.6 - 24.4h; t(36067) = 101.58, *p*<0.001, Welch’s two sample t-test; Supplementary Fig. 2). There was no significant difference between mean periods across the three wheat sub-genomes (Fig. 1c, F(2, 28,276) = 0.179, *p*=0.836, One-way ANOVA). We used meta2d to compare period means and distributions from four previously published circadian datasets, and found that period lengths in wheat were longer and had higher standard deviation than period distributions from *Arabidopsis*, *Brassica rapa*, *Brachypodium distachyon* and *Glycine max* (Fig 1b; Supplementary Table 2).

We investigated how wheat periods changed over the course of the three-day experiment and found that periods were longer immediately after transfer to constant light (28.61h, SD=3.421h), and progressively shortened over the following days (Supplementary Note 2). One explanation for this initial lengthening of period is that the dusk zeitgeber is an important signaling cue for wheat circadian expression. It is possible that the mean period of expressed transcripts initially lengthens trying to follow this missing dark signal before the free-running endogenous period asserts itself.

For both *Arabidopsis* and wheat, we recalculated phases of rhythmic transcripts relative to endogenous period (circadian time; CT). Across all algorithms, most transcripts in *Arabidopsis* peaked during the subjective night (around CT12-24; Supplementary Fig. 3). In wheat, the greatest numbers of rhythmic genes peaked during the subjective day (around CT6-8) with a second, smaller group being expressed in the night (∼CT20). When we grouped transcripts into 2h period bins, we found that transcripts with short periods contained proportionally more dawn-peaking transcripts, whereas those with longer periods contained proportionally more dusk-peaking transcripts (Supplementary Note 3, Supplementary Fig. 4).

### Balance of circadian regulation within triads

Although previous studies have examined the relationships between circadian regulated orthologs in different plant species^26–28^ and within paralogs in *Brassica rapa*^29^, hexaploid wheat provides an opportunity to study the relationships between recently formed circadian regulated homoeologs acting within the same organism. In wheat, over 72% of syntenic triads are estimated to have “balanced” expression, with similar relative abundance of transcripts from each of the three homoeologs^30^. Due to the importance of the clock in coordinating dosage of gene expression, our hypothesis was that many circadian triads would also have balanced circadian regulation. We defined imbalanced circadian regulation as triads harbouring differences in rhythmicity (i.e., BH *q*-values), period lengths, phases, and relative amplitudes.

Of the 16,359 expressed triads in our dataset, 9901 (60.52%) had at least one rhythmic homoeolog, and 3448 (21.08%) had three rhythmically expressed genes (BH *q* < 0.05), with the latter hereafter termed “rhythmic triads” (purple segment, Fig. 1d). 6453 triads lacked rhythmicity in either one or two expressed homoeolog(s) (green and blue segments, Fig. 1d). In both cases, there was no bias for absence of rhythmicity in the A, B or D copy (χ^2^(2)= 6.8415, *p* = 0.40 where one gene is arrhythmic, χ^2^(2)= 6.8415, *p* = 0.03 where two genes are arrhythmic). We found cases where high-confidence rhythmic homoeologs (BH *q* < 0.01) occurred alongside arrhythmic homoeologs (BH *q* > 0.05) represented by light-shaded outer-ring segments in Fig. 1d. In total there were 3450 of these imbalanced-rhythmicity triads (Fig. 1h,i). To explore other forms of circadian imbalance, we assessed whether phase, period and relative amplitude were conserved between homoeologs within the rhythmic triad set (purple segment, Fig. 1d). Differences in phases were quantified by a cross-correlation analysis to assess whether the correlation between homoeologs was improved with a time lag of 4, 8 or 12 hours. We identified 464 triads with imbalanced phases with an optimum lag of >0h between homoeologs (Fig. 1e,j). 1,139 triads had imbalanced periods with more than 2h difference in period between homoeologs (Fig. 1f,k). 701 triads had imbalanced relative amplitudes with more than two-fold difference in relative amplitude (Fig. 1g,l). Within this last group, the homoeolog with the lowest amplitude was still rhythmic, as observed when data are mean-normalized (Fig. 1l,m). In summary, the largest cause of imbalanced circadian expression within triads was absence of rhythmicity (67.89%) with differences in period (22.41%), relative amplitude (13.79%) and phase (9.13%) occurring more infrequently and with some overlap between categories.

Out of all expressed triads in our dataset, around 11.1% had balanced circadian expression, 31.1% had imbalanced circadian expression, 39.5% were arrhythmic and 18.4% were borderline triads which did not fit into the categories imposed by our cut-offs. There is therefore a ratio of approximately 3:1 imbalanced to balanced circadian triads in wheat. This finding was initially surprising given that Ramírez-González et al. reported 72.5% of wheat triads showed balanced expression. We found that 64.15% of the triads classified as circadian imbalanced in our data would be classified as balanced in the Ramírez-González study (Supplementary Fig. 5). However, if we consider that triads with highly imbalanced circadian regulation can be classified as balanced in their expression at a single timepoint (as demonstrated in Supplementary Fig. 6), and that there are multiple ways in which homeologs can become circadian imbalanced (phase, period, rhythmicity etc.), then it is quite reasonable that only a small proportion of triads are classified as having balanced circadian regulation in this study. This insight highlights the importance of considering temporal dynamics when studying gene expression.

One explanation for imbalanced rhythmicity is that arrhythmic homoeologs are silenced. In support of this, we found that the rhythmic homoeologs in imbalanced triads were expressed at a significantly higher baseline level than their arrhythmic homoeologs (Fig. 1n; F(16, 35,148) = 6.94, p<0.001, Two-level, nested ANOVA on Log10 transformed data). We also found that triads with balanced rhythmicity were expressed at a uniformly higher level than the most highly expressed homoeolog(s) in the imbalanced triads (Fig. 1n; F(7, 35,148) = 570.909, p<0.001). Therefore, in imbalanced rhythmicity triads, the rhythmic homoeolog does not appear to compensate for reduced expression of the other homoeolog(s), so the overall expression across the triad is reduced. This is supported by data from diploid *Brassica rapa*, where circadian regulated paralogs are expressed at a higher level than single copy genes^29^.

To investigate whether certain biological processes were associated with circadian balance, we compared GO-slim terms enriched in the 1816 circadian balanced triads, the 5082 differently circadian regulated triads and the 6458 arrhythmic triads, identifying significant terms unique to each group (Table 2). Some terms were enriched only in circadian balanced triads (*p*-value <0.0001, Fisher’s exact test e.g., “photosynthesis”, “generation of precursor metabolites and energy”, “gene-expression” and “translation”). In contrast, GO-slim terms: “developmental process involved in reproduction”, “and “system development” were enriched significantly in triads with differently regulated homoeologs (*p*-value <0.0001) but not in triads with circadian balanced homoeologs (*p*-value >0.5). A possible explanation for this enrichment could be that imbalanced circadian triads are more likely to be dynamically expressed over developmental stages or show local dominance of a sub-genome in a particular tissue type. Transcription factor (TF) triads were as likely to be circadian balanced/imbalanced as non-transcription factors (χ^2^ (2, N = 13,356) = 3.03, *p* = 0.08, chi-square test). Previously validated wheat TFs with imbalanced circadian expression included *WCBF2* (aka *TaCBF1*) and *TaPCF5*, both of which regulate abiotic stress responses^31, 32^ (Supplementary Note 4, Supplementary Fig. 7).

**Table 1:** Numbers of rhythmic genes at (BH *q* < 0.05 or BH *q* < 0.01) in *Arabidopsis* and wheat identified using Metacycle Benjamini Hochberg *q*-values. Periods, relative amplitudes, and *q*-values are estimates from meta2d. Data windows reflect hours relative to transfer to constant light from entrained 12:12h light conditions. A repeat of this table with pre-filtering to remove low-expression genes is provided in Supplementary Figure 1, and the effects on proportions of rhythmic genes are discussed in Supplementary Note 1.

|  | Wheat data from this study |  | *Arabidopsis* data from Romanowski et al. |
| --- | --- | --- | --- |
|  | 24-68 data window | 0-68 data window | 24-68 data window |
| Total number of expressed genes | 86,567 | 86,567 | 26,392 |
| Total rhythmic genes (BH $q < 0.05$) | 28,594 | 28,530 | 13,392 |
| Total rhythmic genes (BH $q < 0.01$) | 18,633 | 21,059 | 10,317 |
| Mean Period (h) (BH $q < 0.05$) | 26.60h (SD 3.62) | 26.75h (SD 2.82) | 23.50 (SD 2.52) |
| Mean Period (h) (BH $q < 0.01$) | 26.82h (SD 3.21) | 26.83h (SD 2.42) | 23.62 (SD 2.04) |
| Mean relative Amplitude (BH $q < 0.05$) | 0.24 (SD 0.19) | 0.26 (SD 0.20) | 0.28 (SD 0.20) |
| Mean relative Amplitude (BH $q < 0.01$) | 0.27 (SD 0.19) | 0.29 (SD 0.21) | 0.30 (SD 0.20) |

**Table 2:**
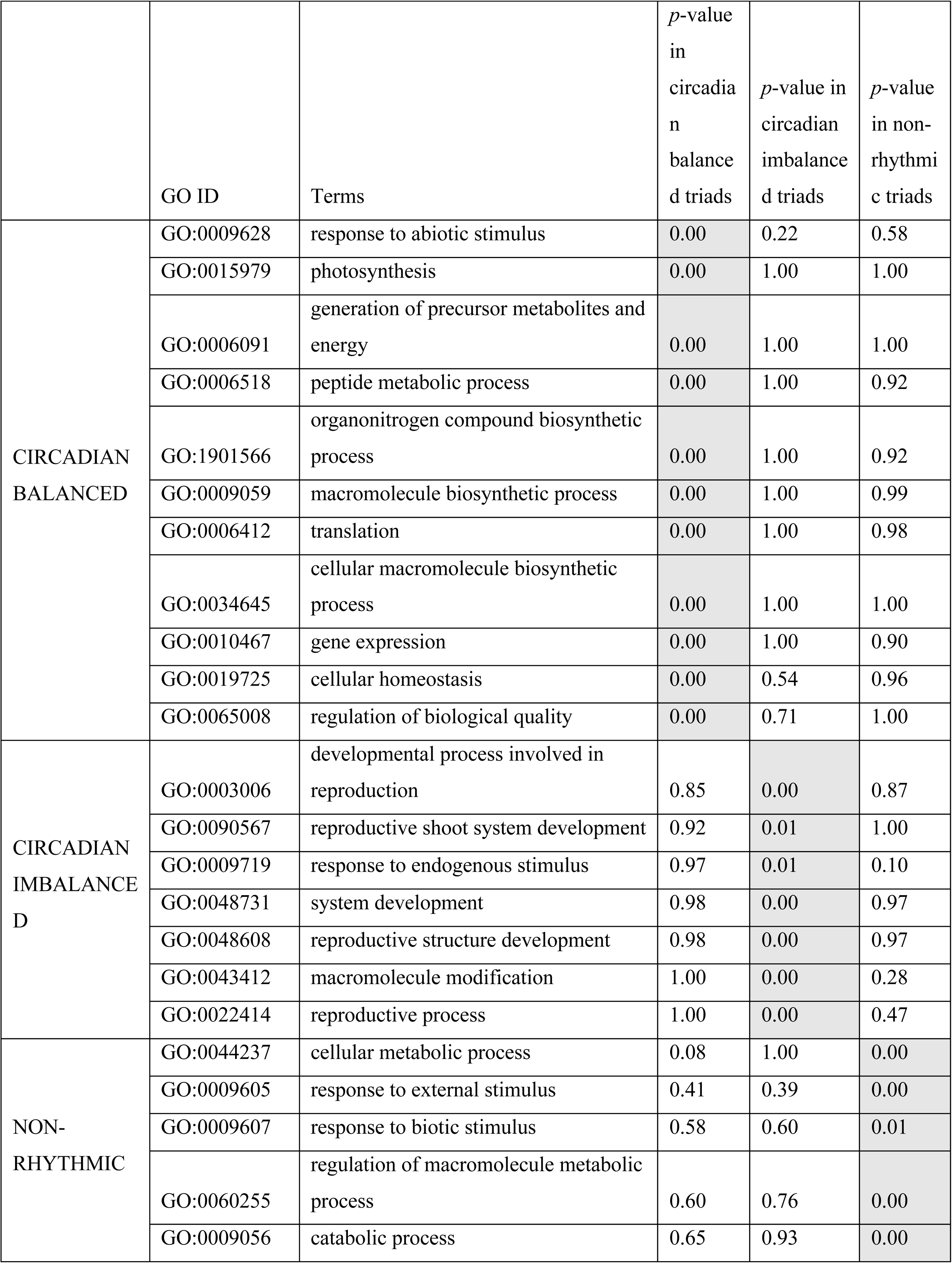
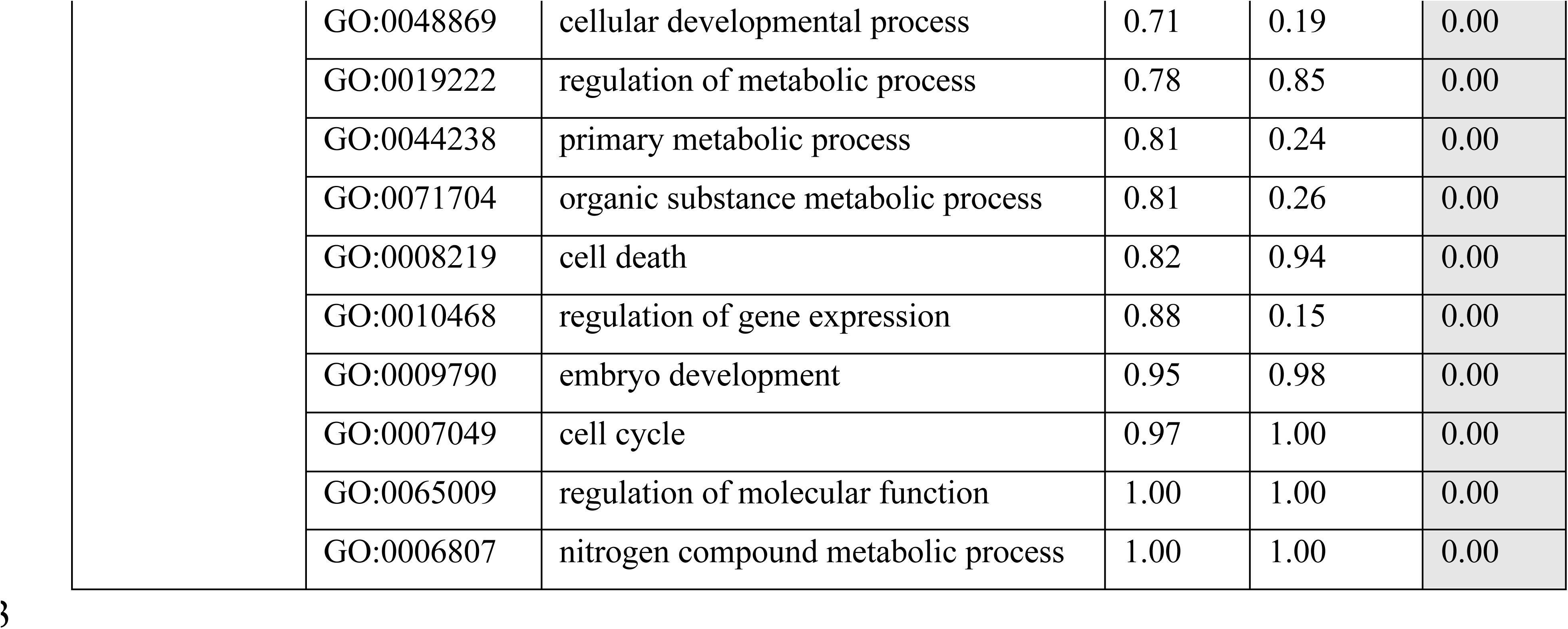
GO-slim terms for biological processes associated with circadian balanced, circadian imbalanced, and arrhythmic wheat triads. Only enriched terms which were highly enriched (Fisher’s exact test *p*<0.01) in one category and non-significantly expressed (*p*>0.05) in other categories is displayed.

When we compared GO-slim terms associated with paralogs in *Brassica rapa*^29^, we found that *Brassica* paralogs with similar circadian expression patterns (as characterized in Greenham et al, (2020) using DiPALM^29^) were also associated with “photosynthesis” and “generation of precursor metabolites and energy” (*p*-value <0.001, Fisher’s exact test). These terms were not enriched in circadian paralogs with differential expression patterns (*p*-value >0.5; Supplementary Table 3). Examples of genes having similar (balanced) circadian expression within *Brassica rapa* paralogs and in all three homoeologs in wheat triads included orthologs of the PSI light harvesting complex genes *LHCA1*, *LHCA2* and *LHCA3*, and the RNA polymerase sigma factor *SIG5.* It is possible that conservation of circadian expression of these duplicate genes poses an evolutionary benefit, as similar regulation has been retained for both duplicate copies in ancient paralogs of *Brassica rapa* and within more recently formed wheat homoeologs arising through polyploidization.

### Patterns of triad circadian balance across the genome

Next, we wanted to determine whether there are genomic regions that incorporate a clustering of circadian balance in rhythmicity. We hypothesized that if we found sequential “runs” of triads with arrhythmic expression on a particular chromosome, this might indicate an overlying region of differential chromatin accessibility or transcriptional suppression. To investigate this, we looked for regions on each set of chromosomes where there were sequential triads having the same number of rhythmic homeologs (i.e. runs of three, two or one rhythmic genes). We also looked for runs of sequential rhythmic triads specific to a particular chromosome (e.g. runs of two rhythmic genes on chromosomes A and B). In both cases, we found no evidence that triads with specific categories of rhythmic homoeologs were grouped together more often than would be expected by chance (Supplementary Note 5, Supplementary Table 4). This suggests that distributions of rhythmic balance appear to be randomly distributed across the genome (Supplementary Fig. 8).

### Clustering of gene expression and GO-term enrichment

To establish whether similar phased transcripts in wheat and *Arabidopsis* had similar biological roles, we clustered rhythmic transcripts (BH *q* < 0.01) into 9 expression modules for each species and identified GO-slim terms enriched in each module (*p* < 0.01). Circadian characteristics of module eigengenes are shown in Supplementary Table 6. We compared the correlation and cross-correlation of pairwise modules in the two species to find modules that correlated with a peak lag of 0 (synchronous phase) or with a peak lag of 4, 8, or 12h (asynchronous phase). Overall, modules with synchronous phases in wheat and *Arabidopsis* shared more GO-slim terms than modules with asynchronous phases (F(3,77) = 4.79, p <0.01, One-Way ANOVA), indicating that these modules in wheat and *Arabidopsis* contain genes with similar functions (Supplementary Fig. 9). We focused on four of these synchronous module pairs, broadly peaking at dawn, midday, dusk, and night for further analysis (Fig. 2). Eigengenes for dawn peaking modules A9 and W9 were highly correlated (r>0.9) and shared 14 overlapping enriched GO-slim terms (*p* < 0.01; Fig. 2a,b,f). These included terms for translation and gene expression as well as terms related to protein, amide, nitrogen and organonitrogen biosynthetic and metabolic processes (full lists in Supplementary Table 7). Co-expressed genes in the dawn-expressed modules included several orthologs involved in light, heat, and biological defence, as well as 45 ribosomal protein orthologs (Supplementary Note 6). Transcripts for ribosomal proteins in mouse liver and *Neurospora crassa* have also been reported to oscillate, suggesting a conserved role for the circadian clock in co-ordinating ribosome biogenesis^33, 34^.

**Fig. 2.**
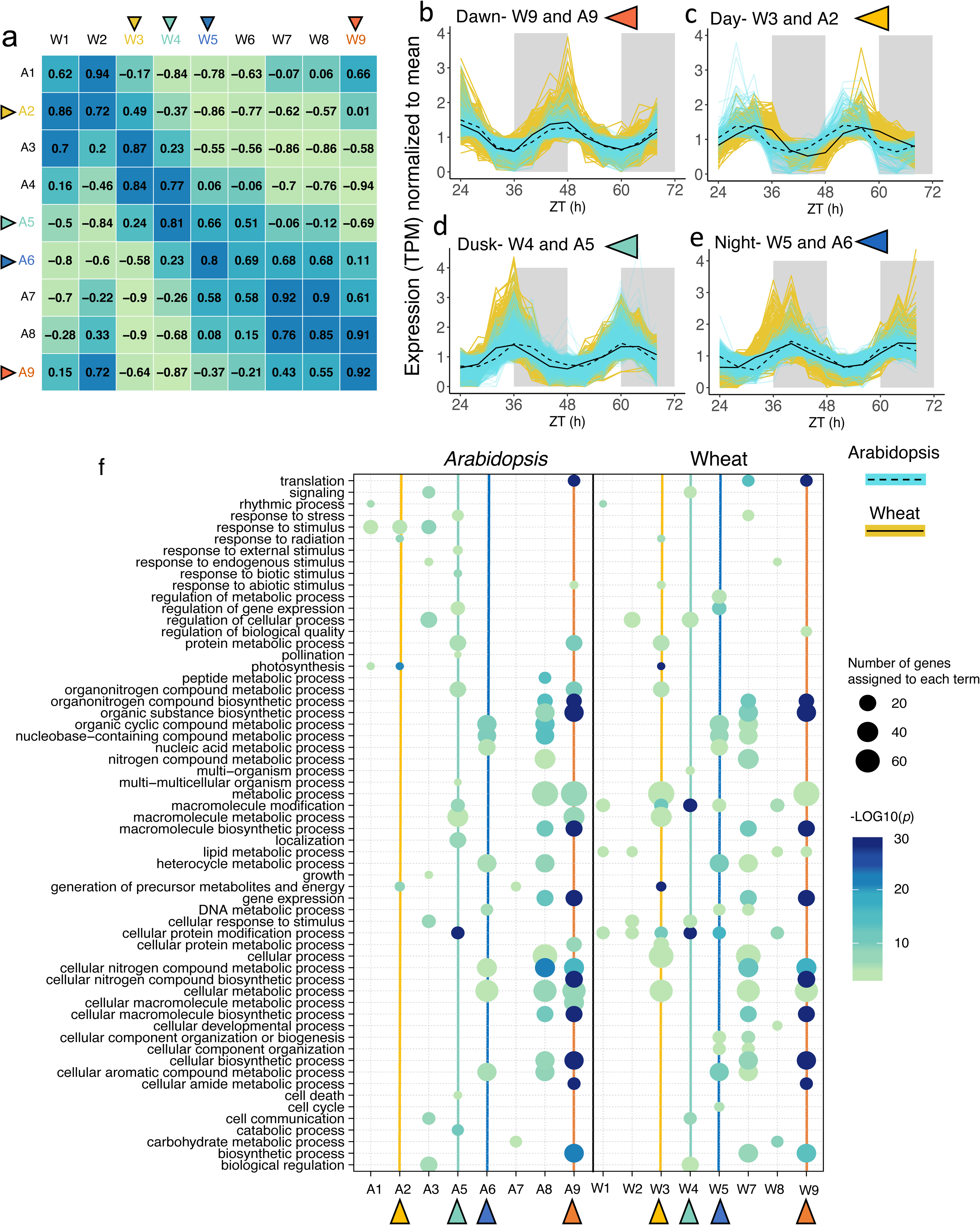
Overlapping GO-slim terms shared between *Arabidopsis* and wheat modules expressed at similar times in the day. **a,** Pearson correlation coefficient (r) between eigengenes for wheat and *Arabidopsis* expression modules ordered by circadian phase. Coloured triangles and axes labels correspond to module expression profiles and columns in bubble-plot. **b-e,** Expression profiles of *Arabidopsis* and wheat modules compared in the main text normalised to their mean. Solid and dashed black lines represent the module eigengene for wheat and *Arabidopsis* modules respectively. **f,** GO-slim terms associated with *Arabidopsis* and Wheat modules. Modules are ordered by predicted CT phase for each species. Only terms with -Log10*p* >3 are shown. Wheat W6 and *Arabidopsis* A4 contained no terms above the significance cut-off and so are not shown. Bubble color indicates the -Log10*p*-value significance from Fisher’s exact test and size indicates the frequency of the GO-slim term in the underlying EBI Gene Ontology Annotation database (larger bubbles indicate more general terms).

In addition to enriched GO-slim terms, we investigated enrichment for transcription factor (TF) superfamilies and transcription factor binding sites in each wheat module of clock-regulated genes. In late-night/dawn modules W8 and W9, transcripts encoding MYB transcription factors were significantly enriched and included putative TFS involved in leaf morphogenesis, plant growth, regulation of flavonoid biosynthesis, and developmental transition to flowering (Supplementary Note 6, Supplementary Fig. 10).

Eigengenes for wheat and *Arabidopsis* modules peaking in the day (W3 and A2) had a relatively low correlation (*r*=0.491), but peaked with similar CT values (CT 6.34h and 6.19h) given the longer circadian period in wheat. 5 out of 15 of the GO-slim terms enriched in the W3 module were also found in the A2 module (*p* < 0.01; yellow triangles in Fig. 2a,c,f). These included terms relating to “photosynthesis”, “response to radiation” and “generation of precursor metabolites and energy”. Co-expressed genes peaking in day-time modules included light-harvesting and light signaling genes as well as *CYP709B3,* which protects the plant from transpiration-triggered salinity stress during the day^35, 36^. In dusk-peaking modules A5 and W4, 8 significantly enriched GO-slim terms were shared between *Arabidopsis* and wheat (*p* < 0.01, green triangles in Fig. 2a,d,f). Several genes co-expressed in these dusk modules were involved in auxin transport and signalling including the endosomal sorting complex protein *CHMP1A* which ensures proper sorting of auxin carriers (Supplementary Note 6)^37^. There was also a significant enrichment for expression of transcripts encoding AP2-EREBP (ethylene responsive) and ARF (auxin responsive) transcription factor superfamilies within the W4 module (Supplementary Fig. 10). Interestingly, this was followed two hours later by the expression of genes with AP2-EREBP transcription factor binding sites in their promoter region (W5, Supplementary Fig 11). Finally, two evening-phased modules W5 and A6 (*r*=0.80) were enriched for GO-slim terms concerning several metabolic processes (blue triangles in Fig. 2f). Co-expressed orthologs in these two modules included *SEVEN IN ABSENTIA2* that regulates ABA-mediated stomatal closure and drought tolerance in *Arabidopsis*^38^, and *HYDROPEROXIDE LYASE1* that contributes to responses to insect attack and mechanical wounding^39^.

### Components of the core circadian clock in *Arabidopsis* and wheat

We next compared the dynamics of circadian oscillator components in wheat and *Arabidopsis*. Clock gene orthologs belonging to large gene families were detected by phylogenetic analysis (Supplementary Fig. 12–16; Supplementary Table 8). Overall, wheat circadian clock genes were expressed rhythmically and with a similar phase to their *Arabidopsis* counterparts (Fig. 3). However, the free-running rhythms of clock transcripts in wheat had a mean circadian period that was approximately 3.49h longer than in *Arabidopsis* (27.23h and 23.74h, respectively).

**Fig. 3.**
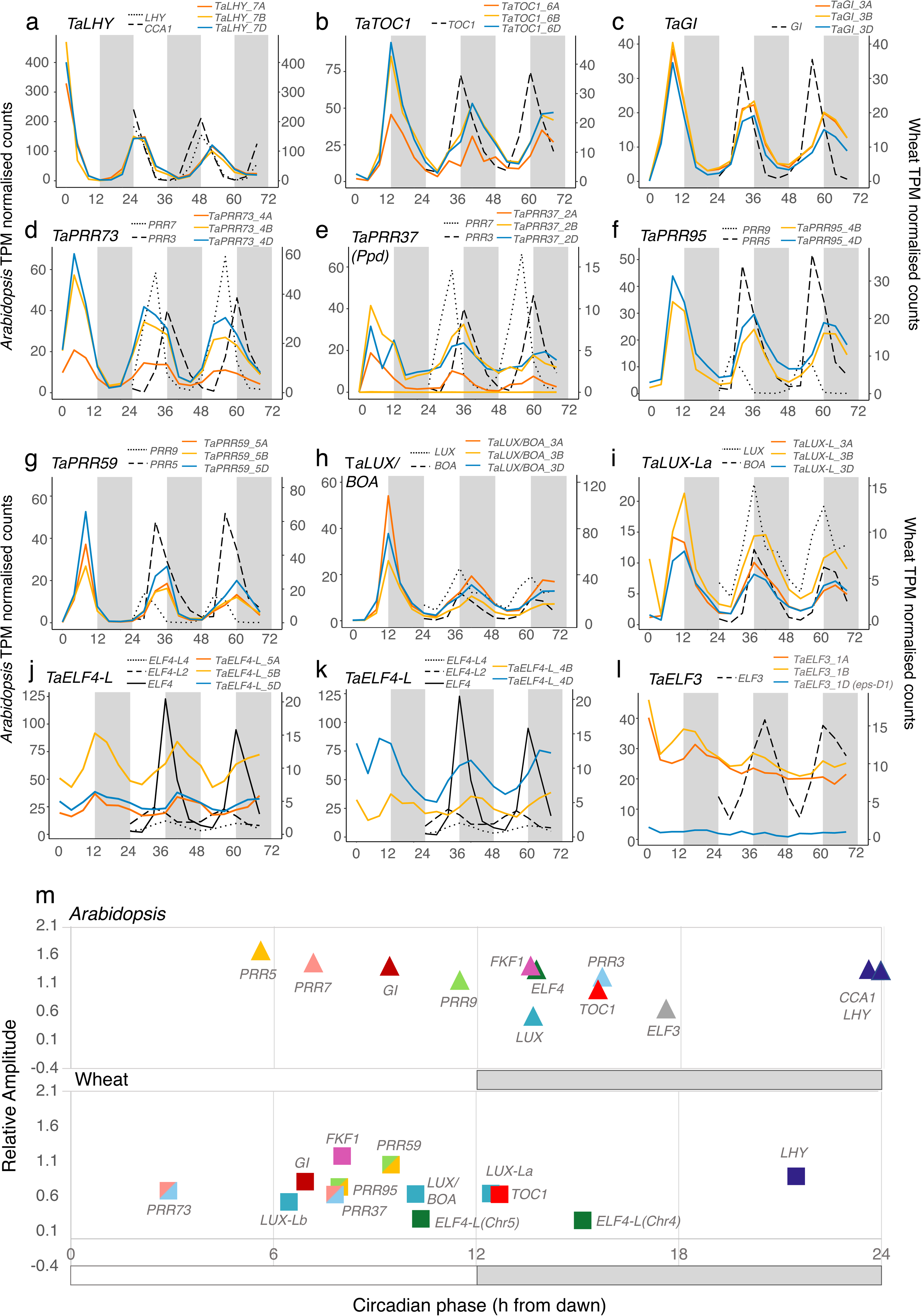
Free-running expression of core circadian clock genes in wheat and their homologs in *Arabidopsis.* **a-l,** Wheat circadian clock genes were identified through alignment of phylogenetic protein family trees or BLASTP to known clock gene homologs. Gene IDs for each gene set are in Supplementary Table 8. Wheat homoeologs are coloured according to their identity to either the A genome (orange), B genome (yellow) or D genome (blue) and grey and white blocks indicate subjective dark and light time periods under constant conditions. Data represent the mean of three biological replicates and transcript expression is collapsed to gene level. Expression profiles for additional core circadian clock genes are in Supplementary Fig. 17. **m,** phases of core clock genes in *Arabidopsis* and wheat (meta2d estimates from data 24-68h after transfer to L:L). Genes were not plotted if B.H *q*-values were > 0.01. Wheat values represent circular mean circadian phases (CT) across homoeologs calculated in Supplementary Table 9.

*TaLHY* and *TaTOC1* peaked sharply at dawn and dusk, respectively, during the first cycle in constant light, and maintained an >8h difference in phase throughout the experiment (Fig. 3a,b). This is consistent with their phasing in *Arabidopsis.* All three homoeologs for *TaGI* were robustly rhythmic (BH *q* < 0.01) and peaked at CT7 (Fig. 3c). *TaPRR73* transcripts peaked approximately 5h before *TaPRR37* transcripts, consistent with the phase divergence of *Arabidopsis PRR7* and *PRR3* (Fig. 3d,e). However, wheat homoeologs *TaPRR59* and *TaPRR95* had similar expression profiles (R^2^=0.68) peaking marginally apart at CT8 and CT10, in between the peak phases of *Arabidopsis PRR9* (CT5) and *PRR5* (CT11) (Fig. 3f,g). Therefore, the *PRR* gene family in wheat peaks in the order of; *TaPRR73*, [*TaPRR37*, *TaPRR95*, *TaPRR59]* in quick succession, and finally *TaTOC1.* This sequential pattern matches the expression of PRR homologs in rice^40^.

Transcripts encoding evening complex components, *LUX, ELF3* and *ELF4,* are circadian regulated in *Arabidopsis* and peak simultaneously at dusk. Three wheat triads for LUX-like genes were identified; one with higher identity to *LUX/BOA* and two similar to other *LUX-* like *Arabidopsis* genes (Supplementary Fig. 15). Transcripts from all three of these triads accumulated rhythmically and peaked from midday to dusk, *TaLUX-Lb* at CT7, *TaLUX/BOA* at CT10 and *TaLUX-La* at CT12 (Fig. 3h,i; Supplementary Fig. 17). Five wheat transcripts with homology to *Arabidopsis ELF4* and *ELF4-L1-4* accumulated with a mean circadian phase of 12.6h; similar to *ELF4,* but with lower relative amplitudes (Fig. 3j,k; Supplementary Fig. 17). *TaELF3* transcripts were all arrhythmic (BH *q* > 0.36), and the *TaELF3-1D* homoeolog was expressed with a particularly low baseline level of 0.73 TPM in comparison to the other two homoeologs (8.7-9.7 TPM) (Fig. 3l). This is consistent with previous findings linking a deletion in *TaELF3* to the *eps* QTL on chromosome 1D in Cadenza^18^. We next assessed the balance of circadian expression between triad homoeologs in the core clock network. *TaLHY, TaGI*, and *TaPRR59* had notably similar expression patterns in terms of phase, period, relative amplitudes, and baseline expression over all timepoints (Fig. 3a,c,f,g), suggesting that unbalance in these triads is strongly selected against. Homoeologs of *TaTOC1*, *TaPRR73* and *TaPRR37(Ppd)* had similar phases, but all had reduced expression in the A-genome homoeolog (Fig. 3b,d,e). *TaLUX/BOA-3A*, *TaLUX-La-3B* and *TaLUX-Lb-1D* had marginally shorter periods (>2h) and delayed phases (>2h) compared to their respective homoeologs (Supplementary Table 9).

The REVEILLE family are CCA1/LHY-like MYB-domain transcription factors that are predominantly activators of evening expressed genes^41, 42^. The wheat *RVE* genes could be split into a *LHY* clade (containing the *TaLHY* triad described above), a *RVE6/8*-like clade containing three wheat triads and a *RVE1/2/7*-like clade also containing three triads (Supplementary Fig. 12). All *TaRVE6/8* transcripts peaked at CT0-4 concurrently with *TaLHY* (Supplementary Fig. 17). The *TaRVE2/7* transcripts peaked with distinct phases; *TaRVE27b* in phase with *TaLHY*, *TaRVE27c* 4h before *TaLHY* and *TaRVE27a* approximately 12h before *TaLHY* (Supplementary Fig. 17). Based on their phylogenetic relationships, it is probable that several *RVE2/7* clade paralogs in wheat and *Arabidopsis* arose independently after their evolutionary divergence, and it is therefore interesting that they both show distinct phases of expression, suggesting homoplastic circadian functions. Expression of orthologs for additional transcripts involved in circadian regulation (*FKF1*, *ZTL*, *LKP2*, *LNK1/2*, *CHE* and *LWD*) are reviewed in Supplementary Note 7 and Supplementary Fig. 17.

### Circadian control of photosystem and light signalling gene expression is largely conserved between *Arabidopsis* and wheat

A further GO-slim analysis across all rhythmically-expressed genes in *Arabidopsis* and wheat identified enrichment of similar GO-slim processes including “photosynthesis” (p<1x10-14), “rhythmic process” (p<1x10-6), “response to abiotic stimulus” (p<1x10-13) and “cellular macromolecule biosynthetic process” (p<1x10-5, Fisher’s exact test, Supplementary Table 10). We used genes associated with some of these GO-slim terms as case-studies to highlight similarities and differences in circadian control between the two species. Expression data and Metacycle statistics for all transcripts in this analysis are in Supplementary Table 11.

In considering photosynthesis, we examined specifically nuclear genome-encoded photosystem (PS) proteins. Transcripts encoding the PSI components *LHCA1-6,* the PSI reaction centre subunits *PSAD* and *PSAE* and the *PSII* subunits *LHCB1-7* were rhythmically expressed in both species and had conserved phases (Supplementary Fig. 18). *LHCA1-4* peaked towards the end of the subjective day and *LHCA5* and *6* peaked during the subjective night. *PSAD* and *PSAE* peaked concurrently with *LHCA1-4*. In both species, *LHCB7* transcripts had lower relative amplitudes compared to other LHCB transcripts. PSB27 is a protein associated transiently with the PSII complex involved in adaption to fluctuating light intensities^43^. Transcripts for this protein peaked during the subjective day in *Arabidopsis* and during the subjective night in wheat.

In considering the GO-slim term “response to abiotic stimulus”, we next investigated expression of transcripts for photoreceptors and light signalling proteins due to their pervasive influence upon development, metabolism, and circadian timing. Although transcripts for the UV-B photoreceptor *UVR8* accumulated with a circadian rhythm in both species, only one PHYTOCHROME ortholog (*PHYA)* and three *CRYPTOCHROME* orthologs *(CRY1)* were rhythmic in wheat out of 18 orthologs identified (Supplementary Fig. 18). This contrasts with *Arabidopsis*, where *PHYA-C, CRY1* and *CRY2* accumulated with a circadian rhythm.

Downstream light signalling proteins COP1 and SPA form complexes that degrade positive regulators of photomorphogenesis (e.g. *HFR1* and *HY5*) under dark conditions^44^. Transcripts for *COP1*, *SPA4*, *HFR1* and *HY5* accumulated rhythmically and with conserved phases in both species (Supplementary Fig. 19). *COP1/SPA4* peaked synchronously around the end of the subjective night. Surprisingly, given the similar role HFR1 and HY5 proteins have in preventing hypocotyl elongation in low light, *HFR1* and *HY5* transcripts were expressed anti-phase to each other. HY5 and HFR1 act synergistically to coordinate the photomorphogenesis response, although it has been suggested that their activation is regulated through independent pathways^45^.

Wheat triads with identity to *Arabidopsis PIN1*, *PIN4, PIN5,* and *PIF4/5* were rhythmically expressed, alongside two triads with high similarity to rice *OsPIL11* and *OsPIL13* (^46^; Supplementary Fig. 19). Overall, we observe that the arrhythmic accumulation of most of the wheat PHY and CRY transcripts is not reflected in the rhythmic expression of several downstream light signalling transcripts. This supports the notion that regulatory signals from photoreceptors might occur at the level of protein stability and localisation rather than at the level of transcript accumulation, as occurs for *ZTL* or *HY5* in *Arabidopsis*.

A set of proteins that link light signalling, circadian regulation and chloroplasts are the sigma factors^47^. These light-responsive nuclear-encoded regulators of chloroplast transcription guide promoter recognition and transcription initiation by plastid encoded RNA-polymerase (PEP) on the chloroplast genome^48–51^. In *Arabidopsis*, *SIG1, 3, 4*, *5* and *6* were rhythmically transcribed (Supplementary Fig. 19). In wheat, all homoeologs in triads orthologous to *SIG1*, *SIG3* and *SIG5* were also rhythmic (BH *q* < 0.01). Whilst the dawn phase of *TaSIG5* transcripts were similar to *AtSIG5*, *TaSIG1* transcripts were expressed over 10h earlier than *AtSIG1* (Supplementary Fig. 19). Previous research has shown that activity of *AtSIG1* can be regulated through redox-dependent phosphorylation^52^, and activity of all sigma factors are likely to be subject to multiple layers of regulation in addition to circadian control of transcript expression.

### Similarities and differences in circadian control of primary metabolism genes in *Arabidopsis* and wheat

Expression profiles of genes with key roles in primary metabolism were compared in *Arabidopsis* and wheat with a focus on enzymes that regulate trehalose 6 phosphate (Tre6P) and starch metabolism (Fig. 4). Tre6P synthase (TPS) and Tre6P phosphatase (TPP) participate in the synthesis and dephosphorylation of Tre6P, respectively. Tre6P is an important signalling metabolite associated with both sucrose regulation and circadian regulation in *Arabidopsis*^53–55^. Tre6P also affects grain yield and drought resilience in wheat, maize, and rice^56^. Transcripts for *TPS1*, *2*, *6*, *8*, *9*, *10* and *11* and *TPPA*, *E*, *F*, *G* and *H* were expressed rhythmically in *Arabidopsis* (Supplementary Fig. 20). Wheat transcripts for *TPS1* (the most well-characterised of the T6P synthases) were arrhythmic, however rhythmic transcripts were found in triads more closely related to TPS11, 6 and 7 (Supplementary Fig. 20). We identified three rhythmic TPP triads in wheat, two of which were orthologous to *Arabidopsis TPPA*, *F* and *G.* The third TPP triad was part of a monocot-specific clade identified by Paul et al. (2018), which also included *Zm00001d032298,* a crop improvement target in maize^56^.

**Fig. 4.**
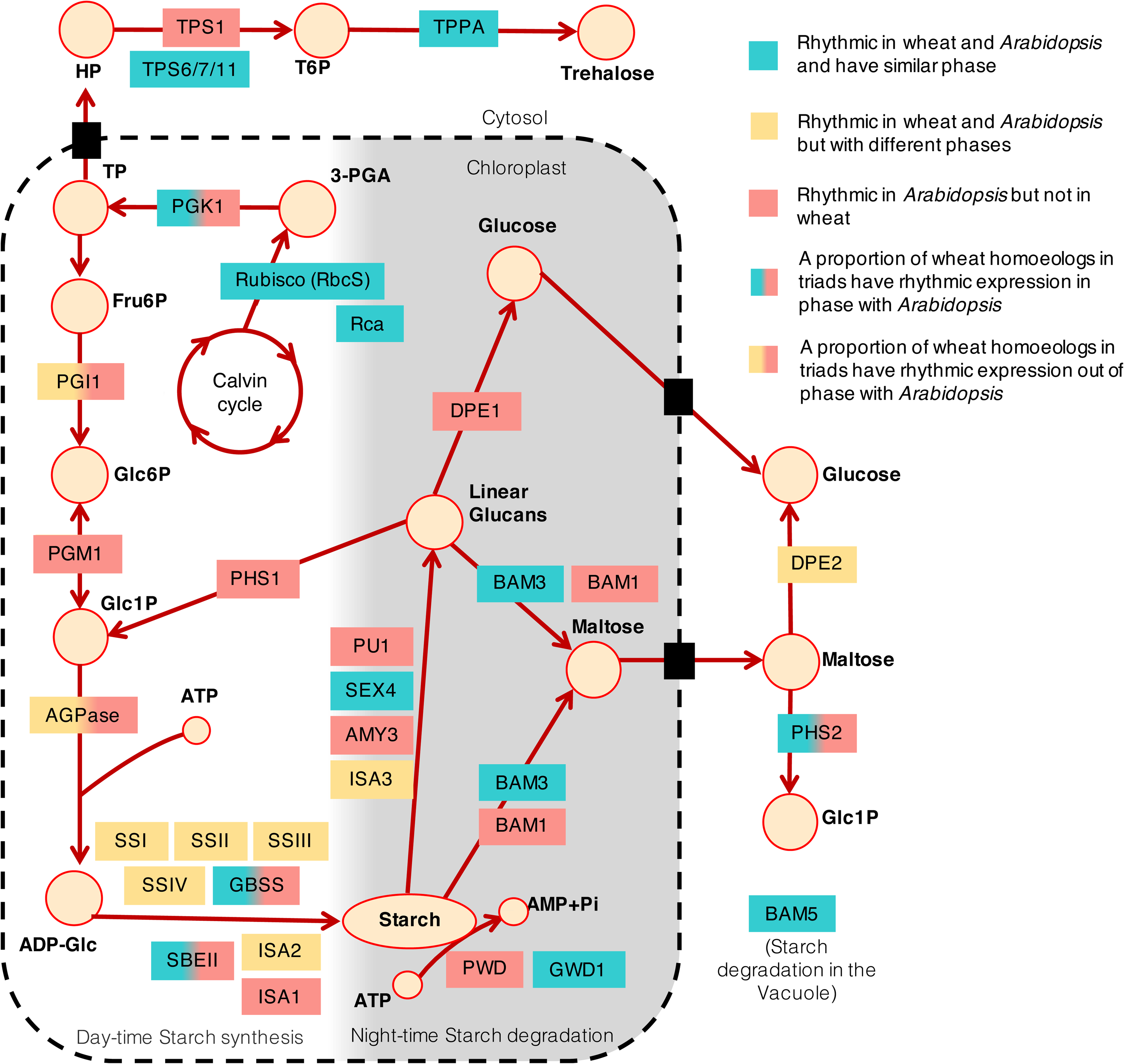
Similarities and differences in circadian control of transcript accumulation in key genes involved in primary metabolism and signalling. Circles represent metabolites involved in the breakdown and biosynthesis of starch. Starch synthesis occurs during the day and breakdown occurs at night as indicated by the yellow to grey shading gradient. The dotted line encloses processes which take place in the chloroplast. Abbreviations: HP: Hexose-phosphate, T6P: Trehalose-6-phosphate, TP: Triose phosphate, 3-PGA: Glycerate 3-phosphate, Fru6P: Fructose-6-phosphate, Glc6P: Glucose-6-phosphate, Glc1P: Glucose-1-phosphate, ATP: Adensoine tri phosphate, ADP-Glc: ADP-glucose, TPS: Trehalose phosphate synthase, TPP: Trehalose phosphate phosphatase, PGK1: Phosphoglycerate kinase 1, PGI1: Glucose-6-phosphate isomerase, PGM1: Phosphoglucomutase-1, PHS1 and 2: ALPHA-GLUCAN PHOSPHORYLASE 1 and 2, AGPase: ADP-Glc pyrophosphorylase, BAM1,3,5: β-amylase 1,3,5, ISA1,2,3: Isoamylase 1,2,3, DPE1,2: Disproportionating enzyme1 and 2, SBEI,II: Starch branching enzyme I, II, PU1: Pullulanase 1, PWD: Phosphoglucan, water dikinase, GWD: α-glucan, water dikinase, SEX4: starch excess 4, AMY3: α-amylase, GBSS: Granule bound Starch synthase, SSI-IV: Starch synthase I-IV. Pathway references:^109–111^.

Ribulose bisphosphate carboxylase (Rubisco) comprises eight small (RbcS) and 8 large (RbcL) subunits, which are encoded by the nuclear and chloroplast genomes, respectively^57^. Rubisco requires activation by Rubisco activase (RCA) to release its activity from inhibitory substrates^58^. In our wheat expression data, 22 putative wheat orthologs for the small subunit of Rubisco were rhythmic, peaking during the subjective night, as *RBCS1A*, *RBCS1B*, *RBCS2B* and *RBCS3B* do in *Arabidopsis* (Supplementary Fig. 20). Two triads with identity to Rubisco activase were identified, one of which accumulated rhythmically (peaking at CT0, as with *Arabidopsis RCA*).

Circadian regulation has a pervasive influence on starch metabolism in *Arabidopsis*, particularly the nocturnal rate of transitory starch degradation^23, 59^. Chloroplast phospho-glucose isomerase 1 (*PGI1*) and chloroplast phosphoglucose mutase (*PGM1*) are essential enzymes that link the Calvin-Benson cycle with starch biosynthetic pathway^60–62^. In *Arabidopsis,* these transcripts accumulated with a circadian rhythm (BH *q* < 1x10^-4^); *PGM1* peaked just after dusk (CT14), and *PGI1* slightly later at CT20. In contrast, only one wheat *TaPGI1* homoeolog was rhythmic (BH *q* < 0.01), which had a low relative amplitude (0.16) and a peak phase of CT8. No homoeologs for *TaPGM1* were rhythmically expressed (BH q > 0.01, Supplementary Fig. 20).

ADP-glucose pyrophosphorylase (AGPase) mediates the first irreversible and rate-limiting step in starch biosynthesis through the formation of ADP-Glc. In *Arabidopsis,* transcripts encoding the small and large subunits of AGPase (*APL1*, *APL2*, *APL3*, *APS1*) were rhythmic, peaking at night around CT20. In comparison, in wheat only two of the eleven transcripts with homology to *APL1*, *APL2* and *APS1* were rhythmic (BH *q* < 0.01), with the remaining transcripts lacking a discernible rhythm (BH *q* > 0.05) (Supplementary Fig. 20).

Starch synthases (SS) represent another group of metabolically important enzymes that use the glucose from ADP-Glc to elongate glucan chains. In *Arabidopsis*, there are five types: SSI, SSII, SSIII, SSIV and granule bound GBSSI. SSI-IV are responsible for synthesis of amylopectin, with SSIII and IV determining starch granule number and morphology^63^. *GBSSI* is a known dawn-expressed gene, regulated directly by CCA1/LHY, specialised for amylose synthesis^64^. In wheat, *GBSSI* orthologs are called *TaWaxy* and cultivars with three null alleles produce amylose-free starch in their grain^65^. Comparison of starch synthase expression in *Arabidopsis* and wheat revealed several differences between the phases and relative amplitudes of these transcripts (Supplementary Fig. 20). In *Arabidopsis, GBSSI* transcripts had by far the greatest relative amplitude (1.26) with peak expression at dawn. The next greatest amplitudes were of *SSIV* transcripts, which peaked at CT17. *SSII* and *SSIII* peaked together at CT21 and *SS1* peaked at CT8 with a much smaller amplitude (0.12). In contrast, in wheat, an *SSIII* triad (*TaSSIIIb*) had the largest relative amplitude rhythms of the wheat starch synthases identified (0.64 - 0.73). Wheat transcripts for *SSI* and *SSIV* also peaked in the morning, whereas wheat *SSII* peaked instead in the subjective night (∼CT15). In our data, TaWaxy (GBSSI) transcripts were present at a very low baseline level (<0.01 TPM) and without any circadian oscillation. However, another wheat triad, *TaGBSSII,* shared >62% identity with *Arabidopsis GBSSI,* and the B and D homoeologs had rhythmic expression which peaked at dawn. TaWaxy and *TaGBSSII* are specific to endosperm and leaf tissues, respectively^66^, which might explain the distribution of transcript accumulation seen here. We can conclude that the circadian clock regulates the expression of SS transcripts in both *Arabidopsis* and wheat, although there might be an emphasis on different types of SS in each species.

The *Arabidopsis* circadian clock regulates the rate of starch degradation so that starch reserves are depleted precisely at subjective dawn^23^. Many transcripts encoding starch-degrading enzymes in *Arabidopsis* had synchronized dusk peaks: Isoamylase-type starch debranching enzyme *ISA3*; alpha-amylase *AMY3*; plastidial phosphorylase *PHS1-2*; disproportionating enzymes *DPE1-2*; glucan, water dikinases *GWD1* and *PWD* and glucan phosphatase *SEX4. Arabidopsis* transcripts for *BAM3*, *BAM5* and *PU1* also oscillated with a circadian rhythm, peaking later in the subjective night. Strikingly, wheat orthologs for several of these genes were not rhythmic, including *AMY3*, *DPE1*, *PWD, PHS1, PU1* and *BAM1*. Wheat orthologs for *ISA3* and *DPE2* were expressed rhythmically, but peaked approximately 8-12 h ahead of their *Arabidopsis* counterparts. Some starch degradation enzymes had conserved circadian expression patterns in the two species, such as *SEX4*, *GWD1, BAM3* and *BAM5* transcripts. GWD catalyses glucan phosphorylation and *SEX4* encodes a phosphoglucan phosphatase, both of which facilitate hydrolytic attack by β-amylases (BAM) in the early steps of starch degradation^59, 67^.

## Discussion

### Conservation of circadian regulation between homoeologous genes

We identified a large proportion of imbalanced circadian triads in our dataset. It was our initial expectation that there would be strict balance between the majority of circadian regulated homoeologs due to the critical and finely balanced role the clock has in regulating the transcriptome, and due to the reported high levels of balance reported from single-timepoint transcriptomic analysis in wheat^30^. Instead, we find three times as many triads with imbalanced circadian rhythms as triads with balanced circadian rhythms. This is likely to be partly due to our multiple classification of circadian imbalance as any triad with different rhythmicity, period, phase or relative amplitudes between homeologs. Another factor which distinguishes ratios of transcriptional balance (as defined by Ramírez-González et al. (2018)) and circadian balance is that transcriptional balance within circadian triads is often dynamic across a time-course, shifting between balanced, dominant and supressed relationships over time (Supplementary Fig 6).

Most of these imbalanced circadian triads were imbalanced due to arrhythmicity in one or two homoeologs expressed at a lower mean level than the rhythmic homeologs. The reduction of expression could be due to constitutive epigenetic silencing or changes to promoter regions, allowing differential binding of transcription factors^68–70^. These are likely to be triads where one or two homeologs take responsibility for performing the biological function of the triad, and the other homoeolog has reduced functionality. We found additional circadian unbalance in the form of altered phase, period, and relative amplitudes. It is possible that some of these differences are due to retention of circadian regulation from the ancestral genome of each homeolog (Fig. 1a), although it is likely that other differences reflect more recent diversification in expression as a step towards neo-functionalisation. It has been previously suggested that functional divergence is a likely fate for duplicated genes in a sufficiently large population^71^. In *B. rapa,* 42% of circadian controlled paralogs had differential expression patterns^29^, however these paralogs arose through whole genome duplication events around 13-43 million years ago, so have been exposed to longer periods of time during which selection could act upon these duplicate genes^72^. In comparison, specialisation of circadian homeologs in wheat could be comparatively lower due to the relative infancy of its polyploidisation around 10,000 years ago.

### Differences between periods of rhythmic transcripts in *Arabidopsis* and wheat

The mean period of circadian regulated genes in wheat was over three hours longer than in *Arabidopsis*. Period length is affected by a range of exogenous conditions (e.g. light and temperature), and varies between tissues and plant age^73^. There is also evidence that longer periods have been selected for during cultivation of crops at higher latitudes^1, 2, 74^, potentially due to enhanced seasonal tracking capability enabling precision timing of growth and flowering^75^. Compared to other plant circadian transcriptome data sets, rhythmic wheat transcripts also had higher period variance (Fig 1b). The broad period distribution in wheat might be due to inclusion of all aerial material in our sampling strategy. Variation in free-running periods could occur at the organ-, tissue- or cellular-level, and transcripts which are highly expressed in those regions may reflect those period differences^76, 77^. An alternative possibility is that period variation is due to uncoupling of multiple circadian oscillators within the same cell which control expression of subsets of transcripts^78–80^. Future research could examine the relationship between period distributions of circadian transcriptomes and the effects of domestication, latitudinal adaption, monocot-dicot divergence, or polyploidy.

### Similarities and differences in circadian regulation between wheat and *Arabidopsis*

Our analysis revealed extensive conservation of time-of-day specific GO-slim processes and co-expressed genes between *Arabidopsis* and wheat. These included genes involved in photosynthesis (e.g., photosystem proteins), light signalling (e.g., *HFR1, HY5, PINs* etc), translation (e.g., ribosome proteins) and auxin and ethylene responsive transcription factors. The striking conservation of photosynthesis related genes was also reflected by the enrichment of these genes in both balanced wheat triads and similarly expressed circadian paralogs in *Brassica napus*. Photosynthetic outputs have also been reported to be governed by the circadian clock in the liverwort species *Marchantia polymorpha*, in diazotrophic cyanobacterium *Cyanothece* sp. and in alga *Aegagropila linnaei*, perhaps suggesting a widespread control mechanism with an ancient evolutionary origin^81–83^. We also identified several interesting differences between *Arabidopsis* and wheat, including absence of rhythmicity in wheat *PHY* and *CRY* transcripts and antiphase expression of the wheat sigma factor *SIG1.* Furthermore, we found differences in rhythmic expression of many transcripts involved in regulating Tre6P and starch metabolism.

In our data, putative wheat homeologs of *TPS1* were arrhythmic. Instead, rhythmic *TPS* transcripts in wheat had similarity to *Arabidopsis TPS11*, *6* and *7* (Supplementary Fig. 20). In *Arabidopsis*, TPS1 is the most catalytically active and best characterised TPS, and feeds back into the entrainment of the circadian clock^54, 84^. If the lack of rhythmicity in wheat *TPS1* transcripts is reflected at the level of protein activity, it may indicate that Tre6P synthesis is not regulated as tightly by the circadian clock in wheat as in *Arabidopsis*. On the other hand, circadian control of other TPS triads may have implications for biotic or abiotic defence in wheat. TPS5-11 have been previously implicated in control of stomatal aperture^85^, thermotolerance^86^, and defence against fungal, bacterial and aphid attack^87, 88^. In rice, *OsTPS8* influences drought resistance through suberin deposition^89^, and wheat *TaTPS11* participates in a cold stress response^90^.

In wheat, transcripts for starch degradation enzymes (*PHS1, DPE1, BAM1, PU1, AMY3, PWD*) and starch biosynthesis enzymes (*PGI1*, *PGM1*, *ISA1* and *ATPase*) had either arrhythmic expression or low relative amplitudes compared with the robust rhythms of many of these transcripts in *Arabidopsis.* Additionally, *ISA2*, *ISA3* and the starch synthases (*SSI-IV*) had differing circadian phases between the two species. While it is possible that rhythmic expression of a reduced number of genes (e.g.: *SEX4*, *GWD1, BAM3* and *BAM5)* is sufficient to mediate circadian control of starch degradation in wheat, these data suggest that the circadian clock has a less pervasive influence upon transcriptional control of starch metabolism in wheat compared to *Arabidopsis* (Supplementary Note 8).

## Conclusions

Our data reveal the influence of circadian regulation on the wheat transcriptome and highlight several intriguing differences between rhythmically expressed transcripts in *Arabidopsis* and wheat. It explores the added complexity of co-ordinating circadian expression across multiple sub-genomes in a hexaploid species. Given the circadian clock has been under selection during domestication and presents multiple targets for crop improvement, it is likely that this new insight into the clock in wheat will be important in the development of new sustainable and resilient cultivars. It is our hope that these data provide a resource for identifying target genes regulated by the circadian clock, allowing the relationships between chronotype, yield and resilience to be explored in future studies.

## Methods

### Plant materials and growth conditions

Wheat: Wheat seeds of the spring wheat cv. Cadenza were imbibed for three days on damp filter paper on a Petri dish at 4°C. Plates were moved at dawn (06.00 = ZT0), to a growth cabinet set to 22°C under 12:12 light: dark cycles (approximately 200 µmol m−2 s−1). After two days, only seedlings with fully emerged radicles were sown, 3 cm deep in Petersfield cereal mix in 9cm pots. Plants were not vernalized. Seedlings were grown under 12hlight:12hdark conditions for 14 days. After 14 days, at dawn (ZT0) seedlings were transferred to constant light conditions, tissue was sampled every 4h for 3 days (18 samples in total). At each timepoint, we sampled the entire aerial tissue from 3 replicate plants, which was frozen immediately in liquid nitrogen before storage at -80°C. Total RNA was extracted using Qiagen RNeasy plant mini kits (cat. no. 74904) with on-column DNAse treatment (RNAse-Free DNase Set (cat. no. 79254). RNA concentration and integrity were quantified using a Nanodrop Spectrophotometer and Perkin Elmer LabChip GX Nucleic acid analyser before sequencing.

Details of growth conditions for Arabidopsis^26^, Brassica rapa^29^, Brachypodium distachyon^27^ and Glycine max^28^ datasets can be viewed in their source manuscripts. Briefly, all circadian data were measured under constant light and temperature following 12h:12h light:dark entrainment other than Glycine max^28^ which was entrained under 16h:8h light:dark cycles.

### Wheat mRNA sequencing, read alignment and quantification

Library preparation was carried out following the Illumina TruSeq protocol and reads were sequenced on a NovaSeq S2 flow cell at the Earlham Institute. 150bp paired-end reads were generated from each library to an average depth of 84M reads per replicate. Reads were filtered for quality and any remaining adaptor sequence was trimmed with Trimmomatic^91^. Surviving reads were aligned to the Chinese Spring RefSeq v1.1 wheat genome^11^ using HISAT2^92^ with default parameters. Uniquely mapping reads were then quantified using StringTie^93^ and TPM values were extracted for each gene per sample.

### Processing and quantification of previously published datasets

Raw reads from previously published circadian datasets were downloaded for *Arabidopsis*^26^, *Brassica rapa*^29^, and *Brachypodium distachyon*^27^. These reads were filtered for quality, and any remaining adaptor sequence trimmed with Trimmomatic^91^. Surviving reads were aligned using HISAT2^92^ to *A. thaliana* genome (TAIR 10), *B. rapa* genome (v1.0) and the *B. distachyon genome* (v3.0) respectively. For the *Arabidopsis* alignment, maximum intron length was set to 5000nt consistent with pre-processing in^26, 94^. StringTie^93^ was used to quantify uniquely mapping reads before TPM value extraction at gene level. For *Glycine max*^28^, FPKM normalised reads were downloaded from the *Glycine max* RNA-seq Database^95^ (accession GSE94228) and were converted from FPKM to TPM prior to analysis.

### Homolog identification of circadian clock and circadian controlled genes

Wheat homologs of *Arabidopsis* core circadian clock genes were identified in the wheat genome by detecting similarity to the following conserved protein family domains that are present in the proteins encoded by these genes: MYB1R, a subtype of MYB domain that contains a distinctive SHAQKY sequence motif (present in the CCA1, LHY and RVE[1-8]) or a distinctive SHLQKY sequence motif (present in LUX), PAS (present in ZTL), PRR (present in TOC1 and PRR[3579]) and ELF4 (present in ELF4). A hidden Markov model (HMM) for each domain was used in HMMER 3.2.1 HMMSEARCH^96^ to search for members of the domain family in the following proteome datasets: Araport11 (*Arabidopsis thaliana*), RGAP7 (*Oryza sativa*), JGI Phytozome version 12 (*Brachypodium distachyon*), IBSC (*Hordeum vulgare*), SpudDB PGSC v4.03 (*Solanum tuberosum*) and IWGSC Refseq v1.1 (*Triticum aestivum*). The HMMs provided by Pfam (https://pfam.xfam.org/) were used for the PAS domain (PAS_9, PF13426), the PRR domain (Response_reg, PF00072) and the ELF4 domain (PF07011). For the MYB domain, an HMM was built for the MYB1R subfamily using HMMER3 HMMBUILD^96^ with an alignment of protein sequences from *Arabidopsis* and rice, previously established as being members of this subfamily. The sequences found from these genomes were re-aligned to the original alignment using HMMER 3.2.1 HMMALIGN^96^. Amino acids with non-match states in the HMM were removed from the alignment and alignment columns with <70% occupancy were also removed. The longest splice variant of each protein was selected to estimate a phylogenetic tree with bootstrap support using RAxML 8.2.12^97^ with the following method parameters set: -f a, -x 12345, -p 12345, -# 100, -m PROTCATJTT. The trees were mid-point rooted and images created using the Interactive Tree of Life (iToL) tool^98^. For the larger MYB and PRR families, proteins from the tree clades containing known clock gene(s) were re-aligned across their full-length and a “nested” phylogenetic tree was re-estimated with RAxML as described above. The tree was visualised in the Interactive Tree Of Life (iTOL) website alongside the corresponding alignment. This view provided increased detail about the relationships within the clade and enabled orthologous sequences to be inferred. Wheat homologues for *ELF3*, *GI*, *LWD1/2*, *CHE,* and *LNK1/2* were identified by BLASTP searches using previously identified wheat and *Brachypodium* predicted proteins confirmed by reciprocal BLAST searches against *Arabidopsis*. IDs and source references can be viewed in Supplementary Table 8.

Putative wheat orthologs for *Arabidopsis* circadian controlled pathway genes involved in photosynthesis, light-signalling and primary metabolism were first extracted using Biomart v0.7^99^ available from Ensembl Plants and taken forward if they had >40% identity in the DNA sequence. Orthologs were then verified using BLASTP using *Arabidopsis* protein sequences as a query against the wheat protein database to confirm the wheat gene IDs. Complete lists of wheat gene IDs used in the pathway analysis can be viewed in Supplementary Table 11.

### Circadian quantification using Metacycle and Biodare2

To estimate proportions of rhythmic genes expressed in *Arabidopsis* and wheat, we removed only genes with 0 TPM at all timepoints. This approach has been used in several previous studies^26, 100, 101^ and allows detection of low-expression rhythmic transcripts. An analysis of how filtering for low-expression genes affects the estimates of proportions of rhythmically expressed genes is discussed in Supplementary Note 1 and Supplementary Table 1.

The R package MetaCycle^102^ was used to identify rhythmically expressed transcripts (Benjamini-Hochberg *q*-values) and to quantify period lengths (hours), absolute phase (hours), baseline expression (TPM), amplitudes (TPM) and relative amplitudes of circadian waveforms. Relative amplitude is the ratio between amplitude and baseline TPM if the baseline is greater than 1. Metacycle integrates results from three independent algorithms (ARSER, JTK_CYCLE and Lomb-Scargle) to produce summary “meta2d” statistics that combine the outcome from these algorithms. Metacycle was run using the following parameters; minper = 12, maxper = 35, adjustPhase = “predictedPer”. Transcripts were defined as rhythmic if they had *q*-values < 0.05 and high confidence rhythmic transcripts if they have *q*-values < 0.01. To calculate circadian phase (CT; relative to period length=24), meta2d phase estimates were multiplied by 24 and then divided by the period estimates for each transcript. Circular phase means were calculated using the package ‘circular’ implemented in R^103^.

There are many different algorithms available for quantification of rhythmicity within time-series data, some of which perform better on datasets with higher levels of noise, non-24h periods or various sampling frequencies. To validate the meta2d results we also used the FFT-NLLS and MESA algorithms implemented in Biodare2 to verify our observations about period, phase and rhythmicity^104^. FFT-NLLS also provides relative amplitude error (RAE) statistics which represent a useful metric for assessing rhythmic robustness. FFT-NLLS and MESA were run using the BH *q* < 0.01 filtered transcripts categorized in Metacycle, and with the following parameters: no dtr, min-max, p(12.0-35.0).

To enable as close a comparison with the *Arabidopsis* dataset as possible, the wheat time-course was cropped to a data window of 24-68h for approximation of period, phase and relative amplitude unless specified otherwise. This data-window also ensures that measurements are being made under circadian conditions following transfer to constant light. For the triad analysis, meta2d estimates were measured over the full time-course (0-68h) as differentiation of homeolog behaviour was the main interest, including the response to transfer to L:L.

### Clustering of rhythmic genes into expression modules

Gene co-expression analysis was carried out using the R package WGCNA (Langfelder and Horvath, 2008; R version 3.6.0.).

#### Arabidopsis

The 10,317 genes identified by MetaCycle as significantly rhythmic (q-value < 0.01) were filtered and genes with greater than 0.5 TPM average expression at more than three timepoints were retained for further analysis. The average expression at each timepoint for the remaining 10,129 genes was used to construct signed hybrid networks on a replicate basis using the blockwiseModules() function. The soft power threshold was calculated as 18, and the following parameters were used; minModuleSize = 30, corType = bicor, maxPOutliers = 0.05, mergeCutHeight = 0.15. Highly connected hub genes were identified for each of the 9 co-expression modules using the function chooseTopHubInEachModule() and eigengenes were identified for each module using the moduleEigengenes() function. Wheat: The 18,633 genes identified by MetaCycle as significantly rhythmic across 12 timepoints ZT24 - ZT68 (q-value < 0.01) were filtered and genes with greater than 0.5 TPM average expression at more than three timepoints were retained for further analysis. The average expression at each timepoint for the remaining 16,327 genes was used to construct signed hybrid networks using the blockwiseModules() function. A soft power threshold of 18 was used, together with the following parameters; minModuleSize = 30, corType = bicor, maxPOutliers = 0.05, mergeCutHeight = 0.15. Eigengenes were identified for each module using the moduleEigengenes() function. Modules with closely correlated eigengenes were merged using the mergeCloseModules() function, with the parameters; cutHeight = 0.25, iterate = F) and new module eigengenes were calculated for the resulting 9 modules.

### Cross-correlation analysis

A cross-correlation analysis was used to find the shift in time (lag) which produced the highest (peak) correlation between two rhythms. This approach was used to identify modules which peaked synchronously (had a peak lag of 0h) or asynchronously (had a peak lag of 4, 8 or 12h) by correlating eigengenes for each module (Supplementary Fig. 9). We also used cross-correlation to identify imbalanced phases within rhythmic triads (Fig. 1E). Before calculating the cross-correlation between two expression rhythms, we first scaled both expression patterns using their means and standard deviations, so the output reflects a time-dependent Pearson correlation coefficient ranging between -1 and 1:

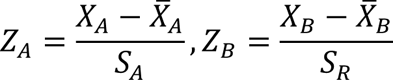

Where *Z_i_*, *X_i_*, 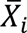 and *S_i_* represent the standardised expression level, tpm expression level, mean expression level, and standard deviation of gene A and B respectively. Once both expression patterns have been scaled, the discrete cross-correlation between the two expression patterns is calculated using the np.correlate function and is divided by the number of time points in the expression signal returning the Pearson correlation coefficient at different lags. The index of the array with the largest Pearson correlation coefficient score corresponds to the lag that maximises the phase similarity between the two temporal expression patterns.

### Mean-normalised data for oscillation plots

Oscillation plots in Supplementary Fig. 18-20 were mean normalised to aid visualisation of period and phase differences between transcripts. Data was adjusted by dividing the TPM values at each timepoint by the mean across all timepoints for each gene so that the baseline expression was equal to 1.

### Gene ontology term enrichment

Functional enrichment of differentially expressed genes for biological processes within each module was performed using the gene ontology enrichment analysis package, topGO^105^ in R (version 3.6.0, with the following parameters: nodeSize = 10, algorithm = “parentchild”, classicFisher test p < 0.05). Enrichment of terms in all rhythmic genes in *Arabidopsis* and wheat was compared against a background ‘gene universe’ of all expressed genes in each dataset (26,392 genes for *Arabidopsis* and 86,567 for wheat). This gene universe was also used in the GO-slim analysis for enrichment in circadian balanced versus imbalanced triads. Enrichment of terms in expression modules was compared against a background of all rhythmically expressed genes (BH *q* < 0.01) which clustered into modules in each dataset (10,129 genes for *Arabidopsis* and 16,327 for wheat). GO-slim terms refer to ontology terms for biological processes unless otherwise specified and were obtained from Ensembl Plants 51^11^, using the BioMart tool. The bubble plot was plotted using ggplot in R adapting code from De Vega et al., 2021^106^.

Enrichment of GO-slim terms in *B. rapa* circadian paralogs with similar and differential expression patterns was conducted using previously published DiPALM results for pattern change in a LDHC circadian time course (Supplementary File 4, Greenham et al. 2020 ^29^). Paralog pairs with a pattern change *p-*value of <0.001 were termed differential patterns and pairs with a pattern change *p*-value of >0.1 were considered to be similar patterns. In this analysis we ignored differences in expression change for consistency with our wheat triads. Data was first filtered for rhythmicity using Metacycle q-values <0.01. Only paralog pairs with two significantly rhythmic paralogs were retained for the GO-slim analysis. Enrichment of terms in similarly expressed circadian paralogs (1562 genes) or differentially expressed paralogs (1438 genes) in B. napus was compared against a background of 4646 genes expressed in the Greenham dataset in paralogs and which had GO-slim annotation available.

### Enrichment analysis of transcription factor superfamilies in wheat co-expression modules

Genes annotated as members of transcription factor superfamilies^30^ were identified in each co-expression module and the frequency of each TF superfamily compared to the frequency observed in the 16,327 genes submitted to WGCNA. TF families were classed as either significantly under or overrepresented in each module using Fisher’s exact test (p <= 0.05).

### Enrichment analysis of transcription factor binding sites in wheat co-expression modules

1.5 kb of sequence upstream of the transcription start site (TSS) was extracted for each of the 16,327 genes submitted to WGCNA. FIMO, from the MEME tool suite (v 4.11.1) was used to predict TFBS in these regions based on similarity with previously DAP-seq validated TFBS identified in *Arabidopsis*^107^. FIMO was run as reported in Ramírez-González et al., 2018 (p-value threshold of <1e-04 (default), --motifpseudo set to 1e-08 as recommended for use with PWMs and a --max-stored-scores of 1,000,000). The background model was generated from the 16,327 promoter sequences using MEME fasta-get-markov. As the significance of multiple matches of a single TFBS family in the putative promoter region for each gene is unknown, we derived a non-redundant (nr) list of matched TFBS motifs for each gene within each of the nine modules and for the complete set of 16,327 genes, where multiple occurrences of a TFBS superfamily in a single promoter sequence were only counted once. The frequency of these nrTFBS motifs for each co-expression module was compared to the frequency of nrTFBS seen across all 16,327 genes and families significantly under or overrepresented in each module were identified using Fisher’s exact test (*p* <= 0.05).

### Loom plots

Genome position data for the plots are based on annotations from Chinese spring (Triticum_aestivum.IWGSC.52.gtf) downloaded from Ensembl Plants. Code for creating Loom plots (Supplementary Fig 8) is implemented in R and a code package with accompanying R markdown notebooks is available from our groups GitHub repository (https://github.com/AHallLab/triad.expression).

### Statistical analysis

Statistical tests including Welch’s two sample t-test, Two-proportions z-test, One-way ANOVA, Two-level, nested ANOVA and Chi-square tests of independence were all conducted in the R ‘stats’ package (version 4.0.0) with default parameters.

## Data availability

Fastq data from the RNA-seq circadian time course are available to view from the Grassroots Data Repository: https://opendata.earlham.ac.uk/opendata/data/wheat_circadian_Rees_2021. *[Data will be uploaded to the European Nucleotide Archive (ENA) during the review process]* A summary csv table with expression of wheat genes (TPM), Metacycle estimates, gene annotations and triad balance classification can be viewed in Supplementary Table 12 also available here: https://opendata.earlham.ac.uk/opendata/data/wheat_circadian_Rees_2021.

## Acknowledgements

H.R., R.R.P. and A.H. were funded by the BBSRC Core Strategic Programme Grant (Genomes to Food Security) BB/CSP1720/1 and its constituent work package, BBS/E/T/000PR9819 (WP2 Regulatory interactions and Complex Phenotypes). B.W., R.R.P. and A.H. was supported by the BBSRC Designing Future Wheat grant BB/P016855/1; BBS/E/T/000PR9783 (DFW WP4 Data Access and Analysis). C.R. by a BBSRC grant BB/V509267/1 and Wave 1 of The UKRI Strategic Priorities Fund under the EPSRC Grant EP/T001569/1, particularly the “AI for Science” theme within that grant & The Alan Turing Institute. J.C. by the BBSRC funded Norwich Research Park Biosciences Doctoral Training Partnership grant BB/M011216/1. We would also like to acknowledge BBS/E/T/000PR9816 (NC1 - Supporting EI’s ISPs and the UK Community with Genomics and Single Cell Analysis) for data generation and BB/CCG1720/1 for the physical HPC infrastructure and data centre delivered via the NBI Computing infrastructure for Science (CiS) group. P.B. was supported by a BBSRC TRDF grant BB/N023145/1. A.N.D., L.L.B.D. and C.A.G. are funded by BBSRC ISP Genes in the Environment (BB/P013511/1). C.A.G. was also funded by UK BBSRC SWBIO DTP (BB/M009122/1). We thank Dr. Susan Duncan for her contribution to preliminary experiments prior to the start of this project.

## Author contributions

H.R. and A.H. were involved with project concept and designed experiments. H.R. performed time-course experiments. R.R.P. processed data, quantified read counts and conducted clustering of rhythmic transcripts. P.B. conducted the phylogenetic analysis of core circadian protein families. J.C. conducted the cross-correlation analysis. C.R. processed previously published circadian datasets. S.J.W produced loom plots for circadian triad balance. H.R., L.L.B.D., C.A.G., B.W., R.R.P. A.H., and A.N.D. analysed and interpreted the RNA-seq data. H.R. wrote the initial manuscript and all authors contributed to subsequent drafts.

## Ethics declarations

Competing interests: The authors declare no competing interests.

