## Supplementary Files for "Circadian regulation of the transcriptome in a complex polyploid crop"

### Supplementary Information: Circadian regulation of the transcriptome in a complex polyploid crop

**Contents:**

Contents: 1

Supplementary Note 1: Proportions of rhythmic genes in expression datasets 5

Supplementary Note 2: Mean period lengths in rhythmic wheat transcripts over sliding windows of circadian experiment 5

Supplementary Note 3: Distribution of phases across different period bins for rhythmic wheat transcripts 6

Supplementary Note 4: Examples of TF triads with unbalanced regulation 6

Supplementary Note 5: Detecting patterns of triad circadian balance 6

Supplementary Note 6: Orthologs co-expressed in similar phased modules of gene expression 7

Supplementary Note 7: Orthologs of further circadian clock components 8

Supplementary Note 8: Considerations for the differences in circadian regulation observed between *Arabidopsis* and wheat 8

Supplementary Information-Regulation of the circadian transcriptome in a complex polyploid crop


Supplementary Figure 1: Correlation of wheat period, circadian phase and absolute amplitude values estimated from meta2d or FFT-NLLS. ____ 10

Supplementary Figure 2: Period estimates for rhythmically categorized data predicted using either Metacycle or Biodare2. ____ 11

Supplementary Figure 3. Phase estimates for highly-rhythmically categorized data from either Metacycle or Biodare2 ____ 12

Supplementary Figure 4. Changes in phase distributions in each of the wheat transcript period groups over three sliding windows relative to dawn. ____ 13

Supplementary Figure 5. Comparison of balanced triads as classified by Ramírez-González et al., and circadian balanced triads classified in this study. ____ 14

Supplementary Figure 6. Examples of how definition of transcriptional triad balance can change in circadian time. ____ 15

Supplementary Figure 7. Expression of homeologs of key transcription factors and their putative downstream targets. ____ 17

Supplementary Figure 8. Loom plots showing positions of runs of imbalanced circadian triads. ____ 18

Supplementary Figure 9. *Arabidopsis* and wheat modules with a peak lag of 0 (synchronous phase) share more GO-slim terms than non-synchronous modules. ____ 24

Supplementary Figure 10. Transcription factor superfamily enrichment in wheat modules. _27

Supplementary Figure 11. Transcription factor binding site (TFBS) superfamily enrichment in wheat modules. ____ 30

Supplementary Figure 12. Phylogenetic relationships for circadian genes based on alignment of LHY-like MYB full length proteins. ____ 32

Supplementary Figure 13. Phylogenetic relationships for circadian genes based on alignment of TOC1-like and PRR full length proteins. ____ 33

Supplementary Information-Regulation of the circadian transcriptome in a complex polyploid crop


Supplementary Figure 14. Phylogenetic relationships for circadian genes based on alignment of PAS/LOV protein domains. ____ 36

Supplementary Figure 15. Phylogenetic relationships for circadian genes based on alignment of LUX-like MYB full length proteins. ____ 37

Supplementary Figure 16. Phylogenetic relationships for circadian genes based on alignment of ELF4-like protein domains. ____ 38

Supplementary Figure 17. Additional free-running expression of core circadian clock genes in wheat and their homologs in *Arabidopsis*. ____ 40

Supplementary Figure 18. Arabidopsis and Wheat photosystem genes under free-running L:L conditions. ____ 42

Supplementary Figure 19. *Arabidopsis* and wheat light signalling genes under free-running L:L conditions. ____ 45

Supplementary Figure 20. *Arabidopsis* and wheat primary metabolism genes under free-running L:L conditions. ____ 49

Supplementary Table 1: Proportions of rhythmically classified genes using datasets filtered to remove low expression genes. ____ 50

Supplementary Table 2: Meta2d estimates from previously published circadian datasets. ____ 51

Supplementary Table 3: GO-slim terms for biological processes associated with similar patterns of circadian expression or different patterns of circadian expression in Brassica rapa paralogs. ____ 52

Supplementary Table 4: Expected and Observed runs of Rhythmic triad categories across the wheat genome. ____ 53

Supplementary Table 5. Characteristics of circadian rhythms in wheat genes split into two hour period bins. ____ 55

Supplementary Table 6. Circadian characteristics of module eigengenes. ____ 55

Supplementary Table 8: Gene IDs for core circadian genes and corresponding *Arabidopsis* orthologs ______ 57

Supplementary references: ______ 66

**Supplementary Tables as independent files:**

Supplementary Table 7: GO-slim terms for each of the nine modules identified in *Arabidopsis* and wheat.

Supplementary Table 9: Metacycle (Meta2d) estimates for core circadian genes and corresponding *Arabidopsis* orthologs.

Supplementary Table 10: Enriched GO-slim terms in rhythmically expressed genes in *Arabidopsis* and wheat.

Supplementary Table 11: Expression data and Metacycle statistics for transcripts involved in key circadian regulated pathways.

Supplementary Table 12: Summary table with expression of wheat genes (TPM), Metacycle estimates across (0-68h and 24-68h), gene annotations and triad balance classification

Supplementary Information-Regulation of the circadian transcriptome in a complex polyploid crop

**Supplementary Note 1: Proportions of rhythmic genes in expression datasets**

The proportion of the wheat transcriptome categorized as rhythmic depends on the window over which measurements are taken, the *q*-value cut-off for rhythmicity and the threshold at which you exclude low expression transcripts. We chose to conduct our primary analysis without filtering for any minimum expression to allow us to capture all rhythmically categorized transcripts. This approach has been used in several previous studies [1–3] and comparison of relative proportions in *Arabidopsis* and wheat is possible as identical criteria are applied to both datasets as shown in Table 1 in the main text. To assess the effects of applying a pre-filtering step to exclude low-expression genes, we removed genes with expression less than 0.1 TPM in six or more timepoints and again ran Metacycle to define numbers of rhythmic genes. This had the effect of reducing the number of genes defined as expressed (and therefore the proportions of rhythmic genes) but had only a small effect on the numbers of rhythmically categorized genes as shown in Supplementary Table 1. With the filter applied to the wheat 24-68h dataset, the number of rhythmic genes reduced by 206 genes at $q<0.05$, and increased by 752 genes at $q<0.01$ which had a minimal impact on mean period and relative amplitudes. However, the proportion of rhythmic genes increased from 33.56% to 46.3% in wheat and from 50.7% to 61.5% in *Arabidopsis* due to the lower expression totals. Regardless of whether a filtering step was applied or whether a $q<0.05$ or $q<0.01$ threshold was used, the proportion of genes in *Arabidopsis* was found to be consistently higher than in wheat ($p < 0.001$, one-tailed, two-proportions z-test).

**Supplementary Note 2: Mean period lengths in rhythmic wheat transcripts over sliding windows of circadian experiment**

We wanted to investigate how period length changed over the time-course following transfer to constant light (L:L). Mean period lengths were calculated for rhythmic transcripts (B.H $q < 0.01$) over a sliding window of 0-44h, 12-56h or 24-68h relative to transfer to L:L. Unexpectedly, the mean period length of all rhythmic wheat genes at 0-44h was even longer at 28.61h (SD=3.421), which shortened over a window of 12-56h to 27.31h (SD=3.461) and finally shortened again to 26.82h (SD=3.208) over 24-68h. This suggests that the period length of rhythmic genes entrained under 24h days became much longer immediately after transfer to constant light (L:L) and then progressively got shorter again over the following two days. There was no correlation between period length of individual genes measured in the first window versus the last window ($R^2$ =0.02, *p*-value =0.06) therefore genes defined as 'short' and 'long" period under 24-68h were not consistently short or long immediately after transfer to L:L.

**Supplementary Note 3: Distribution of phases across different period bins for rhythmic wheat transcripts**

Using the 24-68h data window, we split genes into 5 groups based on their period length; Group A: 21-23h (808 genes), Group B: 23-25h (2229 genes), Group C: 25-27h (2506 genes), Group D: 27-29h (1940 genes), Group E: 30-31h (761 genes). Genes in group C (25-27h) had the highest rhythmicity (defined by q-values and RAE scores) and largest relative amplitudes of the five groups (Supplementary Table 2). When we analysed histograms for phase of transcripts in each period bin, we found that groups A and B contained proportionally more dawn-peaking genes than the other groups, whereas groups D and E contained proportionally more dusk-peaking genes. These phase distributions also shifted over the course of the experiment as can be seen in Supplementary Figure 4.

**Supplementary Note 4: Examples of TF triads with unbalanced regulation**

As case studies, we investigated two transcription factor triads with unbalanced circadian expression that make known contributions to the productivity of wheat. *WCBF2* (aka *TaCBF1*) participates in cold signal transduction in wheat and is induced by drought and low temperature gated by the circadian clock (Badawi, 2007, Kume 2005, Fowler 2005, Jaglo 2001). The TCP transcription factor *PCF5* participates in cold tolerance and leaf development in rice and switchgrass (Yang 2013, Xie 2017). Interestingly, homologs of putative targets of WCBF2 *(WDHN13* and *WRAB17)* and TaPCF5 (*HY5, XTH27, NIA1, NIA2* and *OsPCNA)* also had unbalanced rhythmicity within their triads, although not necessarily with the same dominant homoeolog as their transcription factor (Supplementary Figure 7).

**Supplementary Note 5: Detecting patterns of triad circadian balance**

We looked at triads with unbalanced rhythmicity (one rhythmic gene q <0.01 (2058 triads) or two rhythmic genes q <0.01(1392 triads)) and triads with balanced rhythmicity or three rhythmic genes q<0.05 (3448 triads). For each gene, we found the corresponding chromosome positions based on the IWGSC RefSeq v1.1 gene annotation in cv. Chinese spring. In our dataset, we found 4225 triads with positional information and with either 1, 2 or 3 rhythmic homoeologs. The probability of a triads having 1 rhythmic homoeolog was 45%, 2 rhythmic homoeologs was 30% and 3 rhythmic homeologs was 25% based on their occurrence in the real data. Using the formula:

$$\left|\frac{\ln(n)}{\ln(P)}\right|$$

Where:

n = the total number of triads, P = the probability of each triad class occurring, ln = natural log and |…| = the absolute value

we could calculate the longest expected run of each category of rhythmic triads. Using the Wald–Wolfowitz test, we could also statistically test the randomness of each category to see whether rhythmic triads were grouping together more often than would be expected by chance. The null hypothesis is that all elements in a serial list are independent. The results, shown in Supplementary Table 4, suggest that sets of balanced and unbalanced triads are not grouped together in the genome and, likewise, that there are no regions in the genome where absence of rhythmicity from a particular chromosome is clustered more than would be expected by chance.

**Supplementary Note 6: Orthologs co-expressed in similar phased modules of gene expression**

We used Biomart tool in Ensembl plants to identify *Arabidopsis* orthologs for rhythmically expressed wheat genes in each expression module based on percentages of nucleotide identity >40%. We then compared these putative orthologs with *Arabidopsis* genes expressed in similar phased *Arabidopsis* modules to find co-expressed genes. In dawn-phased A9 and W9, we found 45 ribosomal protein orthologs comprising both large and small ribosomal subunits. Other co-expressed genes in these modules included *CHALCONE SYNTHASE (CHS)*, an anthocyanin biosynthesis enzyme also involved in salicylic acid defence; *PHOSPHATE STARVATION RESPONSE 1 (PHR1)*, involved in maintenance of photosynthesis under light stress and phosphate deficiency; *UV RESISTANCE 2* responsible for repairing UV radiation-induced DNA damage, and a heat shock protein, *HSP60* which ensures correct folding of proteins in mitochondria [4–7]. This suggests that in both *Arabidopsis* and wheat, the circadian clock initiates similar changes in ribosome biosynthesis and biotic and abiotic defence in anticipation of dawn. Genes peaking in both W3 and A2 included several light-harvesting complex proteins and *PHYTOCHROME INTERACTING FACTOR (PIF5)* as well as a cytochrome P450 monooxygenase gene *CYP709B3* which has previously been shown to be circadian regulated in *Arabidopsis* and may protect the plant from transpiration-triggered salinity stress during the day [8,9]. Genes co-expressed in W4 and A5 included several involved in auxin transport and signalling such as: the polar auxin transporter *AVP1*, *AUXIN F-BOX PROTEIN 5, AUXIN RESPONSE FACTOR 6, AUXIN SIGNALING F-BOX 2* and the endosomal sorting complex protein *CHMP1A* which ensures proper sorting of auxin carriers [10]. Orthologs expressed in wheat and *Arabidopsis* modules W5 and A6 included: *SEVEN IN ABSENTIA 2* which has been shown to regulate ABA-mediated stomatal closure and drought tolerance in *Arabidopsis* [11], and *HYDROPEROXIDE LYASE 1* which has a role in responding to insect attack and mechanical wounding [12]. Enriched MYB-TFs in W8 and W9 modules included *ASYMMETRIC LEAVES 1 (AS1)* involved in leaf morphogenesis, *GONIDIALESS A/ZUOTIN RELATED FACTOR A1* involved in plant growth, *G2-LIKE FLAVONOID REGULATOR (GFR*) involved in regulation of flavonoid biosynthesis and *LATE MERISTEM IDENTITY2* involved in transition from vegetative growth to flowering (*LMI2*). TFBS for MYB- and MYB-related-TFs were enriched in the W2 and W3 morning peaking modules and in night expressed modules W6, W7 and W8 in keeping with the broad phase range of the MYB TF expression.

**Supplementary Note 7: Orthologs of further circadian clock components**

We identified two *TaFKF1* transcripts that peaked at dusk and showed high similarity to *AtFKF1* in the PAS-domain phylogenetic tree (Supplementary Figure 14). Six transcripts similar to both *ZTL* and *LKP2* were not significantly rhythmic (Supplementary Figure 17), consistent with circadian regulation in *Arabidopsis*[13,14]. Wheat transcripts orthologous to *LNK1/2* and *CHE* also accumulated rhythmically, however *TaCHE* peaked antiphase to *AtCHE,* at ~CT20 rather than CT8 (Supplementary Figure 17). Two putative wheat *LWD* transcripts were not rhythmically expressed (Supplementary Figure 17).

**Supplementary Note 8: Considerations for the differences in circadian regulation observed between *Arabidopsis* and wheat**

In the main text we observe differences in circadian regulation of several transcripts encoding proteins involved in primary metabolism, and tentatively suggest that this may reflect a less pervasive influence of the circadian clock on Tre6P and starch metabolism in wheat.
An alternative hypothesis could be that regulatory control is conveyed by transcription of enzymes or transcription factors other than the *Arabidopsis* orthologs studied here.
In wheat, the role of starch is different in photosynthetic organs (source tissues) compared to grain producing organs (sink tissues), and the source:sink ratios change throughout wheat development. Therefore, regulation by the clock may change substantially between wheat tissues, organs and developmental stages. For instance, genes which are rhythmically expressed in the shoot may not be rhythmic in the root or genes which are not rhythmic in two-week old seedlings may become rhythmic during reproductive developmental stages. Further work is needed to elucidate tissue specific differences in circadian regulation in crops with pronounced source:sink relationships.
Additionally, rhythms in transcript abundance do not necessarily translate into rhythms in protein synthesis, assembly or post-translational regulation of activity. For example, post-translational phosphorylation regulates the activity of several wheat starch branching enzymes and starch synthases [15,16]. Future studies could explore whether differences in rhythmicity at the transcriptional level are reflected in altered oscillations of key enzymes and metabolites.

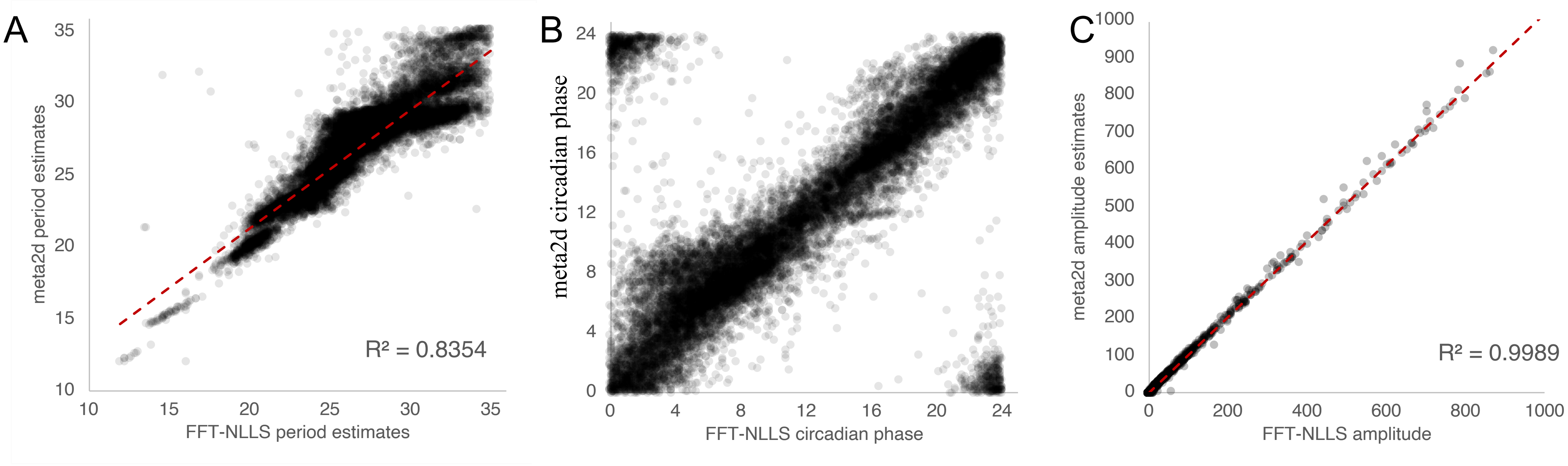


**Supplementary Figure 1: Correlation of wheat period, circadian phase and absolute amplitude values estimated from meta2d or FFT-NLLS.** Meta2d was run in Metacycle and FFT-NLLS in Biodare2 using 24-68h data filtered for rhythmicity B.H q-values <0.01.

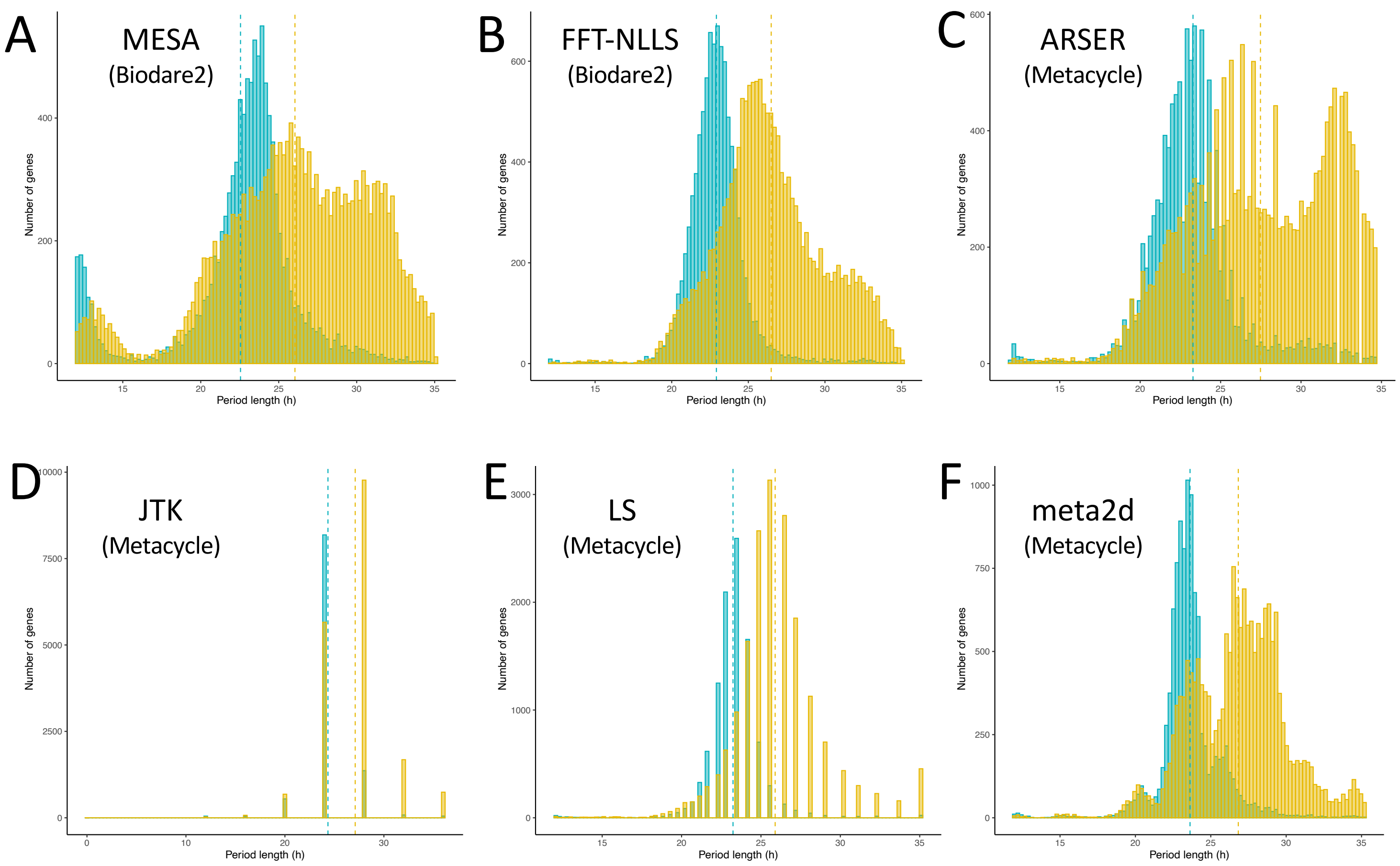


**Supplementary Figure 2: Period estimates for rhythmically categorized data predicted using either Metacycle or Biodare2.** MESA and FFT-NLLS are independently run in Biodare2 (A and B), ARSER, JTK and LS (C-E) are all run through Metacycle to produce an average period prediction meta2d (F). Data for *Arabidopsis* (blue) and wheat (yellow) was filtered for BH q<0.01 on a data window of 24-68h after transfer to constant light.

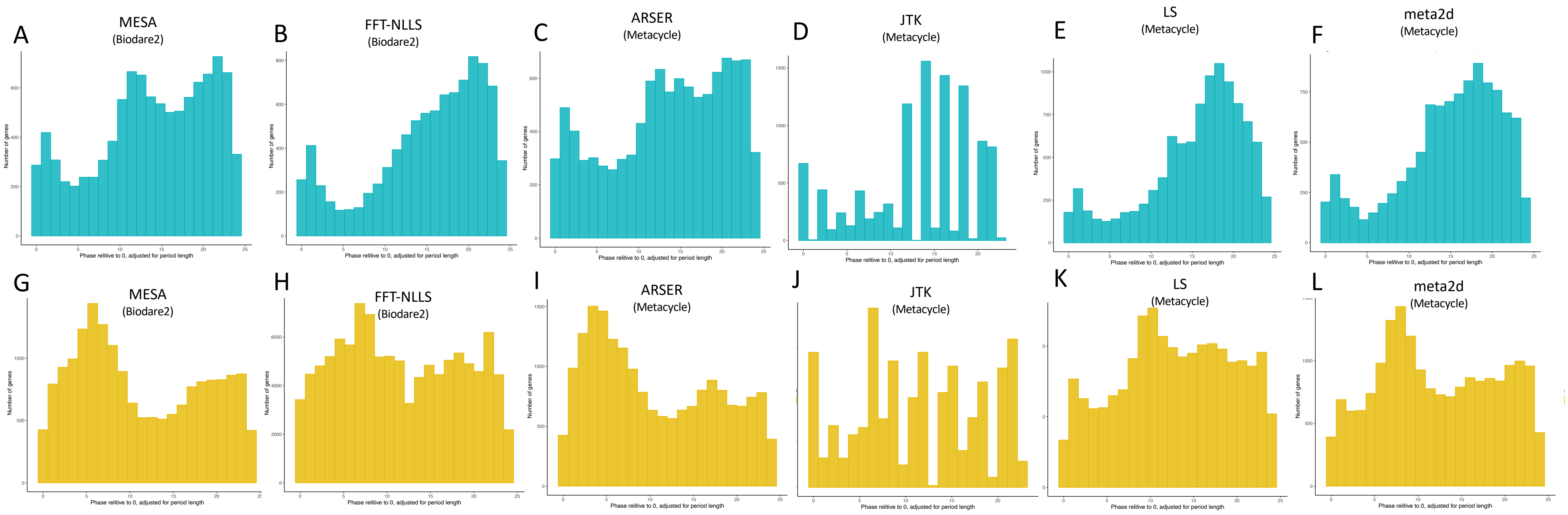


**Supplementary Figure 3. Phase estimates for highly-rhythmically categorized data from either Metacycle or Biodare2.** Phases were adjusted by period length so that one complete cycle is equal to 24h. MESA and FFT-NLLS are independently run in Biodare2 (A and B, G and H), ARSER, JTK and LS (C-E, I-K) are all run through Metacycle to produce an average period prediction meta2d (F and L) Data for *Arabidopsis* (blue) and wheat (yellow) was filtered for BH q<0.01 on a data window of 24-68h after dawn.

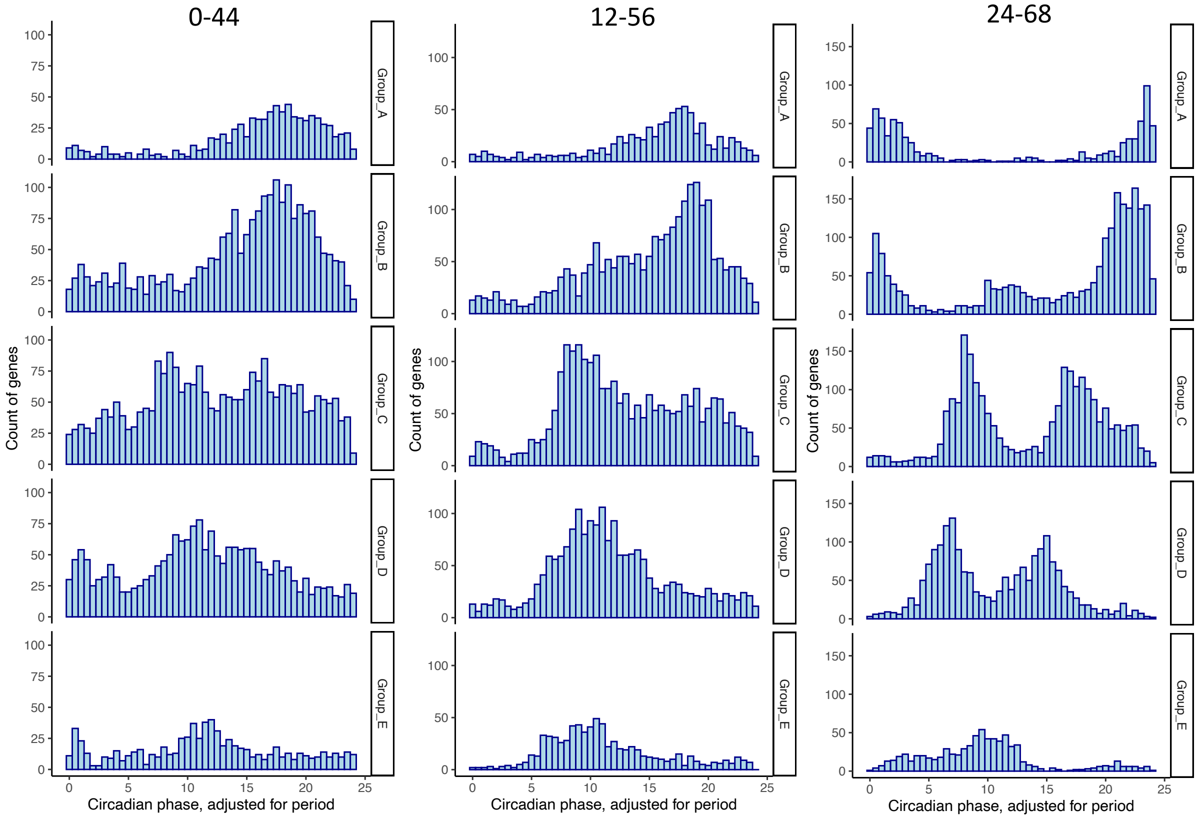


**Supplementary Figure 4. Changes in phase distributions in each of the wheat transcript period groups over three sliding windows relative to dawn.** Rhythmic genes were defined as having Metacycle q-values of <0.01, and individuals were placed in a period category only if the meta2d (Metacycle) period estimate was confirmed with independent estimates from FFT-NLLS (Biodare2). Phases are relative to period length (phase x 24/period) per gene. Period categories are as follows: Group A: 21-23h, Group B: 23-25h, Group C: 25-27h, Group D 27-29, Group E 30-31.

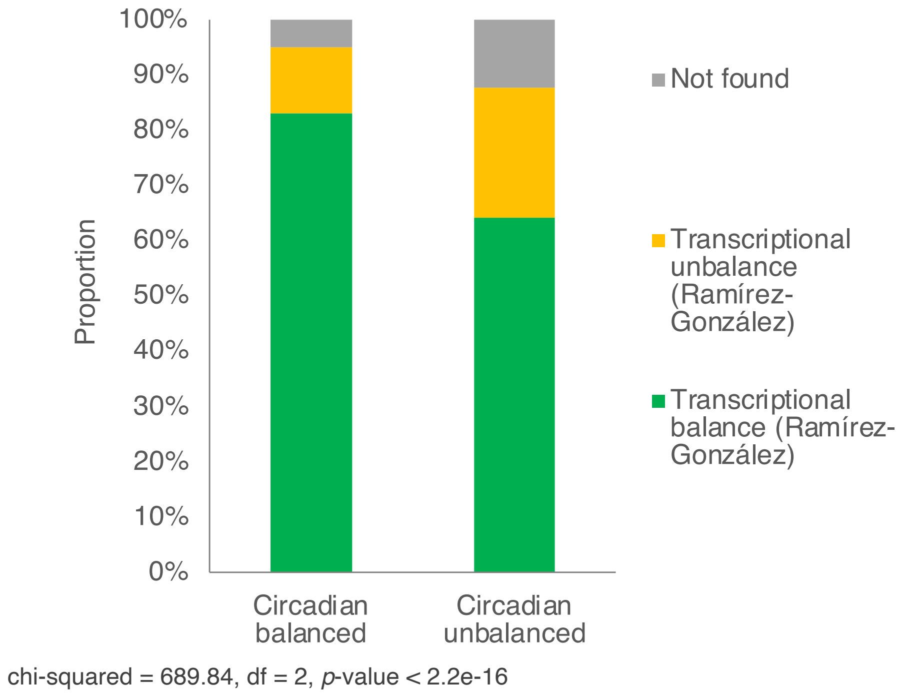


**Supplementary Figure 5. Comparison of balanced triads as classified by Ramírez-González et al., and circadian balanced triads classified in this study.** A chi-square test was used to determine whether there was a significant difference between the proportion of balanced and imbalanced triads (as defined by Ramírez-González) in each of the circadian-balanced and circadian-imbalanced triad categories. Results indicated that 83.20% of circadian-balanced triads were also categorized as balanced in the Ramírez-González data, whereas 64.15% of circadian-imbalanced triads were categorized as balanced in the Ramírez-González data. This difference was significant, $\chi2(2) = 689.8$, $p<0.00001$. These data also show that a large proportion of circadian triads labelled as balanced in the Ramírez-González dataset would be expected to have imbalanced patterns of expression if measured over several timepoints under circadian conditions, highlighting the importance of considering temporal dynamics in transcriptomic studies. Ramírez-González data is from Chinese spring leaves (excluding flag-leaf) under non-stressed conditions to provide as close a match to our data as possible. The diurnal time of collection is unknown.

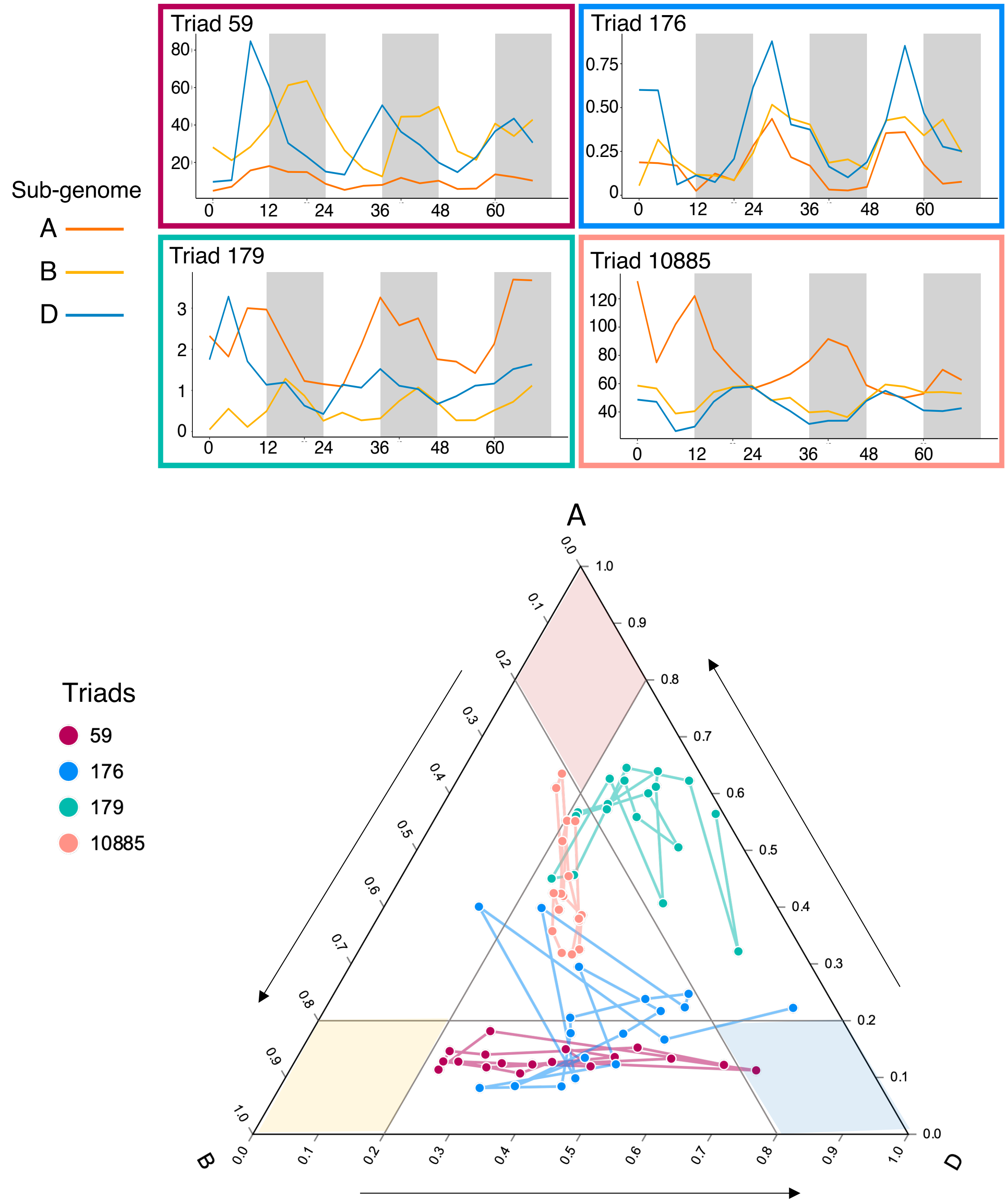


**Supplementary Figure 6. Examples of how definition of transcriptional triad balance can change in circadian time.** Each point represent expression normalized to 1 within each triad. Lines connect the dynamic changes in balance across the time-course. Triad 59: A suppressed across all timepoints, but dominance of B and D vary across the time-course. All homoeologs are classified as rhythmic (q<0.05) but have unbalanced phases.

Triad 176: Mostly appears as balanced across the time-course but occasionally looks as though A is suppressed. All homeologs in this triad are rhythmic ($q<0.05$) but have unbalanced relative amplitudes and periods. Triad 179: B suppressed over most timepoints, but when A homoeolog has a trough of expression appears as balanced. This triad has unbalanced phases and periods but all three homeologs are rhythmic ($q<0.05$).
Triad 10885: In this case, the A homoeolog is antiphase to the other homoeologs, and so in dawn timepoints the triad appears balanced, but in dusk timepoints there is A dominance. Ternary plots were created using https://www.ternaryplot.com/ .

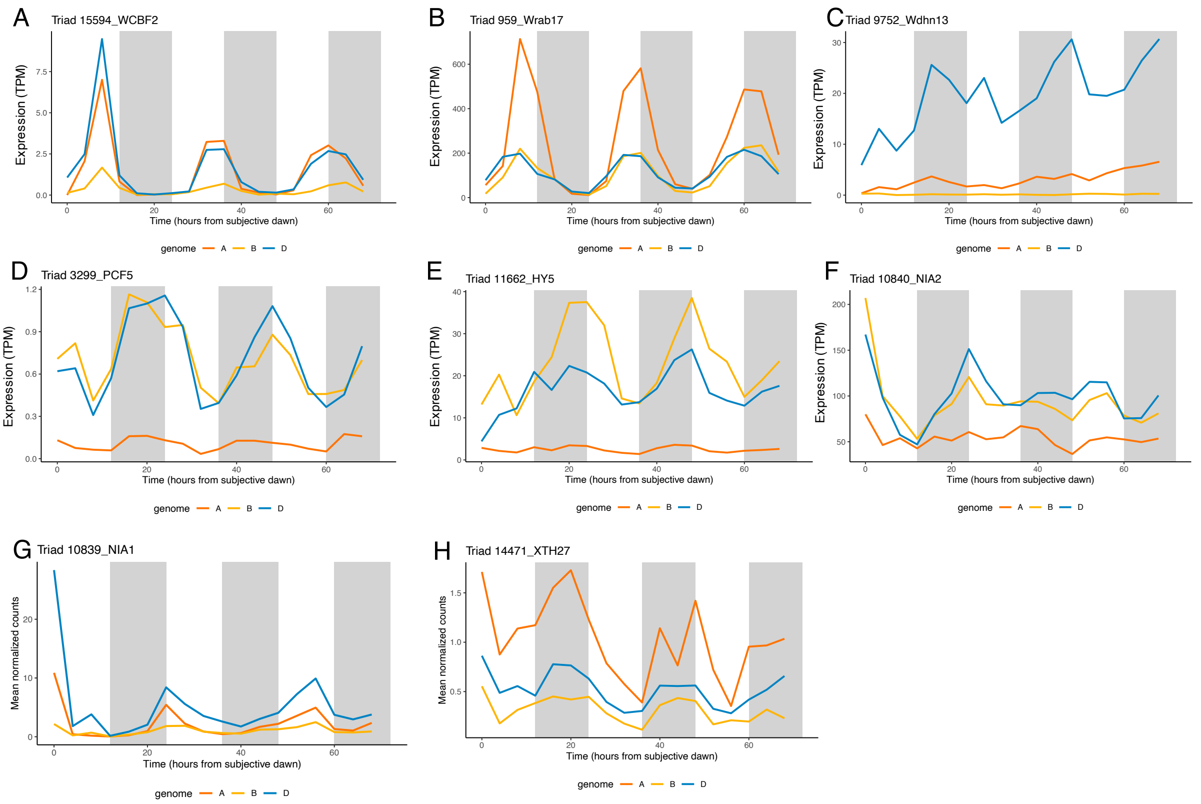


**Supplementary Figure 7. Expression of homeologs of key transcription factors and their putative downstream targets.** Wheat WCBF2 (A) putatively regulates *WDHN13* (B) and *WRAB17* (C), TaPCF5 (D) putatively regulates wheat orthologs of *HY5 (E), NIA2 (F), NIA2 (G)* and *XTH27 (H)*.

Supplementary Information-Regulation of the circadian transcriptome in a complex polyploid crop

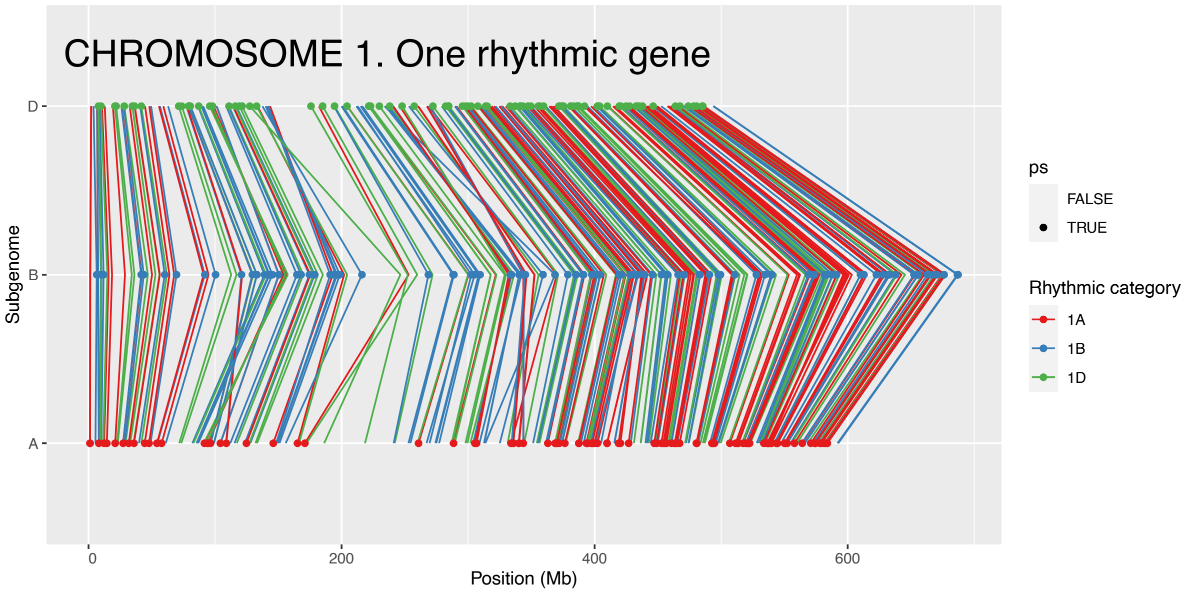


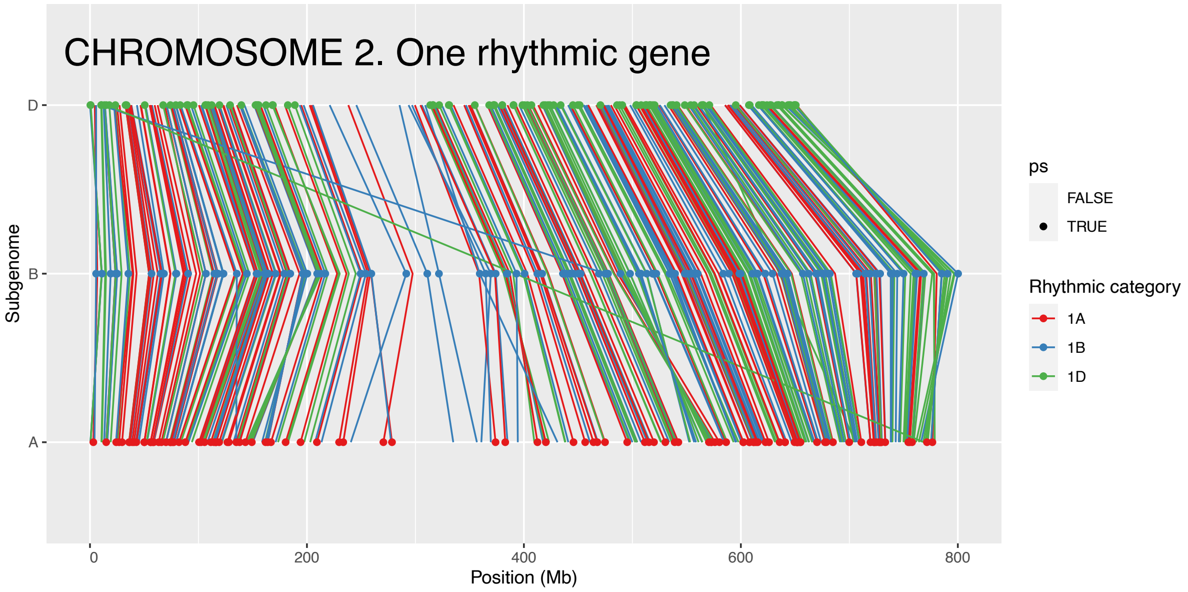


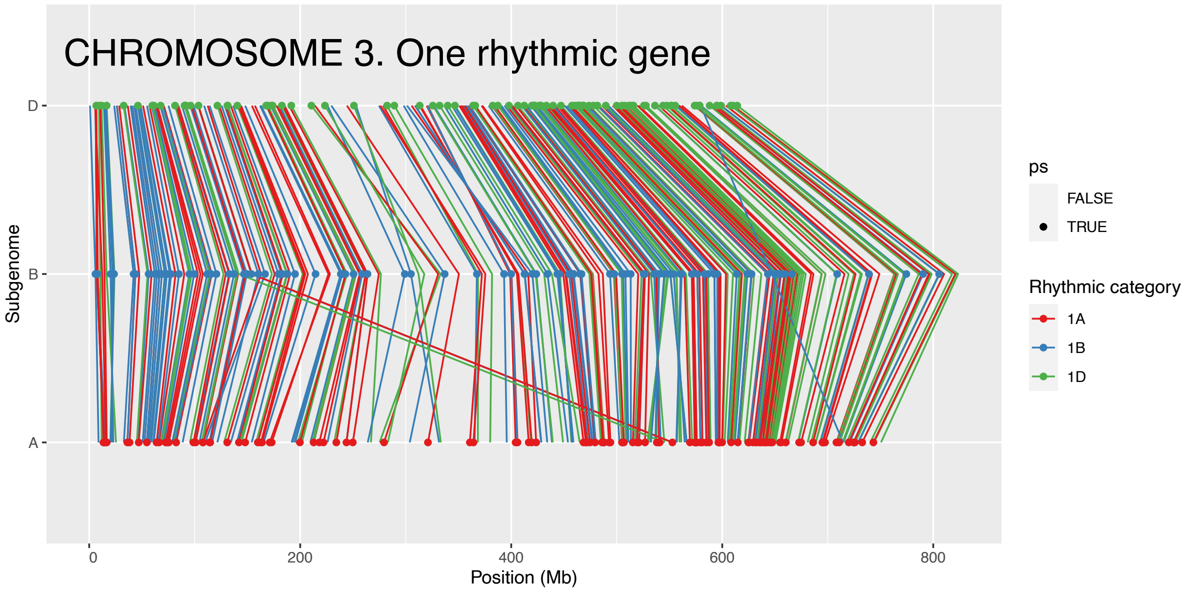

Supplementary Information-Regulation of the circadian transcriptome in a complex polyploid crop

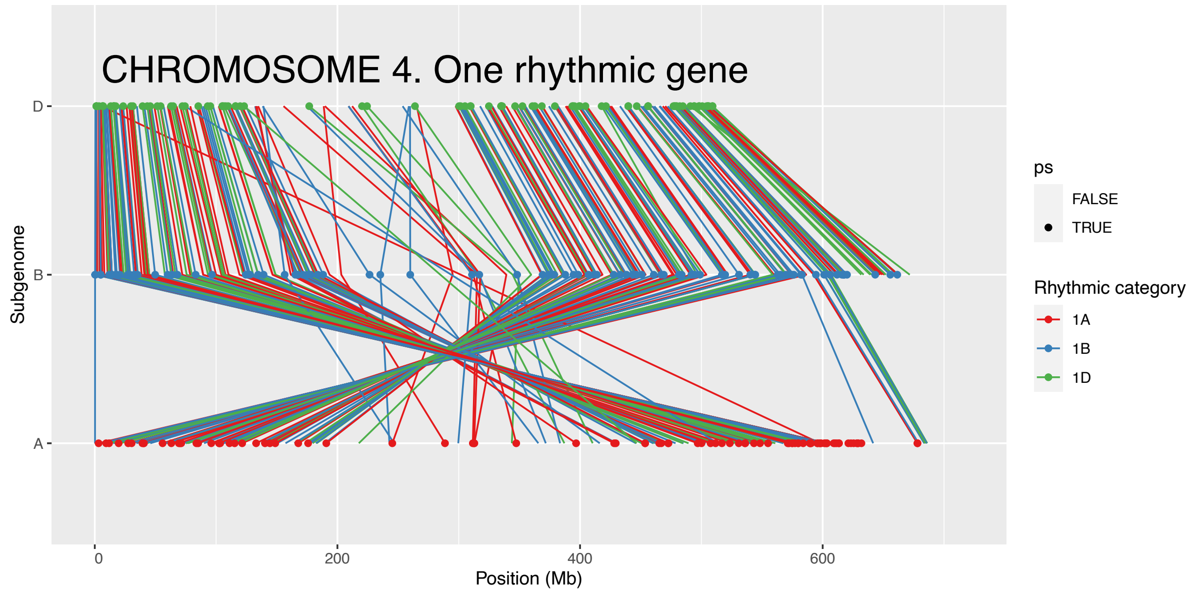


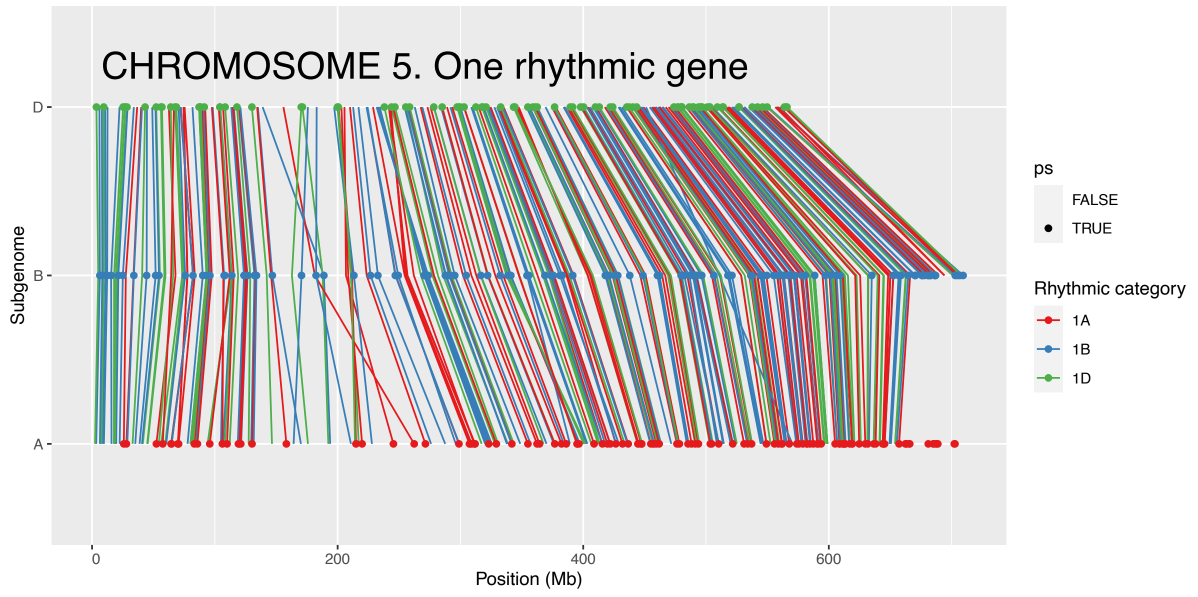


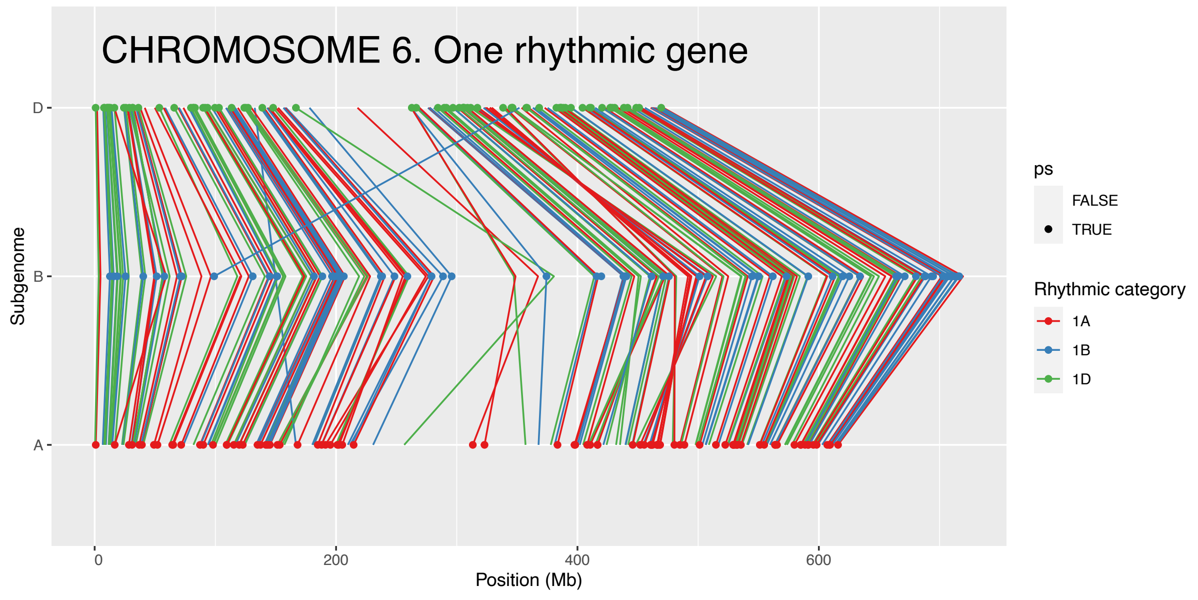

Supplementary Information-Regulation of the circadian transcriptome in a complex polyploid crop

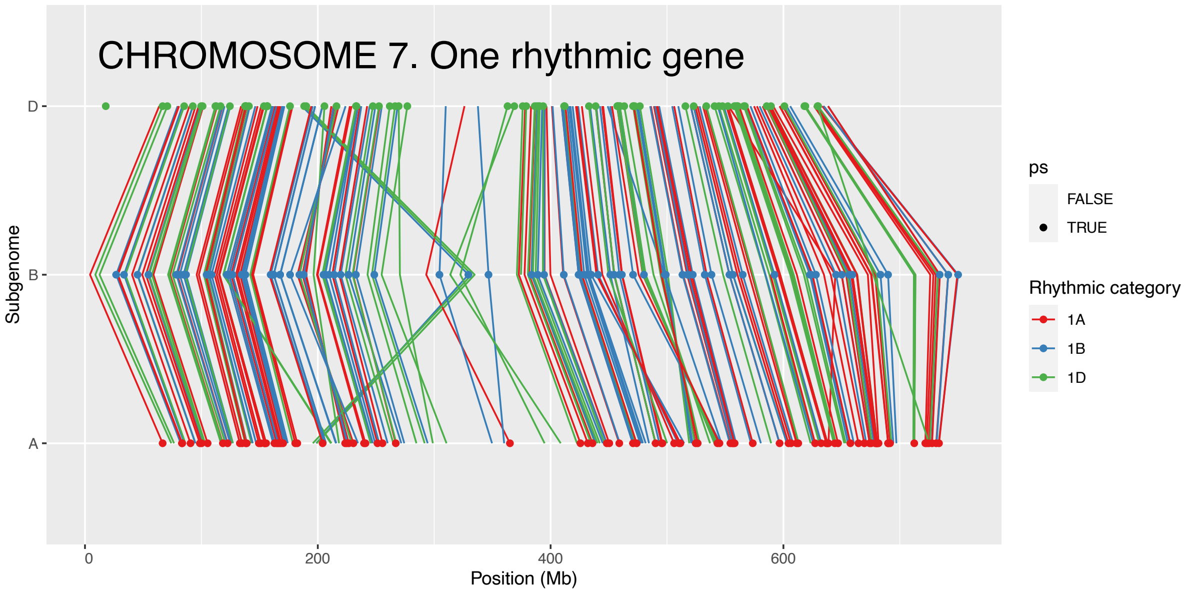


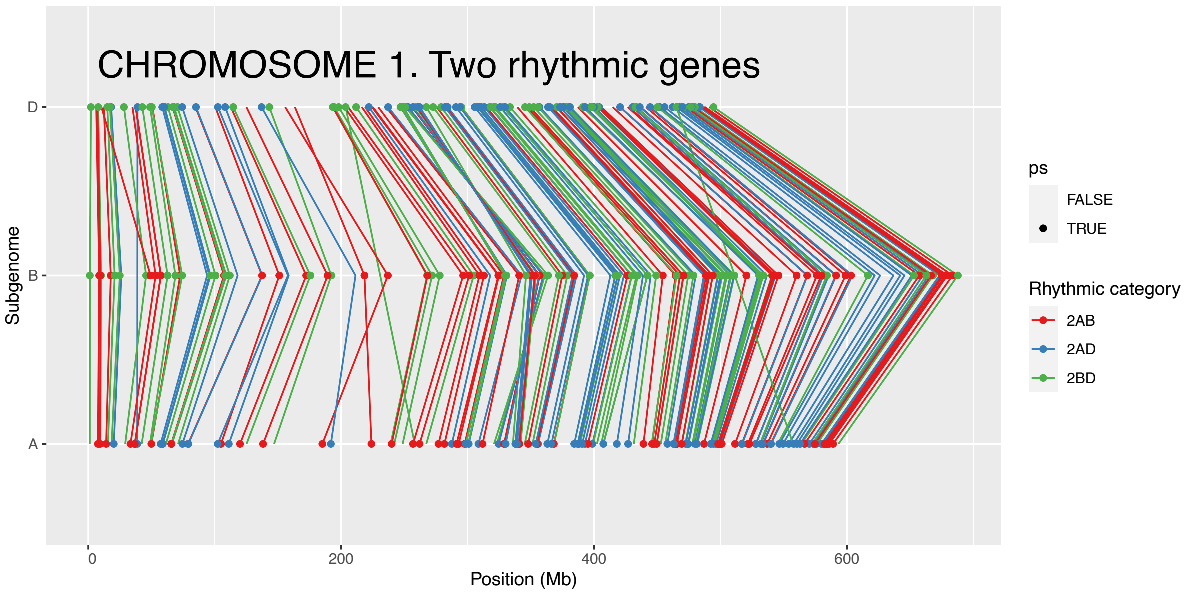


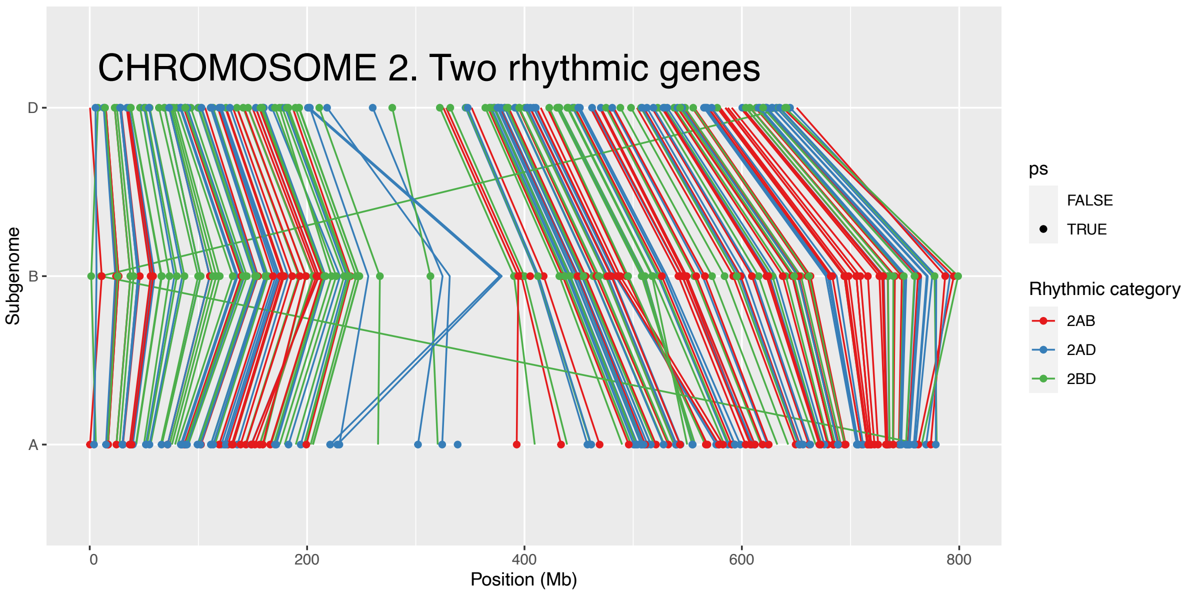

Supplementary Information-Regulation of the circadian transcriptome in a complex polyploid crop

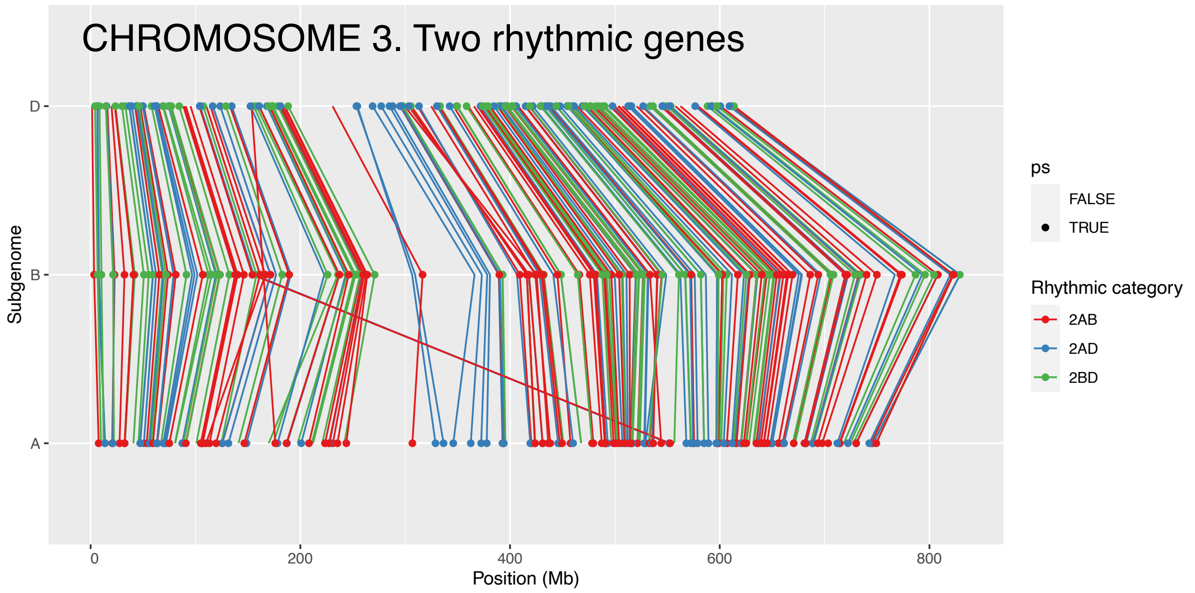


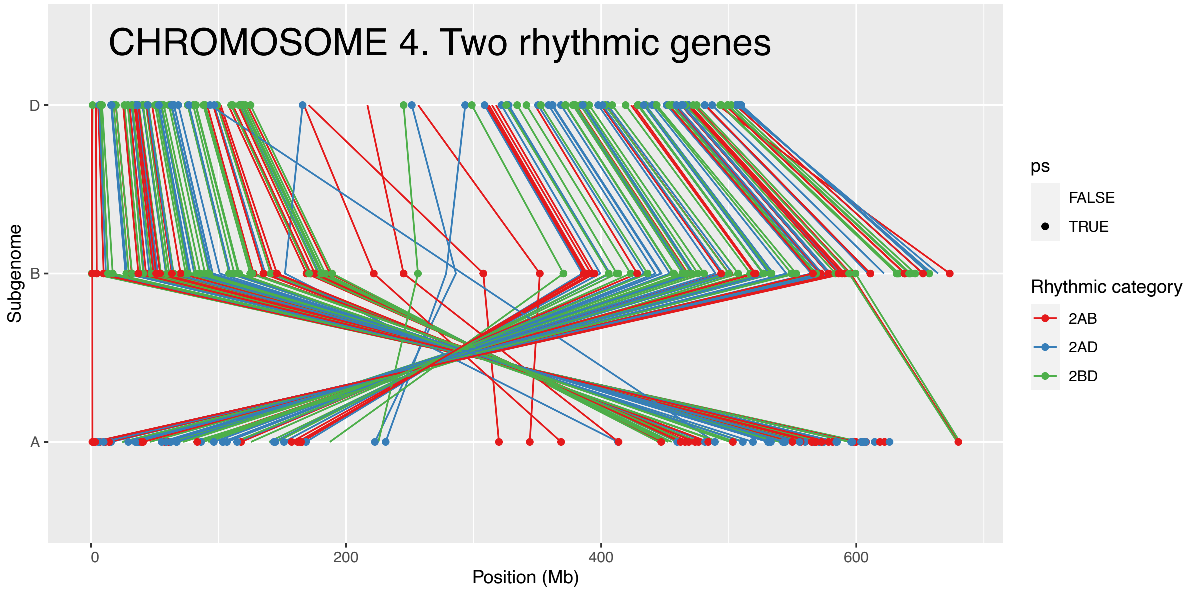


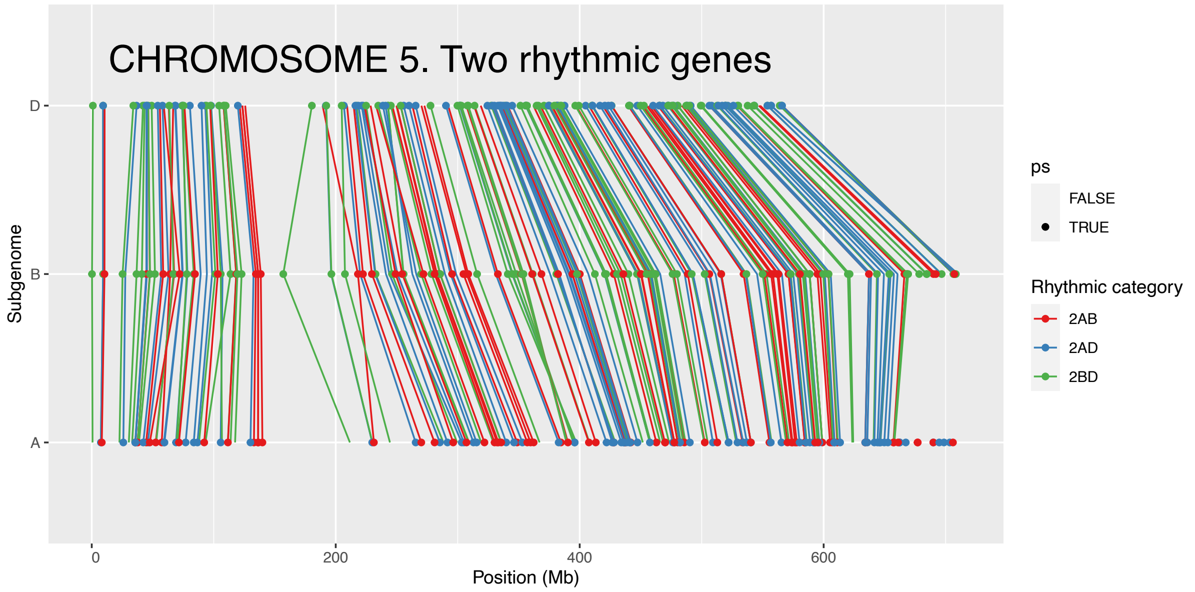

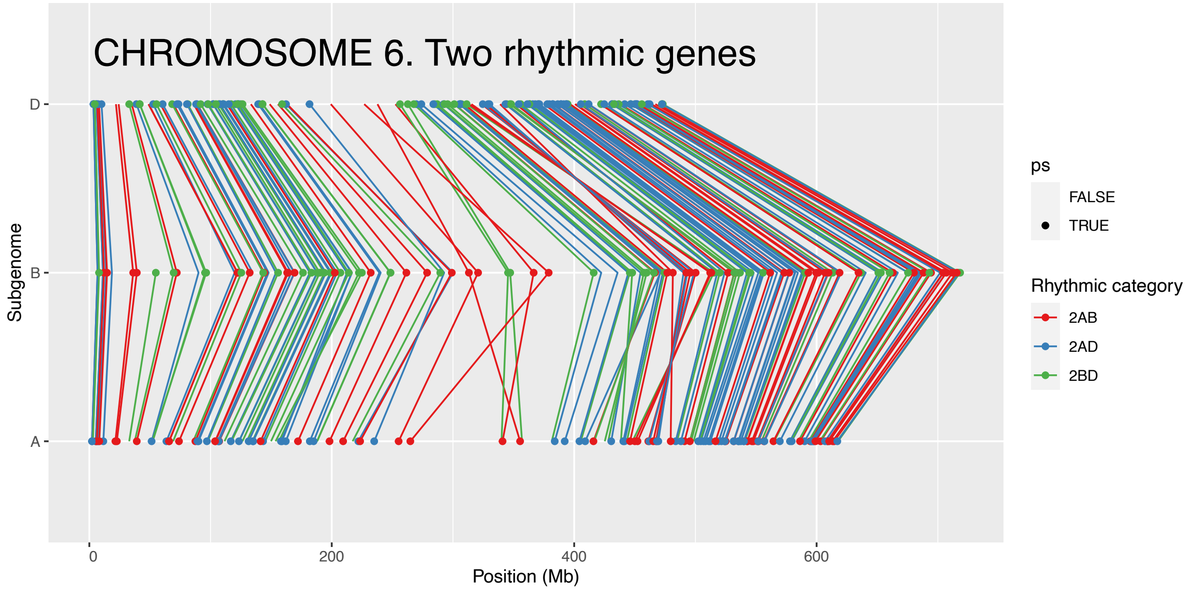
CHROMOSOME 6. Two rhythmic genes
D
B
A
Subgenome
0
200
400
600
Position (Mb)
ps
FALSE
TRUE
Rhythmic category
2AB
2AD
2BD

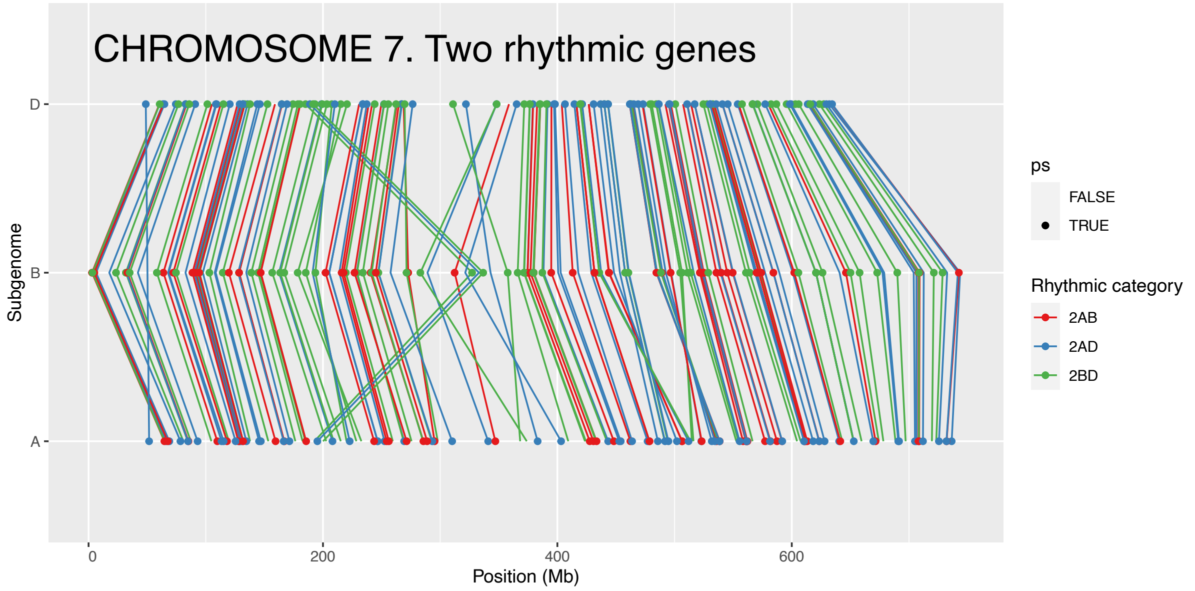
CHROMOSOME 7. Two rhythmic genes
D
B
A
Subgenome
0
200
400
600
Position (Mb)
ps
FALSE
TRUE
Rhythmic category
2AB
2AD
2BD

**Supplementary Figure 8. Loom plots showing positions of runs of imbalanced circadian triads.** Triads in the following categories are shown: 1A, 1B, 1D [triads with one rhythmic gene on the A, B or D chromosomes respectively and two arrhythmic genes], 2AB, 2AD, 2BD [triads with two rhythmic genes on the AB, AD or BD chromosomes respectively and one arrhythmic gene]. Points indicate the rhythmicity of the homoeolog and colored lines represent the category of the triad.

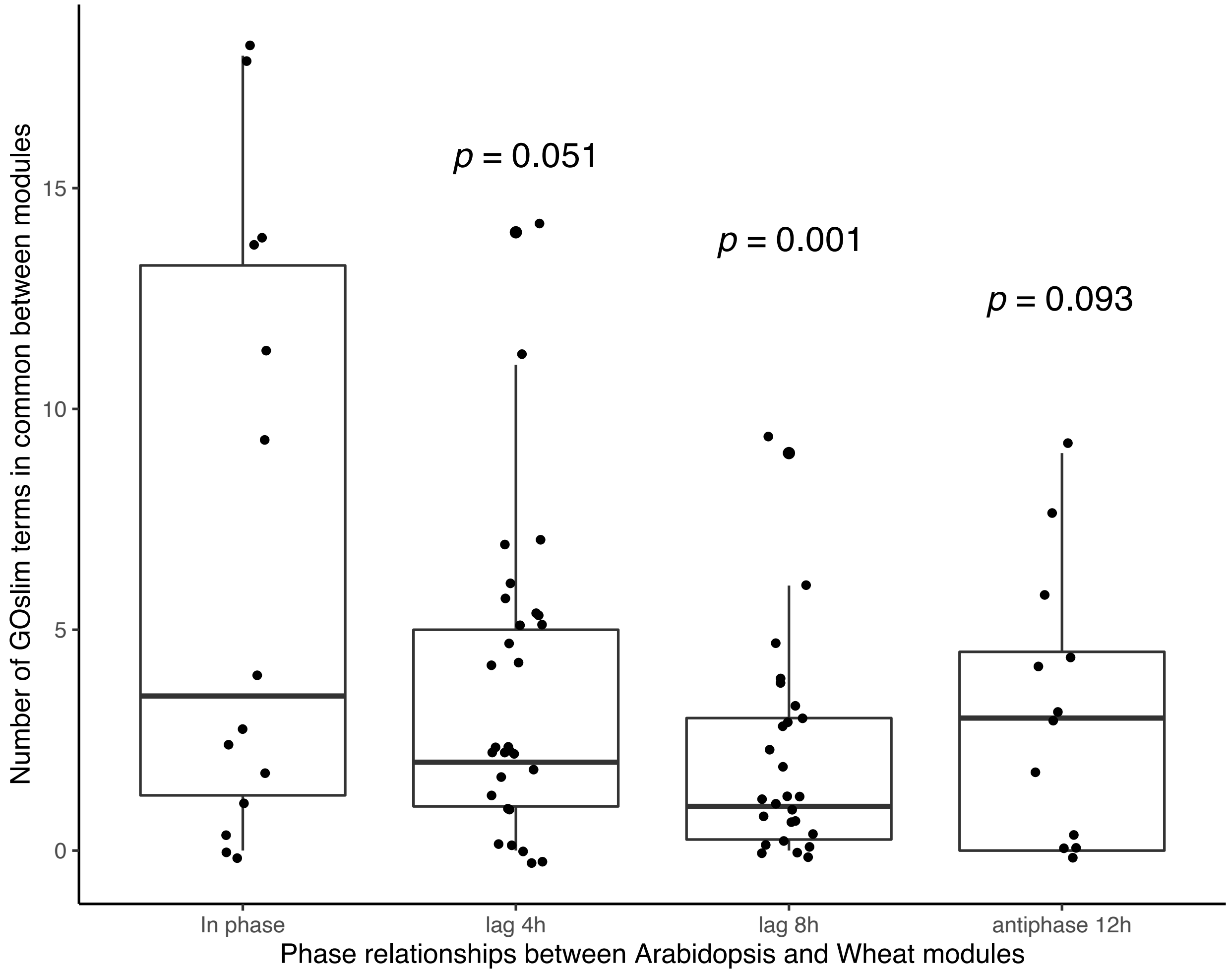


**Supplementary Figure 9. *Arabidopsis* and wheat modules with a peak lag of 0 (synchronous phase) share more GO-slim terms than non-synchronous modules.** Numbers of significantly enriched ($p < 0.05$) GO-slim terms in common between nine pairwise modules in *Arabidopsis* and wheat were counted and the pairwise modules were grouped based on the highest correlation score (peak lag) following cross-correlation with a lag of 0,4, 8 or 12h.

Supplementary Information-Regulation of the circadian transcriptome in a complex polyploid crop

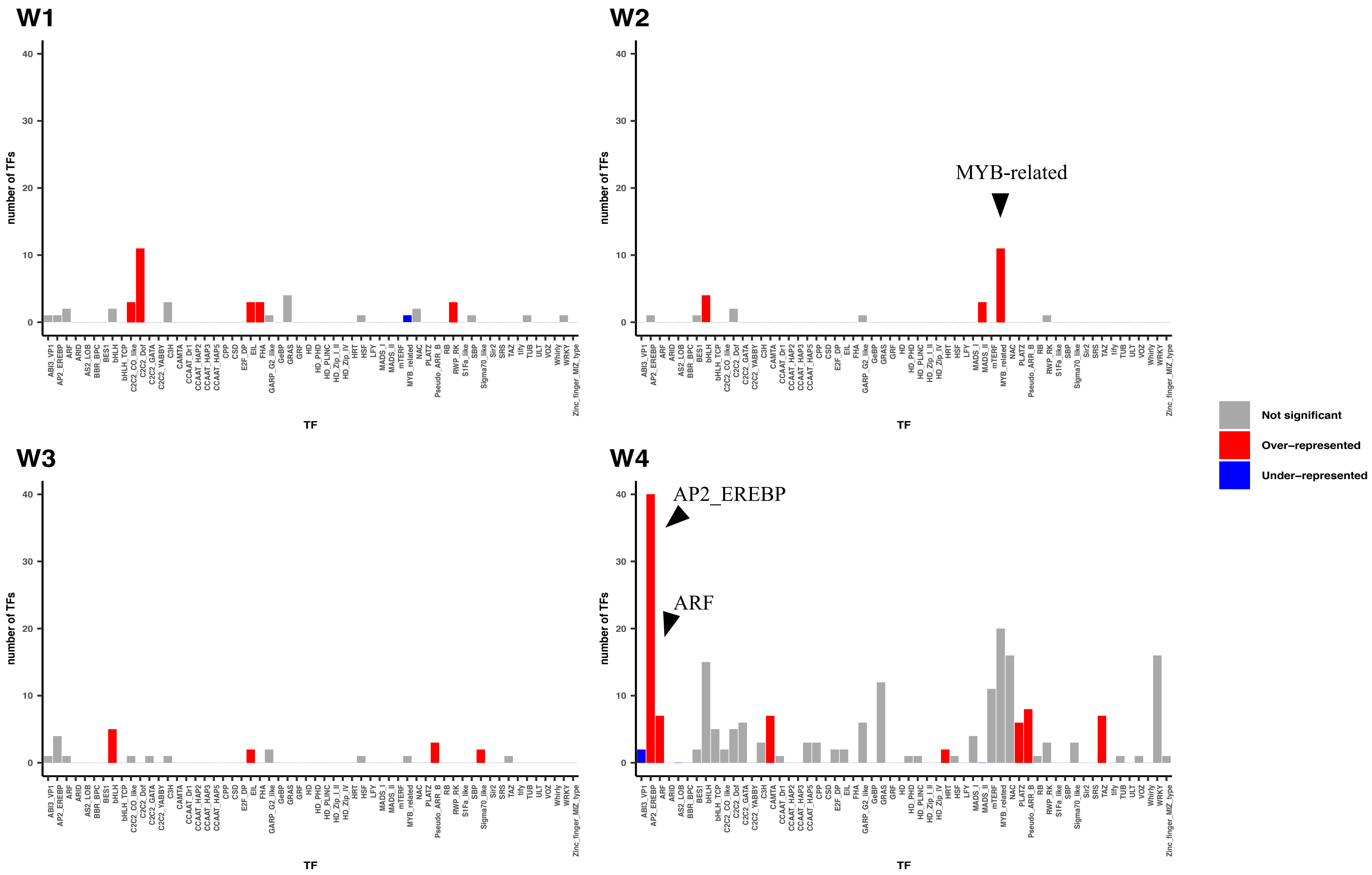

Supplementary Information-Regulation of the circadian transcriptome in a complex polyploid crop

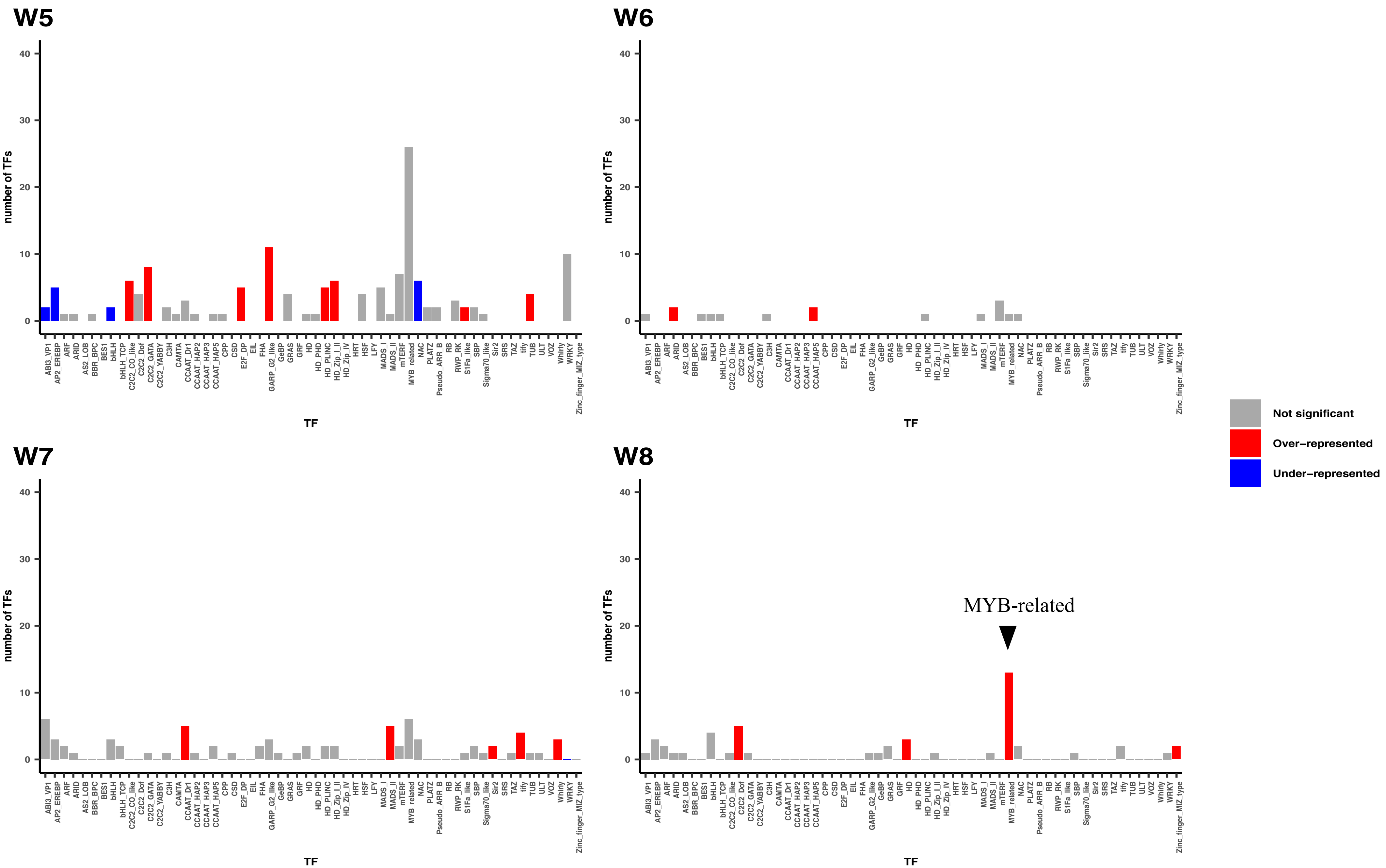

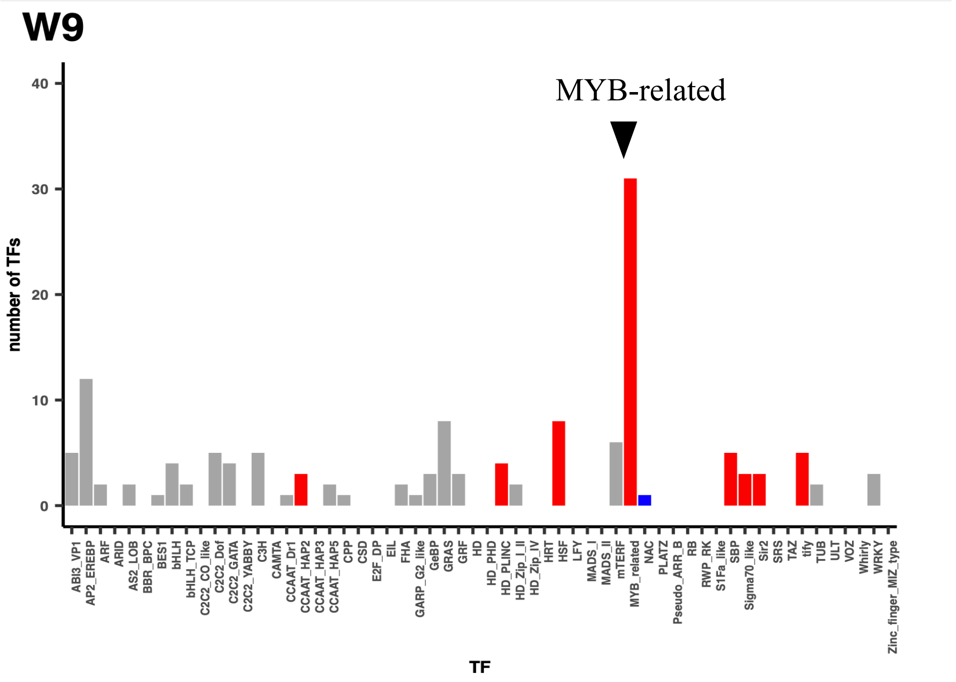


**Supplementary Figure 10. Transcription factor superfamily enrichment in wheat modules.**

Barcharts showing number of genes (y axis) belonging to each TF superfamily (x axis) within wheat co-expression modules W1-W9. Coloured bars denote those superfamilies significantly over-enriched (red) or under-enriched (blue) compared to the total number of TF superfamilies present in the 16,327 genes submitted to WGCNA (Fisher's exact test, p <=0.05). Families not significantly over- or under-enriched are coloured grey.

Supplementary Information-Regulation of the circadian transcriptome in a complex polyploid crop

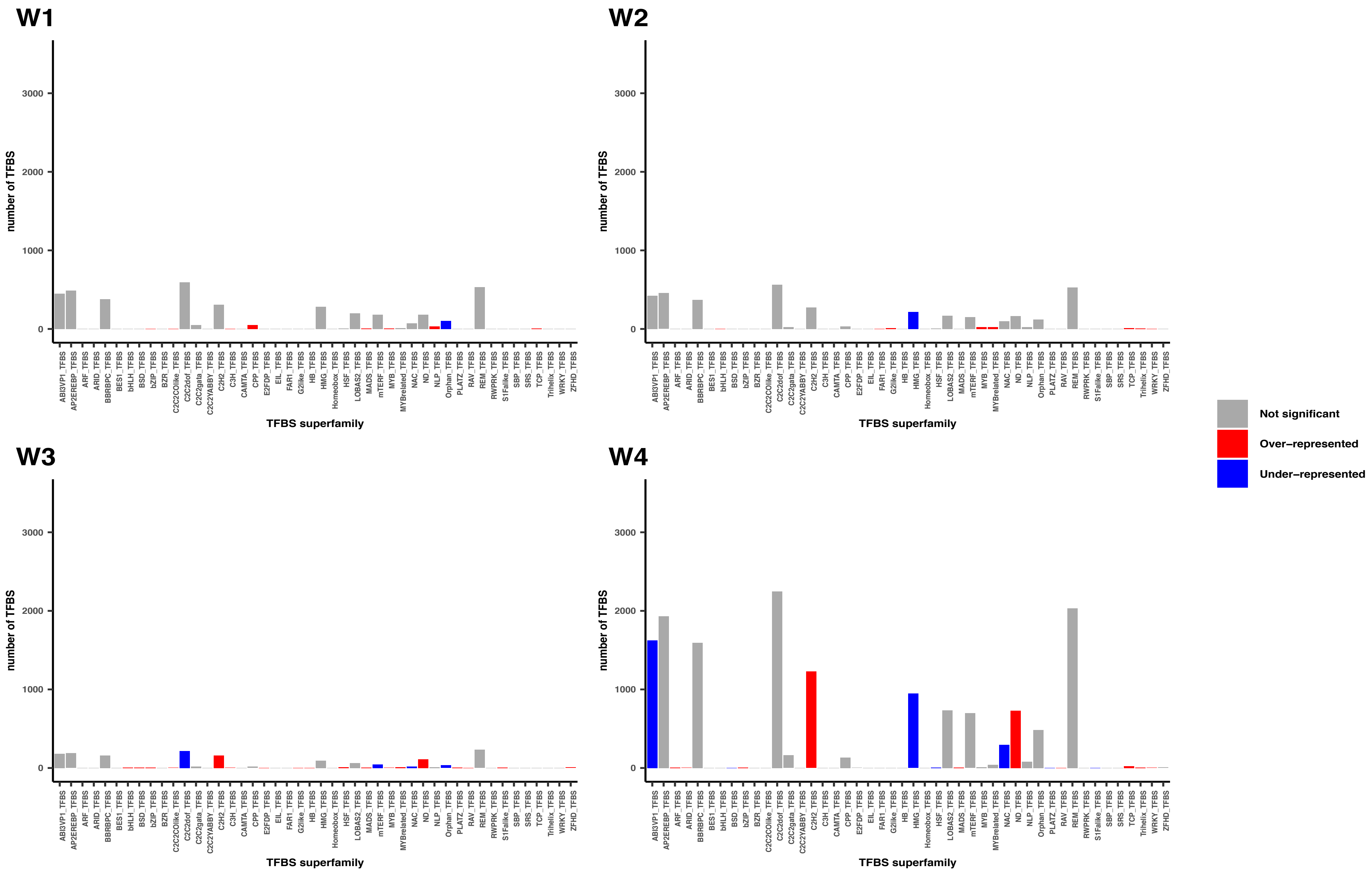

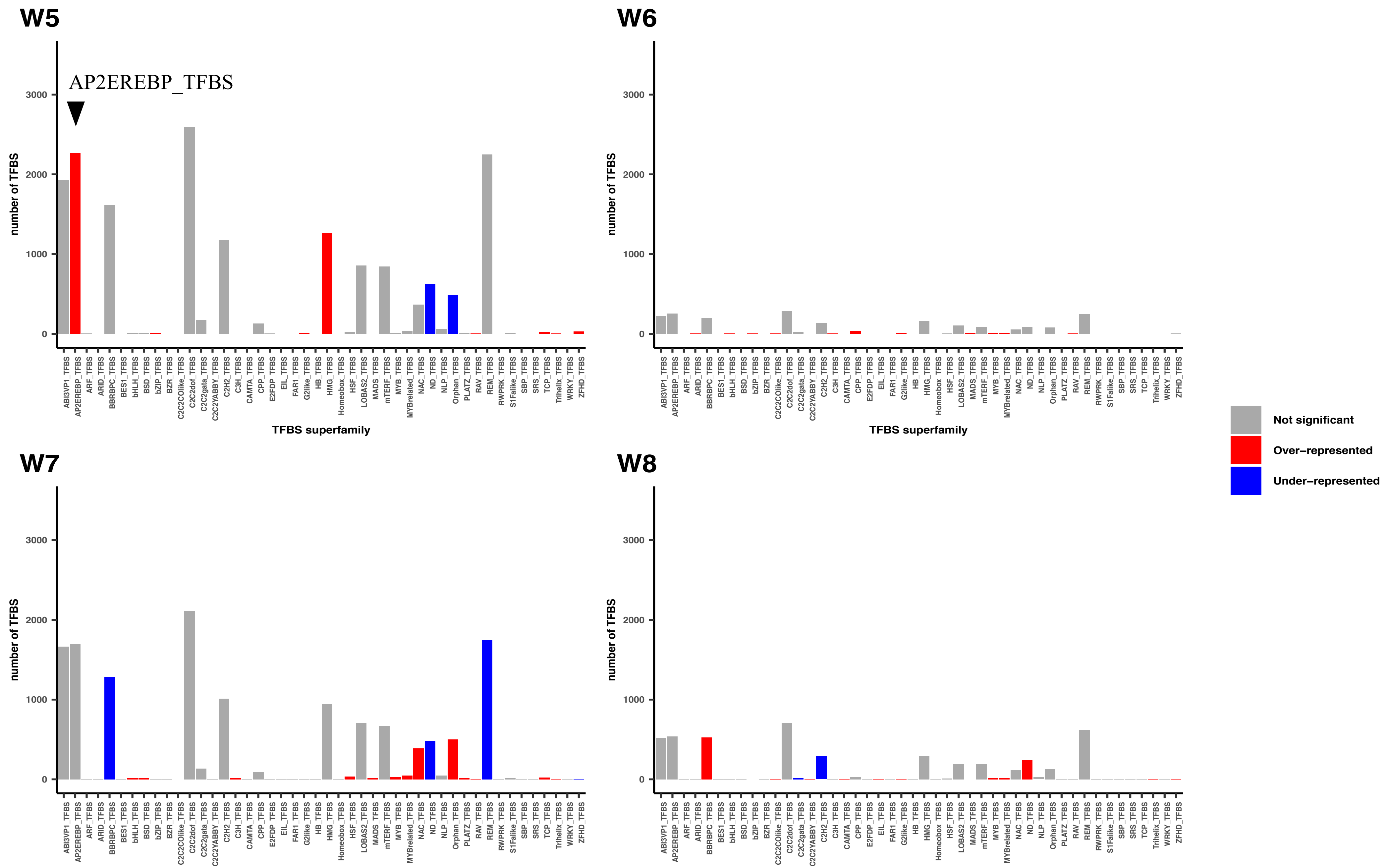
W5
AP2EREBP_TFBS
W6
W7
W8
number of TFBS
0
1000
2000
3000
TFBS superfamily
ABI3VP1_TFBS
AP2EREBP_TFBS
ARF_TFBS
ARID_TFBS
BBRBPC_TFBS
BES1_TFBS
bHLH_TFBS
BSD_TFBS
bZIP_TFBS
BZR_TFBS
C2C2COlike_TFBS
C2C2dof_TFBS
C2C2gata_TFBS
C2C2YABBY_TFBS
C2H2_TFBS
C3H_TFBS
CAMTA_TFBS
CPP_TFBS
E2FDP_TFBS
EIL_TFBS
FAR1_TFBS
G2like_TFBS
HB_TFBS
HMG_TFBS
Homeobox_TFBS
HSF_TFBS
LOBAS2_TFBS
MADS_TFBS
mTERF_TFBS
MYB_TFBS
MYBrelated_TFBS
NAC_TFBS
ND_TFBS
NLP_TFBS
Orphan_TFBS
PLATZ_TFBS
RAV_TFBS
REM_TFBS
RWPRK_TFBS
S1Falike_TFBS
SBP_TFBS
SRS_TFBS
TCP_TFBS
Trihelix_TFBS
WRKY_TFBS
ZFHD_TFBS
Not significant
Over−represented
Under−represented

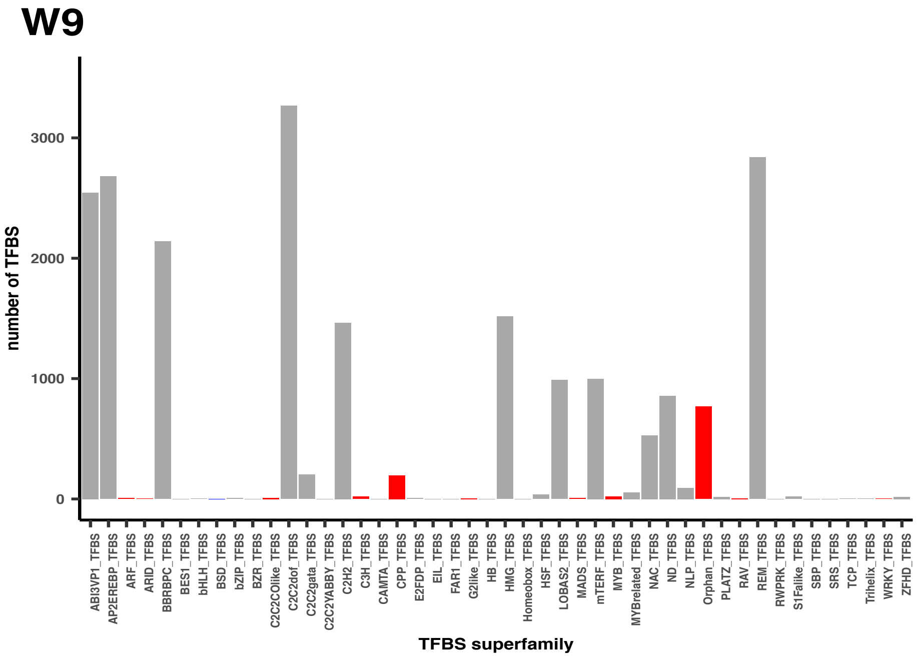


**Supplementary Figure 11. Transcription factor binding site (TFBS) superfamily enrichment in wheat modules.**

Bar charts showing number of non-redundant TFBS motifs (y axis) belonging to each TFBS superfamily (x axis) within wheat co-expression modules W1-W9. Coloured bars denote those superfamilies significantly over-enriched (red) or under-enriched (blue) compared to the total number of non-redundant TFBS motif superfamilies present in the 16,327 genes submitted to WGCNA (Fisher's exact test, $p <= 0.05$). Families not significantly over- or under-enriched are coloured grey.

#### Tree 1: MYB (LHY-like)

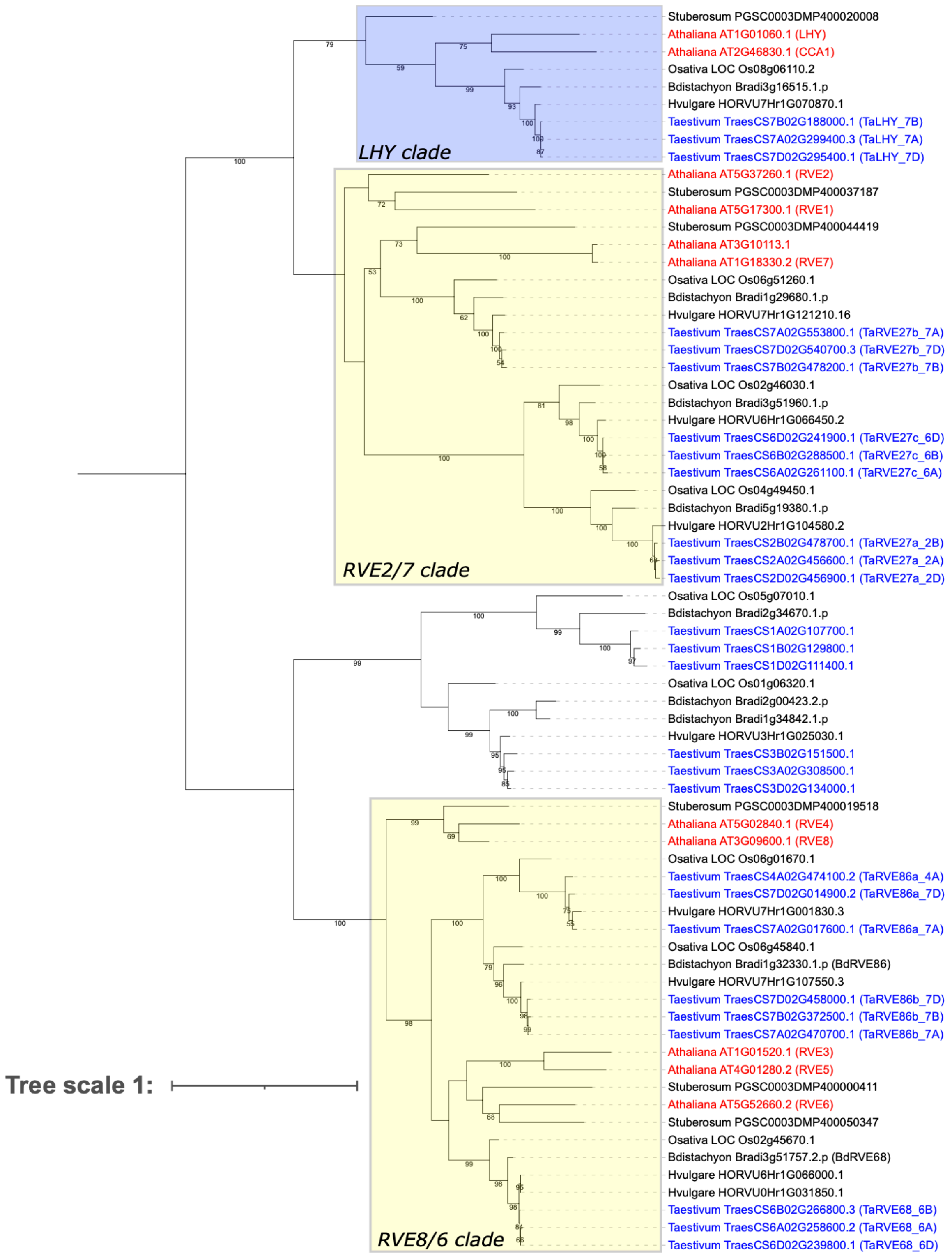

**Supplementary Figure 12. Phylogenetic relationships for circadian genes based on alignment of LHY-like MYB full length proteins.** The species identifiers for each species have been abbreviated as follows; Taestivum: *Triticum aestivum* (hexaploid wheat), Bdistachyon: *Brachypodium distachyon,* Hvulgare: *Hordeum vulgare* (barley), Osativa: *Oryza sativa* (rice), Stuberosum: *Solanum tuberosum* (potato), Athaliana: *Arabidopsis thaliana*. Wheat genes have been highlighted in blue and *Arabidopsis* genes in red for clarity. Bootstrap values are calculated using RAxML, with values over 50 shown on branches.

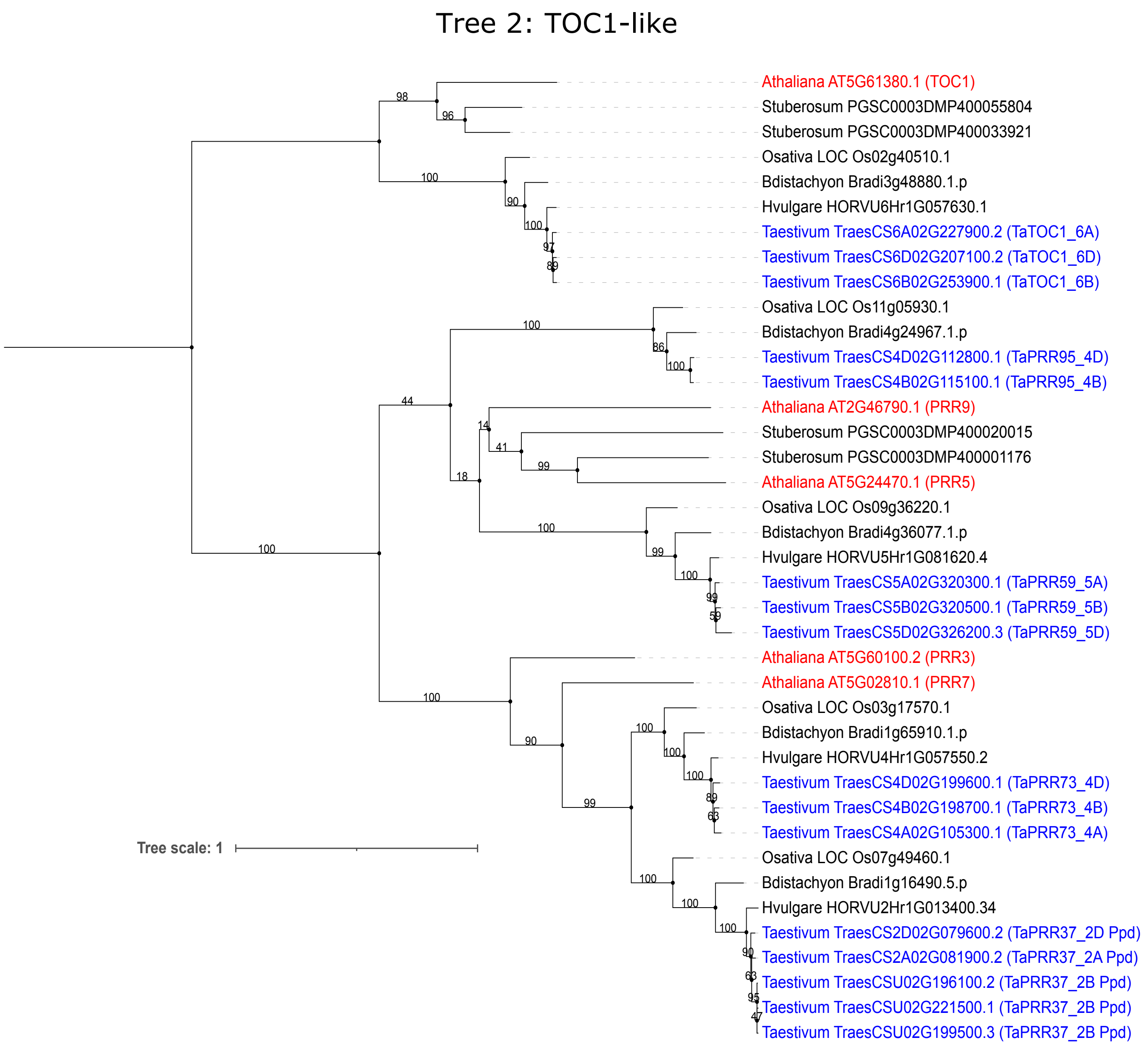


**Supplementary Figure 13. Phylogenetic relationships for circadian genes based on alignment of TOC1-like and PRR full length proteins.** The species identifiers for each species have been abbreviated as follows; Taestivum: *Triticum aestivum* (hexaploid wheat), Bdistachyon: *Brachypodium distachyon,* Hvulgare: *Hordeum vulgare* (barley), Osativa: *Oryza sativa* (rice), Stuberosum: *Solanum tuberosum* (potato), Athaliana: *Arabidopsis*

*thaliana*. Wheat genes have been highlighted in blue and *Arabidopsis* genes in red for clarity. Bootstrap values are calculated using RAxML, with values over 50 shown on branches.

### Tree 3: PAS/LOV (ZTL-like)

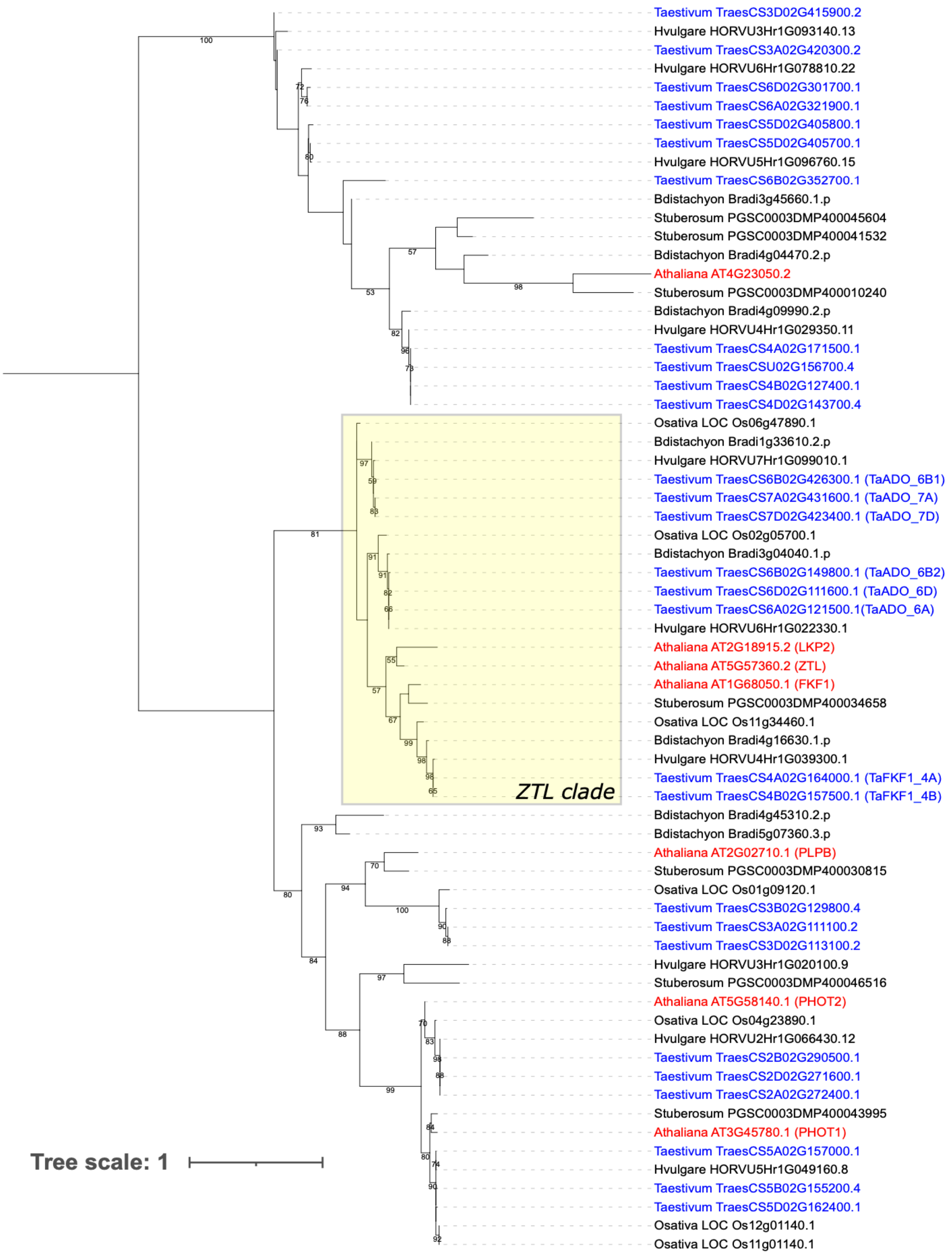

**Supplementary Figure 14. Phylogenetic relationships for circadian genes based on alignment of PAS/LOV protein domains.** The species identifiers for each species have been abbreviated as follows; Taestivum: *Triticum aestivum* (hexaploid wheat), Bdistachyon: *Brachypodium distachyon,* Hvulgare: *Hordeum vulgare* (barley), Osativa: *Oryza sativa* (rice), Stuberosum: *Solanum tuberosum* (potato), Athaliana: *Arabidopsis thaliana*. Wheat genes have been highlighted in blue and *Arabidopsis* genes in red for clarity. Bootstrap values are calculated using RAxML, with values over 50 shown on branches.

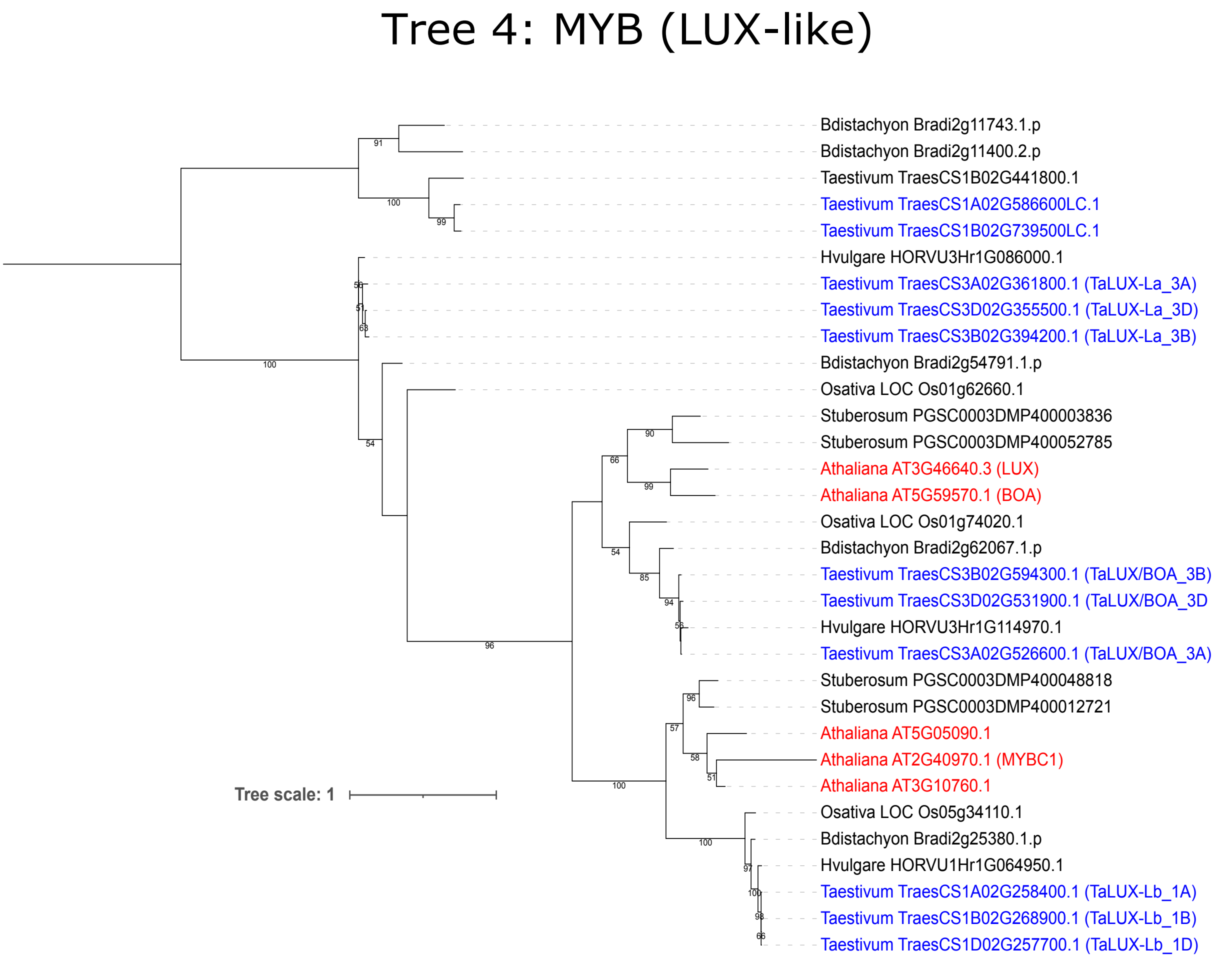


**Supplementary Figure 15. Phylogenetic relationships for circadian genes based on alignment of LUX-like MYB full length proteins.** The species identifiers for each species have been abbreviated as follows; Taestivum: *Triticum aestivum* (hexaploid wheat), Bdistachyon: *Brachypodium distachyon,* Hvulgare: *Hordeum vulgare* (barley), Osativa: *Oryza sativa* (rice), Stuberosum: *Solanum tuberosum* (potato), Athaliana: *Arabidopsis thaliana*. Wheat genes have been highlighted in blue and *Arabidopsis* genes in red for clarity. Bootstrap values are calculated using RAxML, with values over 50 shown on branches.

### Tree 5: ELF4-like

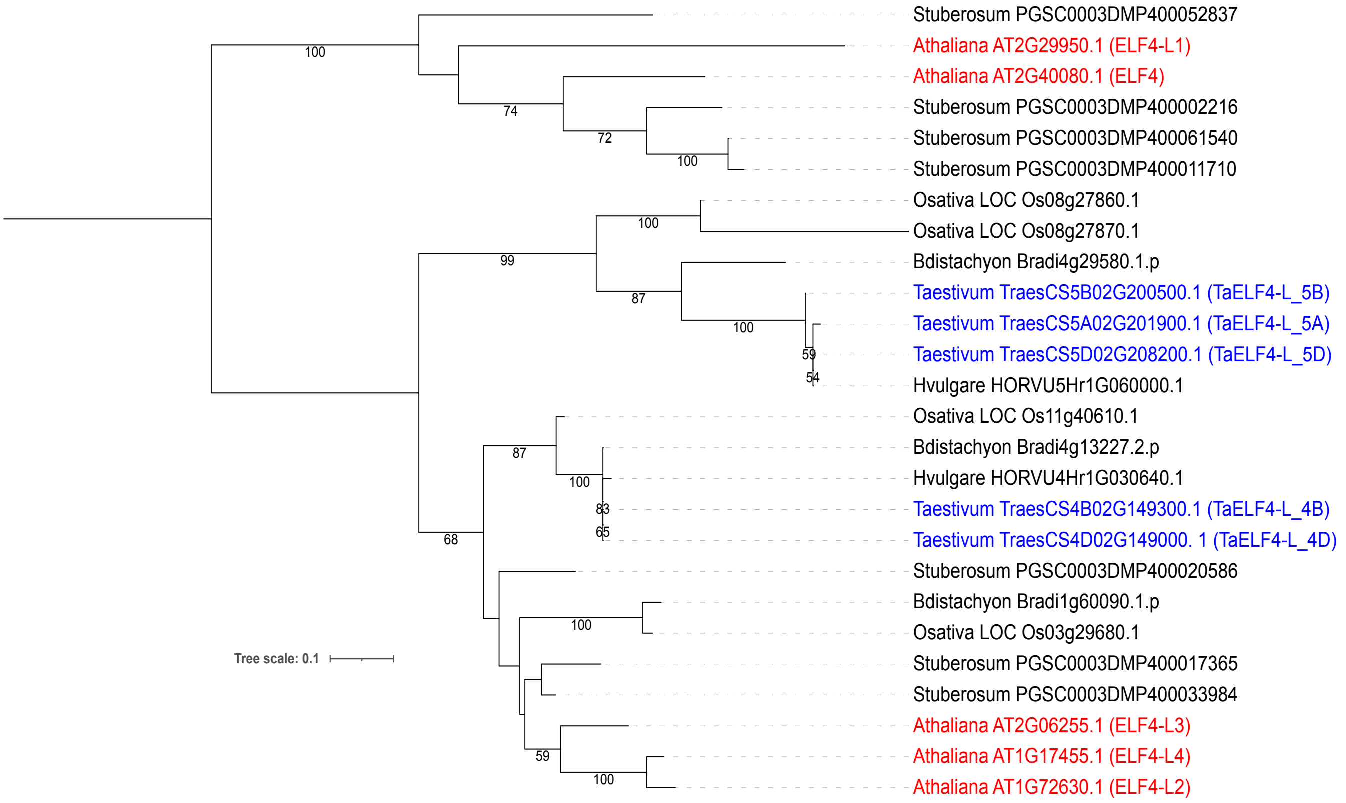


**Supplementary Figure 16. Phylogenetic relationships for circadian genes based on alignment of ELF4-like protein domains.** The species identifiers for each species have been abbreviated as follows; Taestivum: *Triticum aestivum* (hexaploid wheat), Bdistachyon: *Brachypodium distachyon,* Hvulgare: *Hordeum vulgare* (barley), Osativa: *Oryza sativa* (rice), Stuberosum: *Solanum tuberosum* (potato), Athaliana: *Arabidopsis thaliana*. Wheat genes have been highlighted in blue and *Arabidopsis* genes in red for clarity. Bootstrap values are calculated using RAxML, with values over 50 shown on branches.

Supplementary Information-Regulation of the circadian transcriptome in a complex polyploid crop

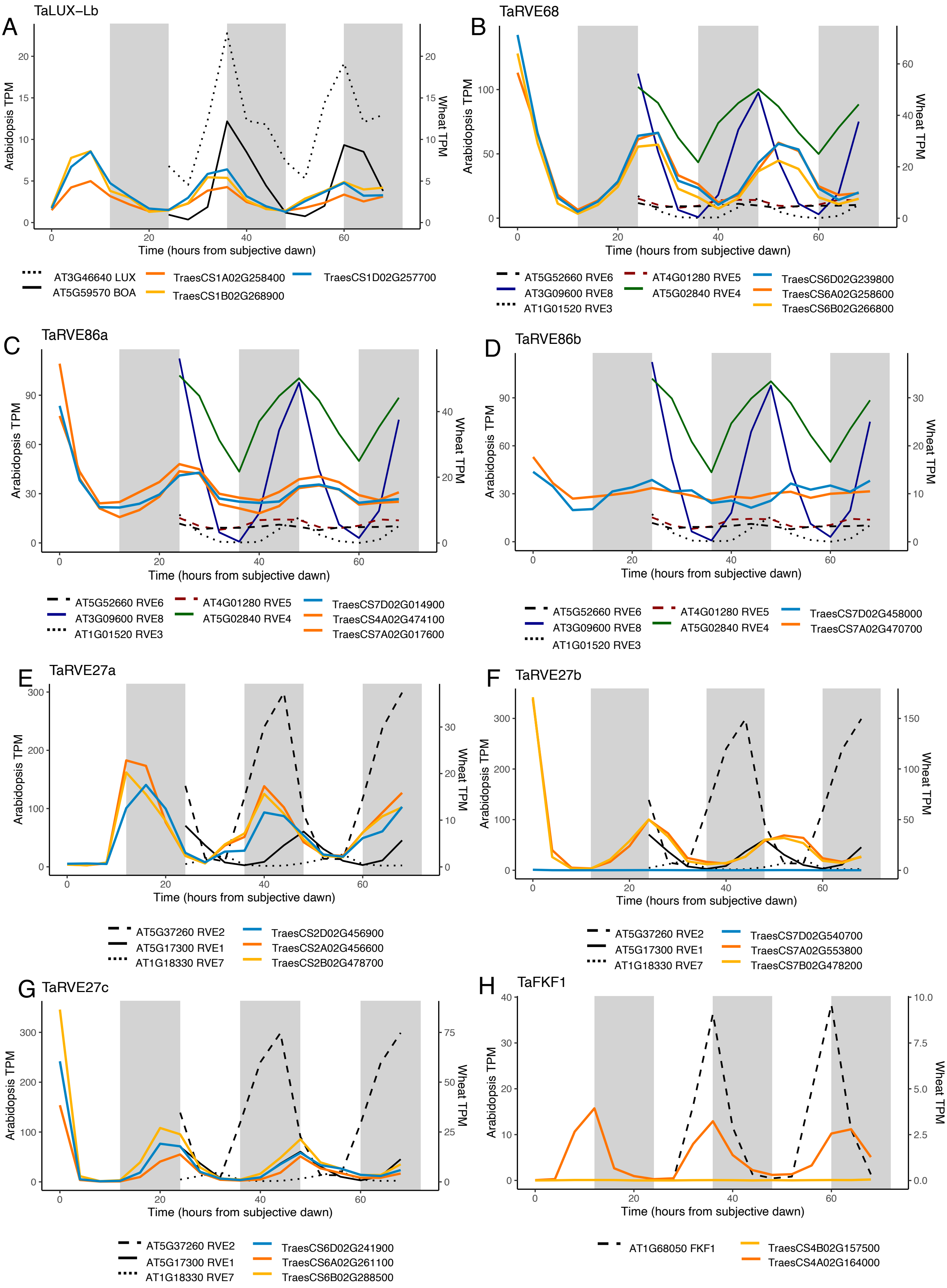

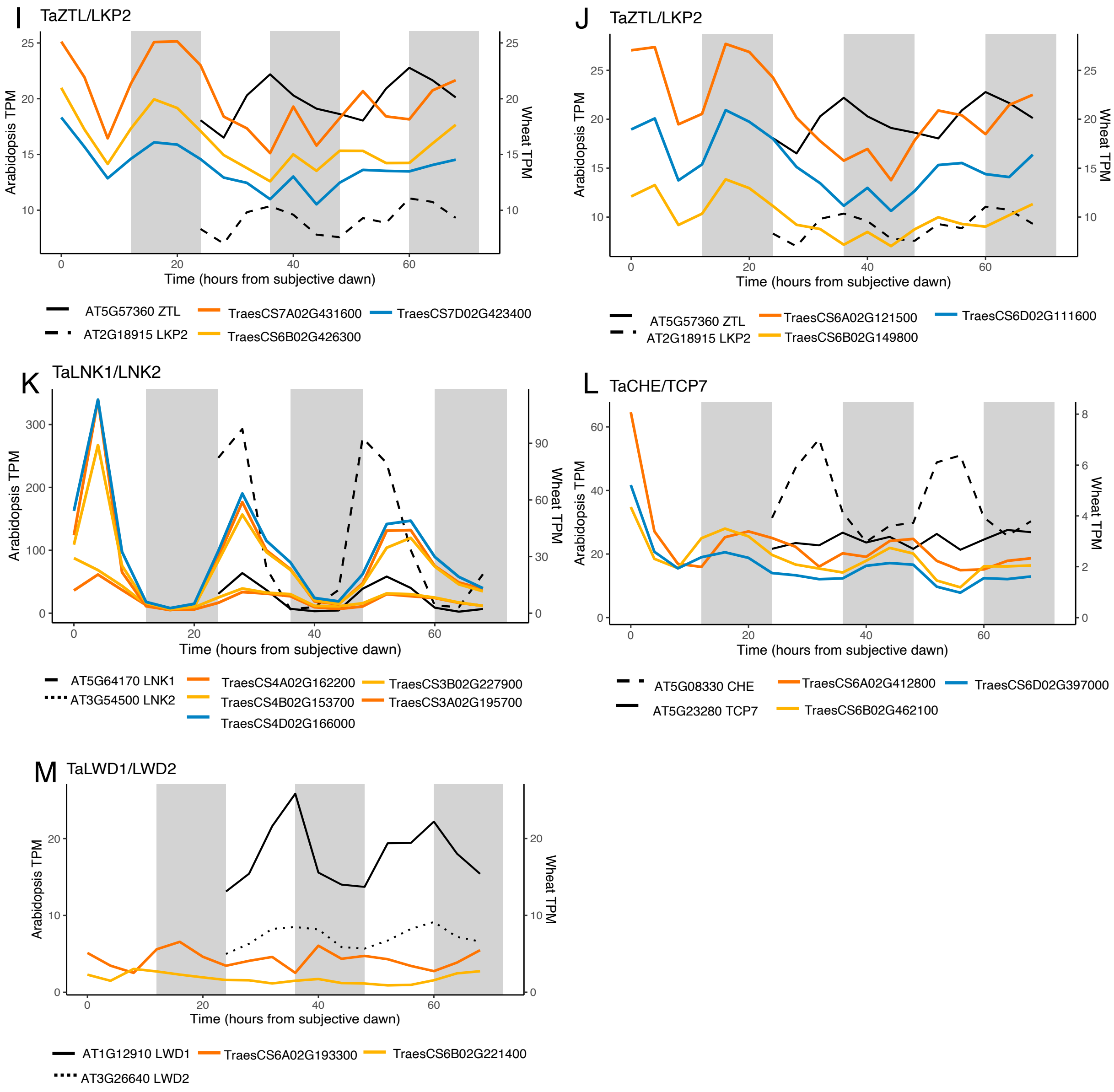


**Supplementary Figure 17. Additional free-running expression of core circadian clock genes in wheat and their homologs in *Arabidopsis*.** Supplementary to Figure 3 in the main text. Wheat circadian clock genes were identified through alignment of phylogenetic protein family trees or BLASTP to known clock gene homologs. Wheat homoeologs are coloured according to their identity to either the A genome (orange), B genome (yellow) or D genome (blue) and grey and white blocks indicate subjective dark and light time periods under constant conditions. Data represent the mean of three biological replicates and transcript expression is collapsed to gene level.

Supplementary Information-Regulation of the circadian transcriptome in a complex polyploid crop

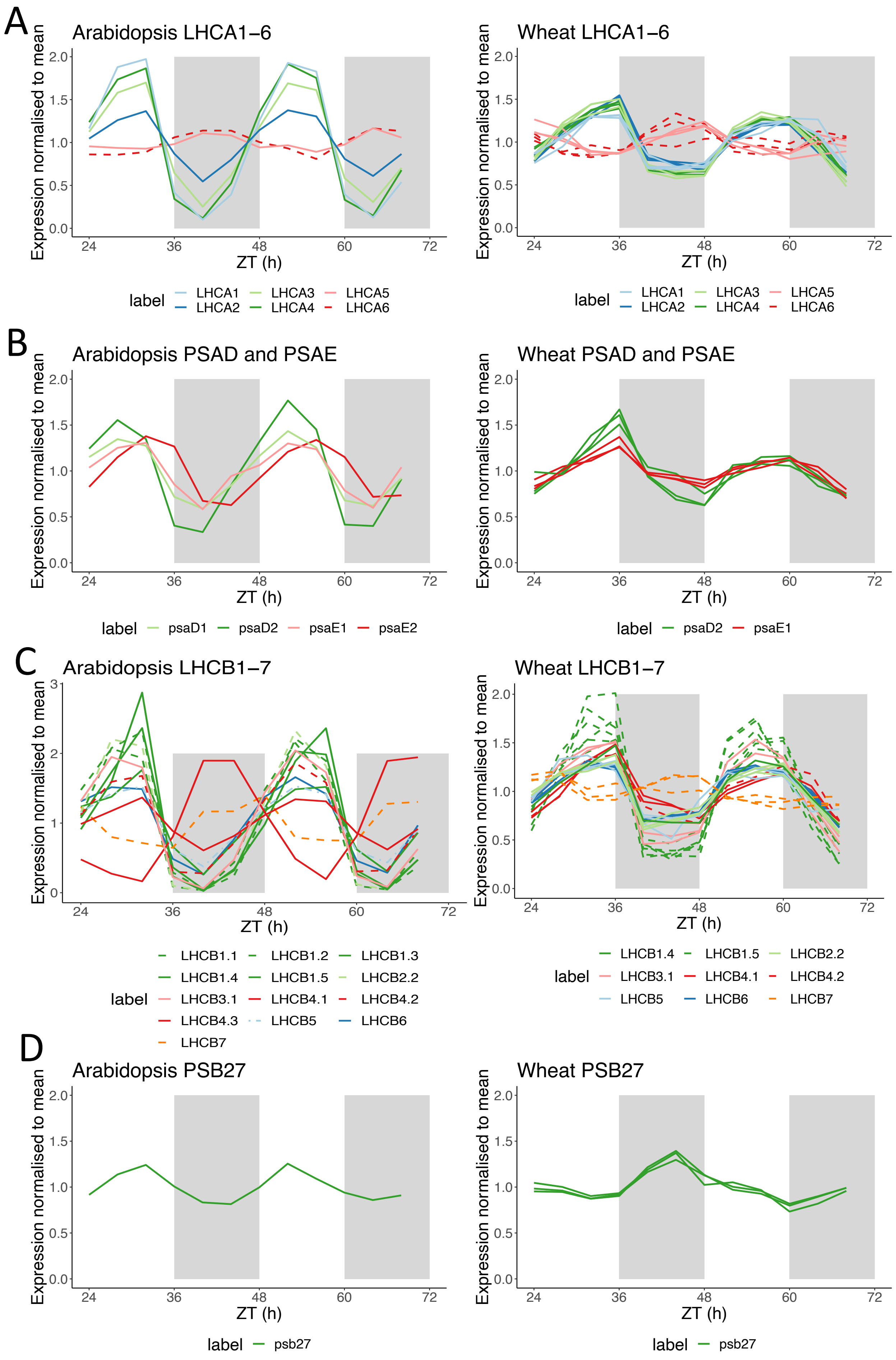

**Supplementary Figure 18. Arabidopsis and Wheat photosystem genes under free-running L:L conditions.** Data is mean normalised TPM. Shaded white and grey backgrounds indicate perceived day and night periods respectively. Gene IDs for all genes plotted can be seen in Supplementary Table 8.

Supplementary Information-Regulation of the circadian transcriptome in a complex polyploid crop

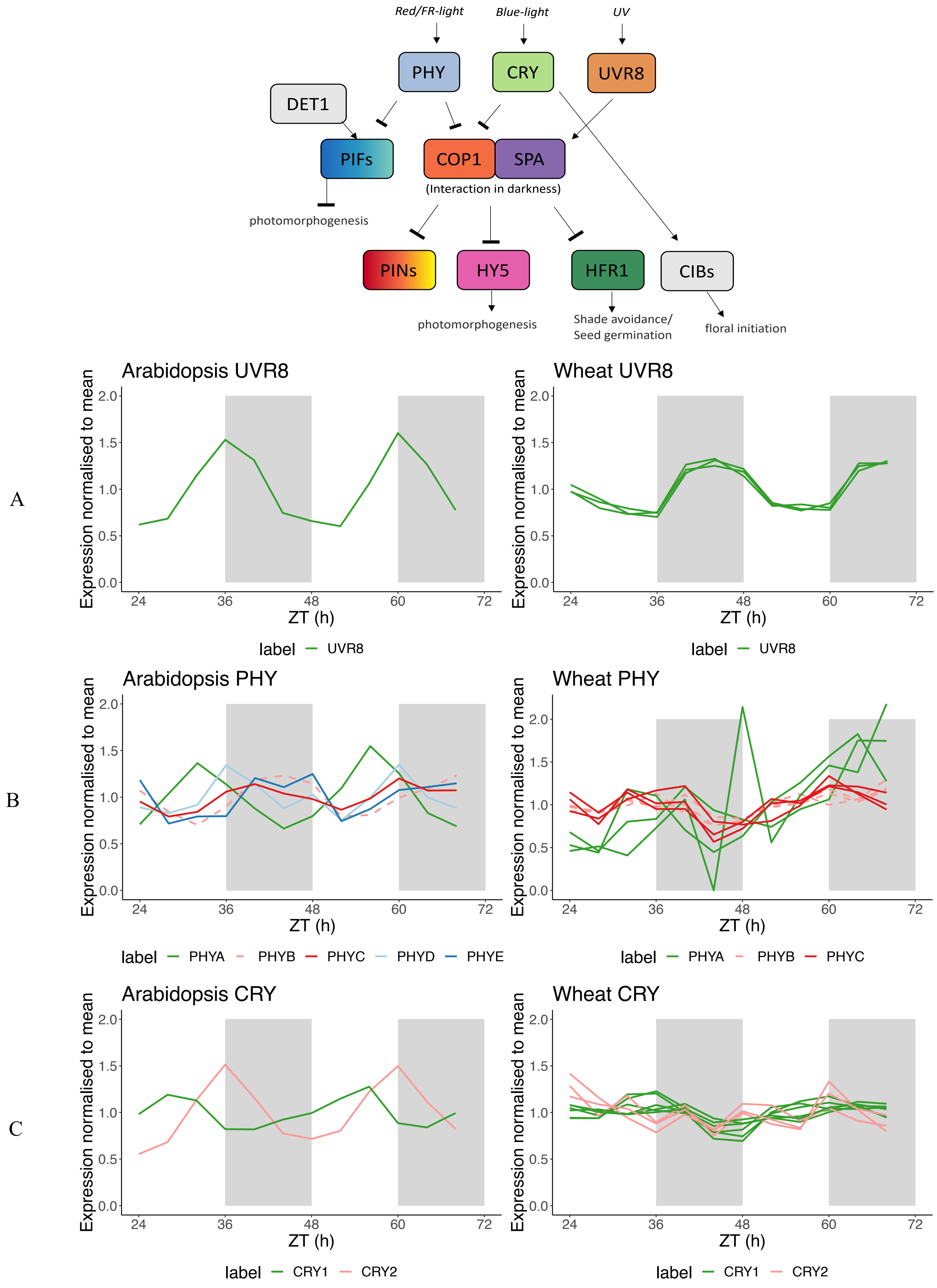

D

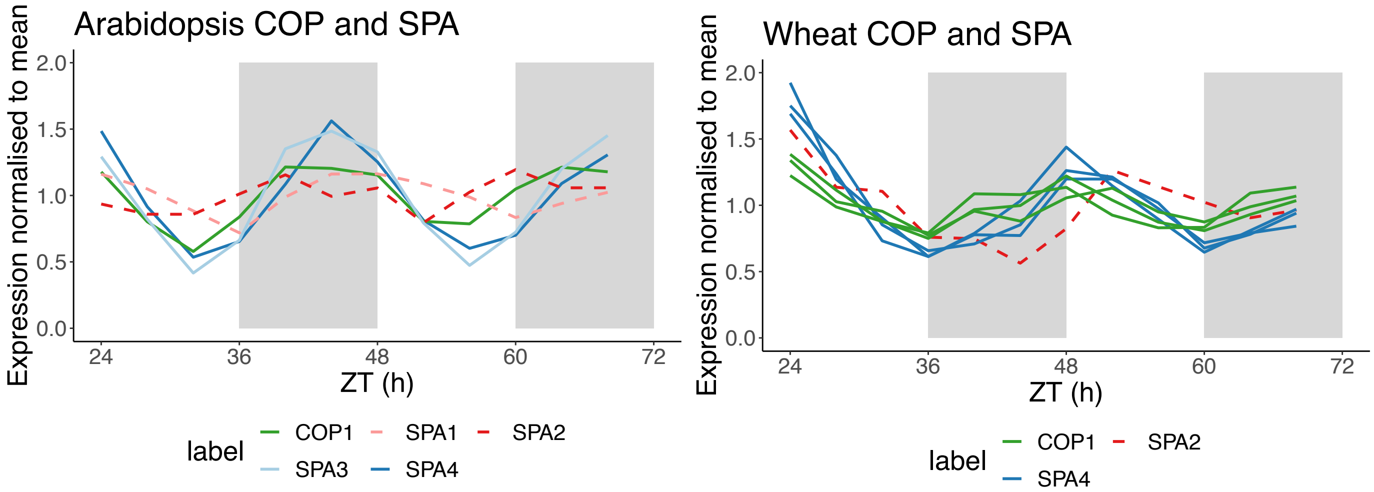
Arabidopsis COP and SPA
Expression normalised to mean
ZT (h)
label
COP1
SPA1
SPA2
SPA3
SPA4
Wheat COP and SPA
Expression normalised to mean
ZT (h)
label
COP1
SPA2
SPA4

E

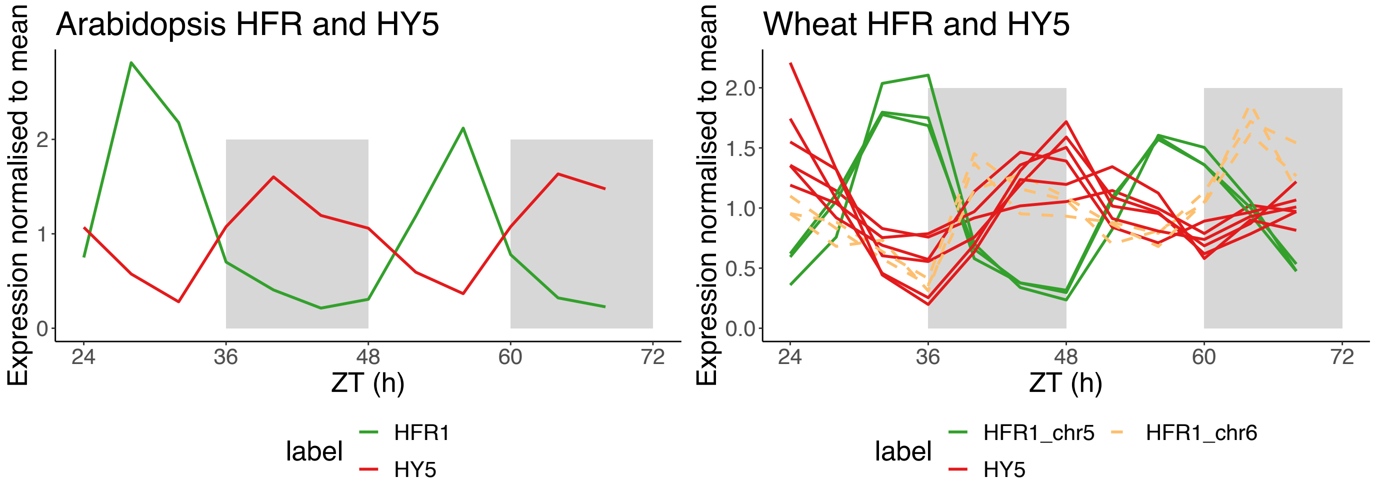
Arabidopsis HFR and HY5
Expression normalised to mean
ZT (h)
label
HFR1
HY5
Wheat HFR and HY5
Expression normalised to mean
ZT (h)
label
HFR1_chr5
HFR1_chr6
HY5

F

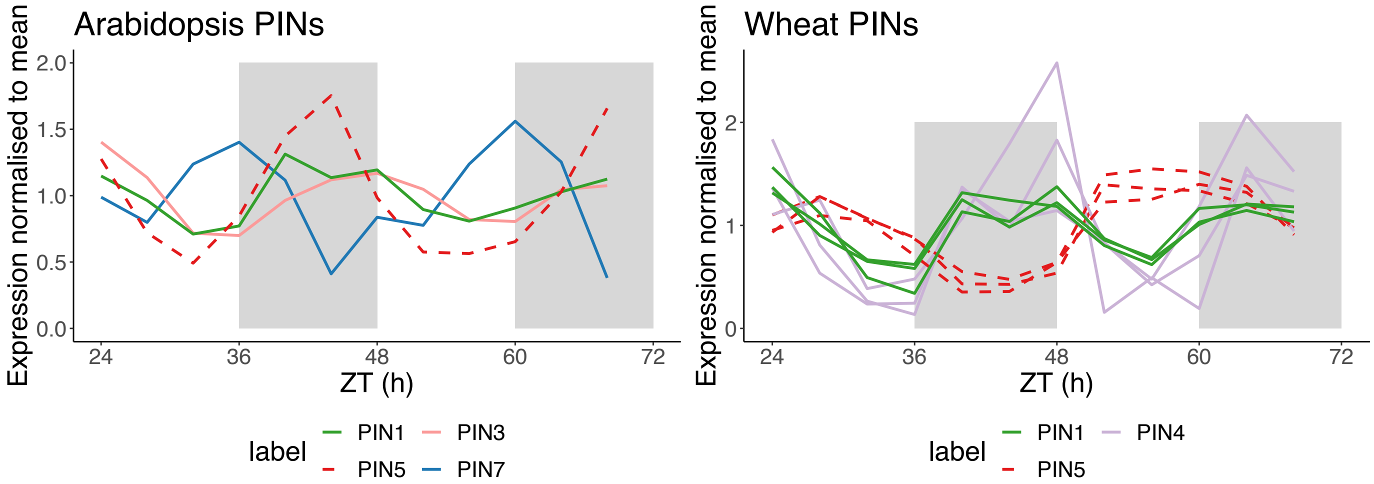
Arabidopsis PINs
Expression normalised to mean
ZT (h)
label
PIN1
PIN3
PIN5
PIN7
Wheat PINs
Expression normalised to mean
ZT (h)
label
PIN1
PIN4
PIN5

G

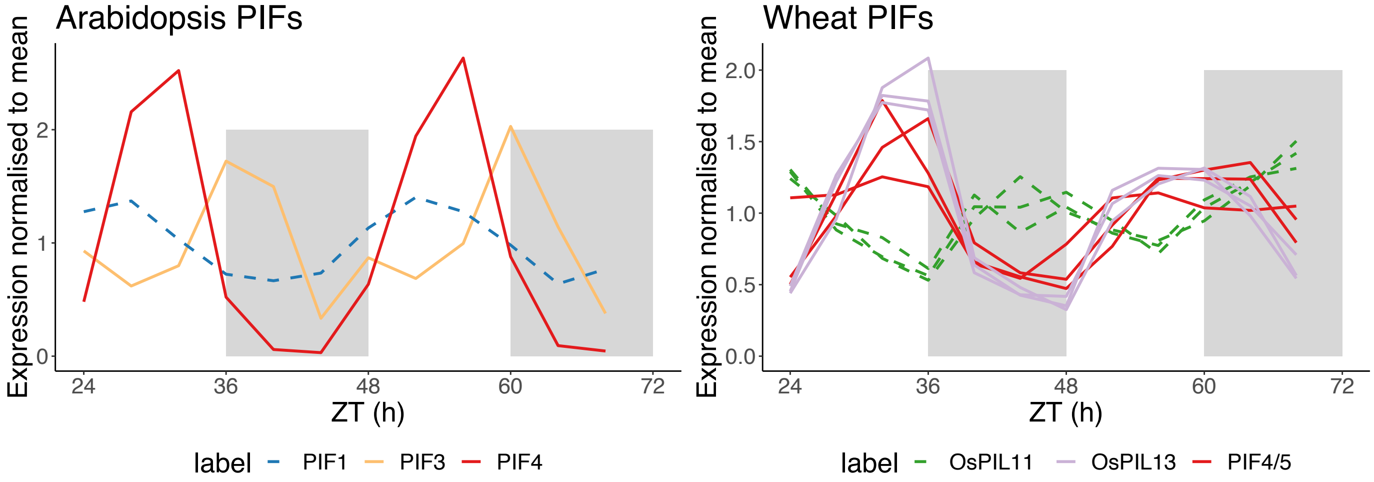
Arabidopsis PIFs
Expression normalised to mean
ZT (h)
label
PIF1
PIF3
PIF4
Wheat PIFs
Expression normalised to mean
ZT (h)
label
OsPIL11
OsPIL13
PIF4/5

H

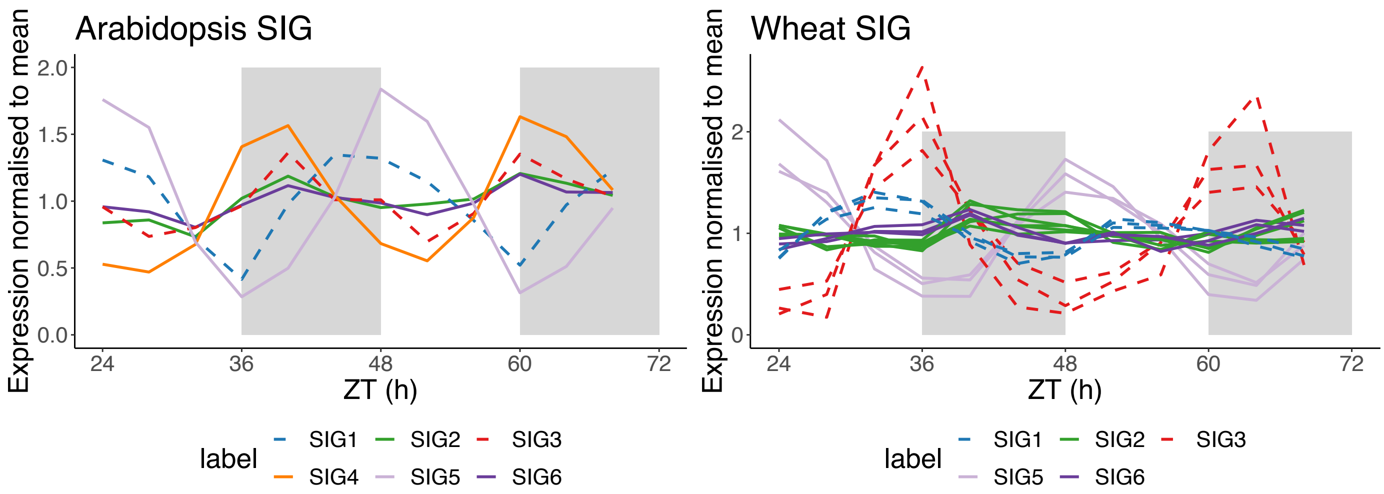


**Supplementary Figure 19. *Arabidopsis* and wheat light signalling genes under free-running L:L conditions.** Data is mean normalised TPM. Shaded white and grey backgrounds indicate perceived day and night periods respectively. Gene IDs for all genes plotted can be seen in Supplementary Table 8.

Supplementary Information-Regulation of the circadian transcriptome in a complex polyploid crop

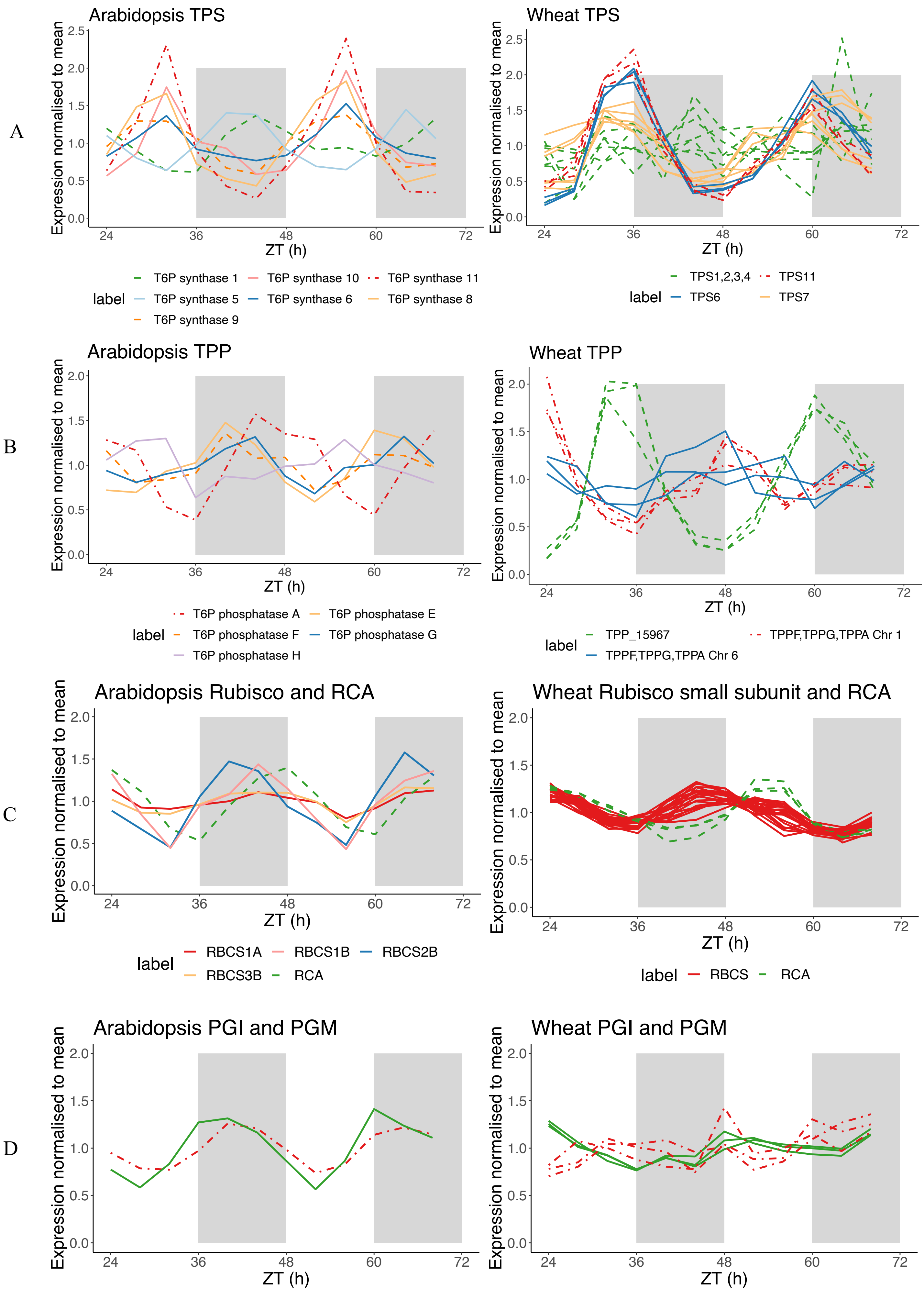

E

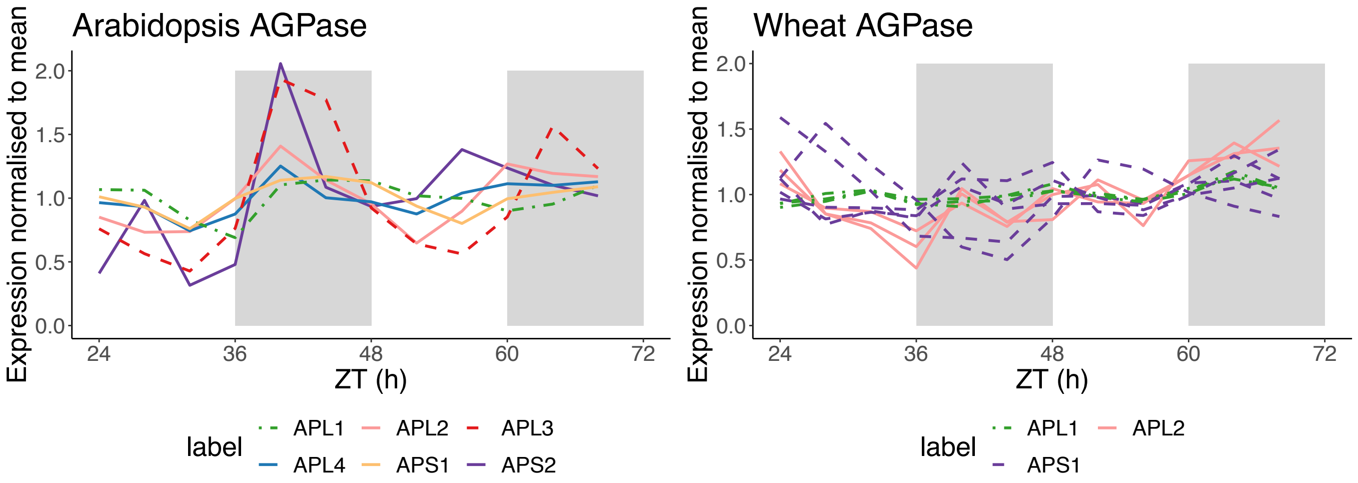
Arabidopsis AGPase
Wheat AGPase
Expression normalised to mean
ZT (h)
label
APL1
APL2
APL3
APL4
APS1
APS2

F

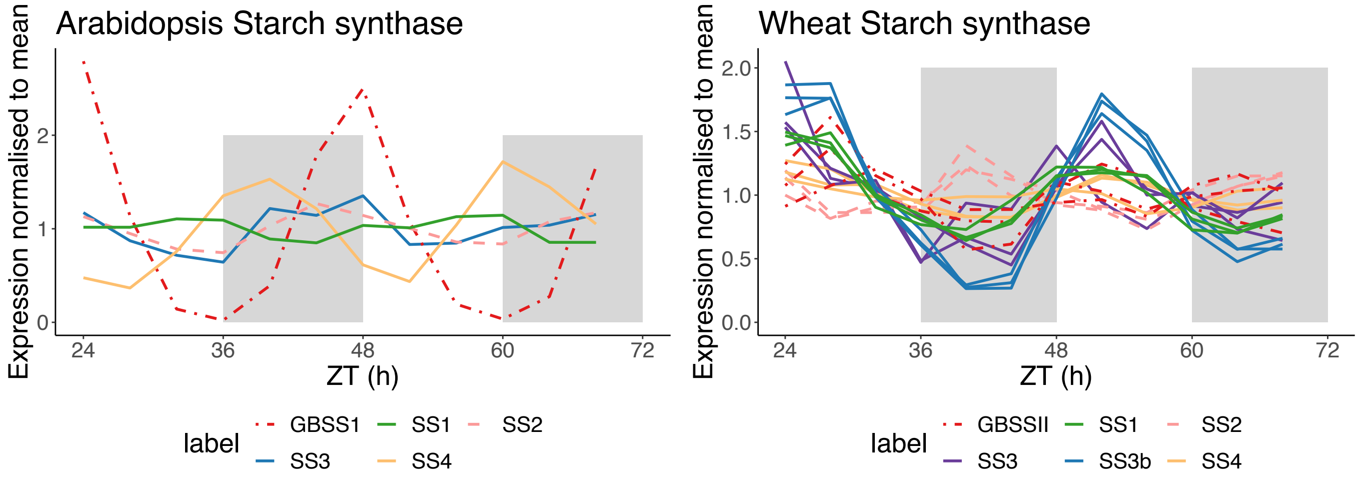
Arabidopsis Starch synthase
Wheat Starch synthase
Expression normalised to mean
ZT (h)
label
GBSS1
SS1
SS2
SS3
SS4
GBSSII
SS3b

G

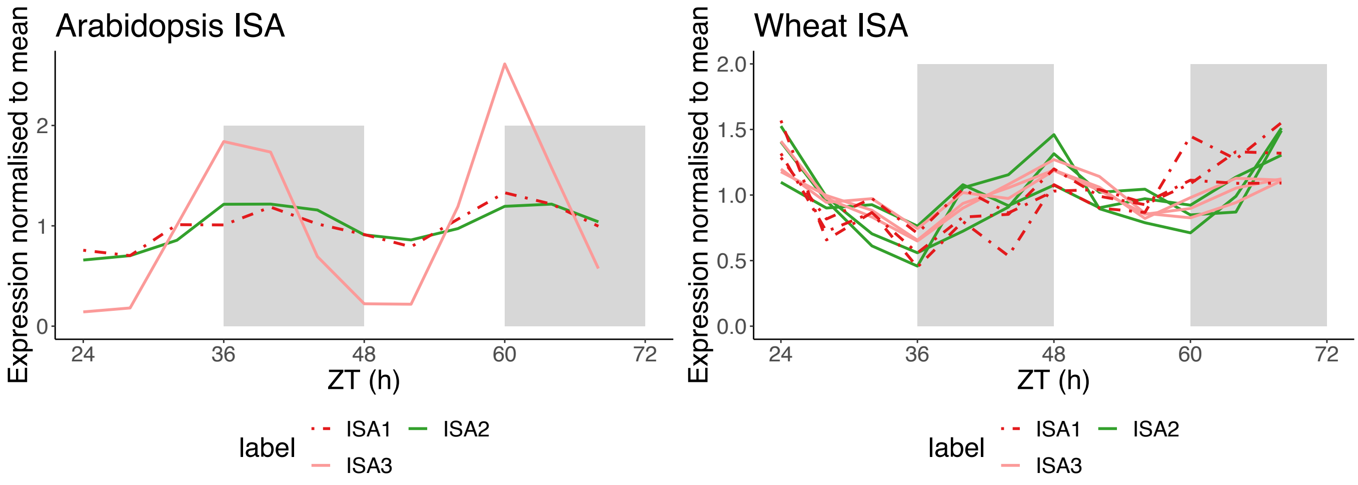
Arabidopsis ISA
Wheat ISA
Expression normalised to mean
ZT (h)
label
ISA1
ISA2
ISA3

H

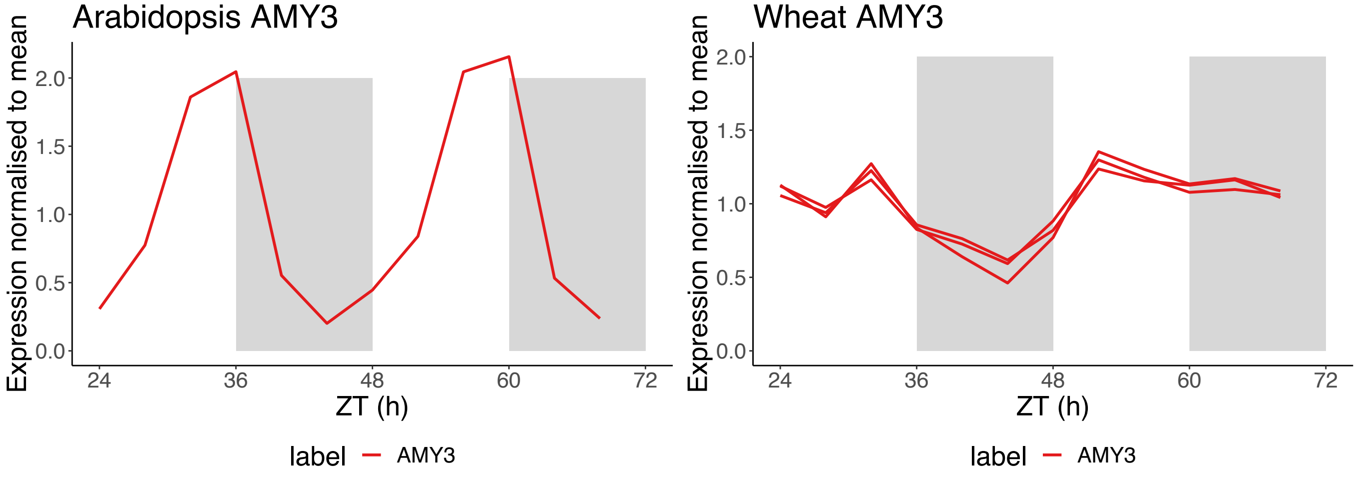
Arabidopsis AMY3
Wheat AMY3
Expression normalised to mean
ZT (h)
label
AMY3

I

J

K

L

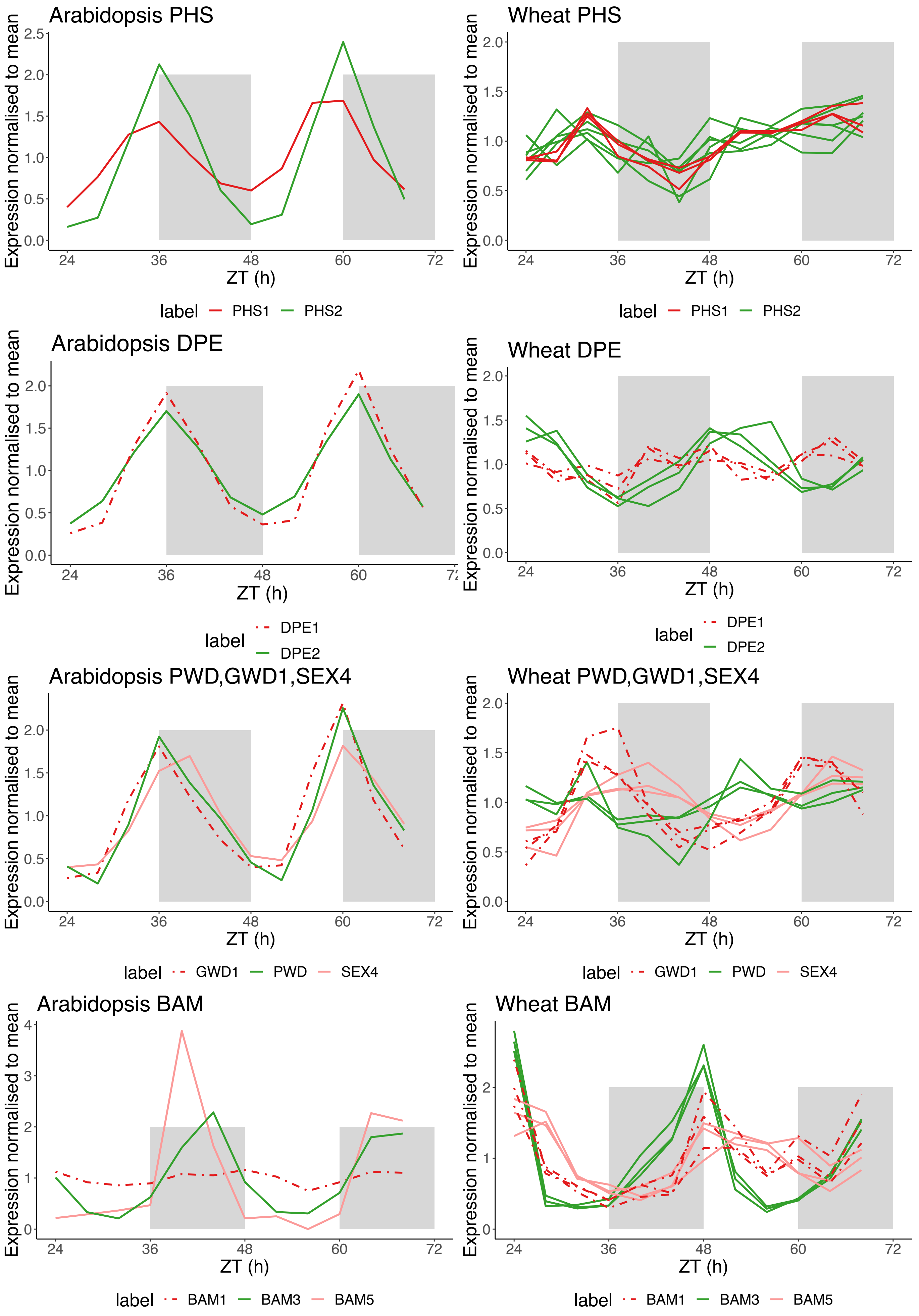
Arabidopsis PHS
Wheat PHS
Expression normalised to mean
ZT (h)
label PHS1 PHS2
Arabidopsis DPE
Wheat DPE
label DPE1 DPE2
Arabidopsis PWD,GWD1,SEX4
Wheat PWD,GWD1,SEX4
label GWD1 PWD SEX4
Arabidopsis BAM
Wheat BAM
label BAM1 BAM3 BAM5

M

N

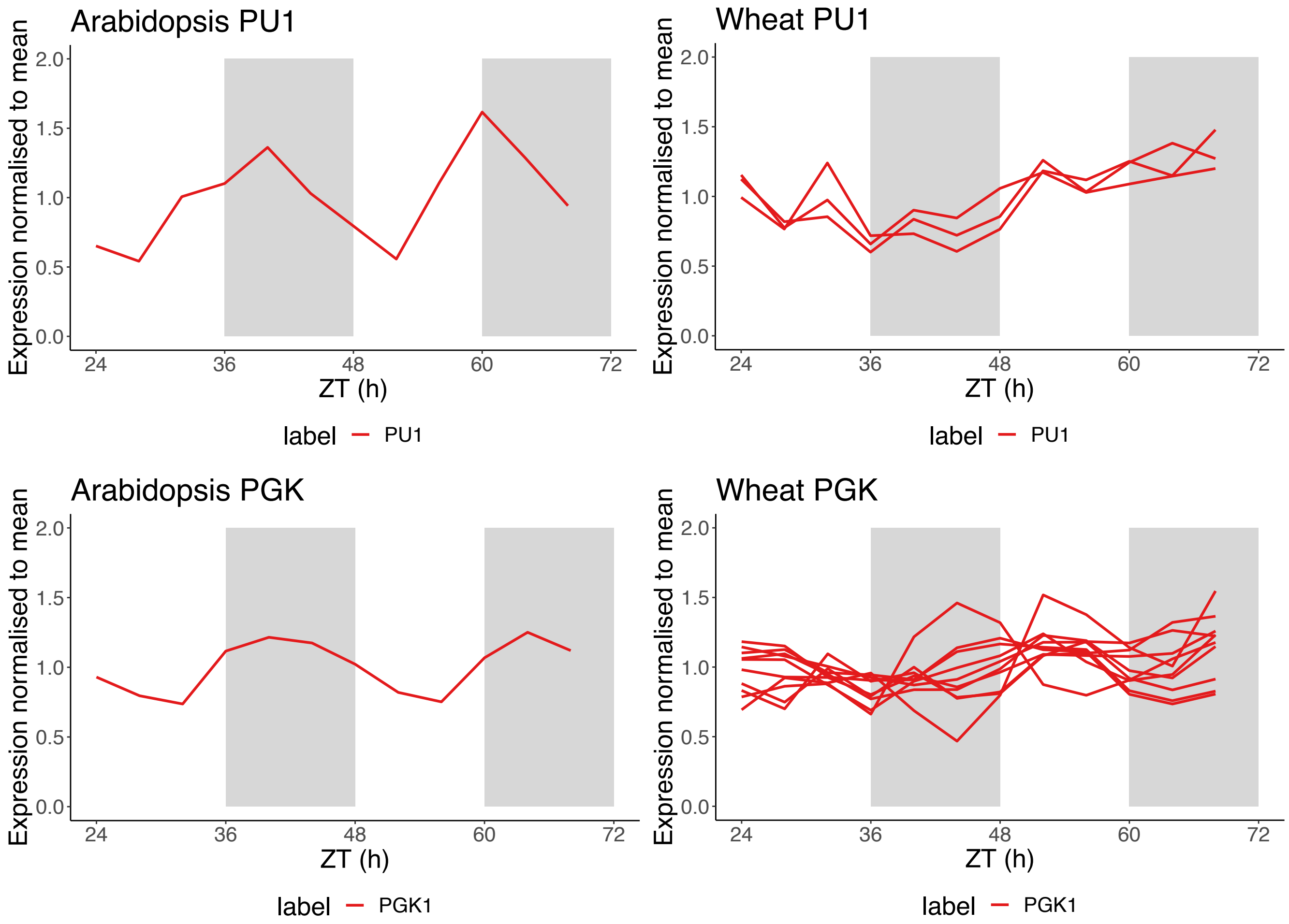


**Supplementary Figure 20.** ***Arabidopsis*** **and wheat primary metabolism genes under free-running L:L conditions.** Data is mean normalised TPM. Shaded white and grey backgrounds indicate perceived day and night periods respectively. Gene IDs for all genes plotted can be seen in Supplementary Table 8.

**Supplementary Table 1: Proportions of rhythmically classified genes using datasets filtered to remove low expression genes.** Numbers of rhythmic genes at ($q < 0.05$ or $q < 0.01$) in *Arabidopsis* and wheat identified using Metacycle Benjamini Hochberg q-values. Data windows reflect hours relative to transfer to constant light. Metacycle was used on expression datasets with genes removed that had fewer than 0.1 TPM in more than 6 timepoints. The effects of filtering data to remove low expression data are discussed in Supplementary Note 1.

| | Wheat data from this study | | *Arabidopsis* data from Romanowski et al. |
|---|---|---|---|
| | 24-68 data window | 0-68 data window | 24-68 data window |
| Total number of expressed genes | 61,233 | 62,624 | 22,020 |
| Total rhythmic genes (q<0.05) | 28,359 | 27,781 | 13,550 |
| Total rhythmic genes (q<0.01) | 19,384 | 21,006 | 11,755 |
| Mean Period (h) (q<0.05) | 26.67 (SD 3.48) | 26.85 (SD 2.66) | 23.49 (SD 2.47) |
| Mean Period (h) (q<0.01) | 26.85 (SD 3.14) | 26.87 (SD 2.34) | 23.57 (SD 2.19) |
| Mean relative Amplitude (q<0.05) | 0.25 (SD 0.18) | 0.27 (0.20) | 0.28 (SD 0.20) |
| Mean relative Amplitude (q<0.01) | 0.28 (SD 0.19) | 0.30 (SD 0.21) | 0.30 (SD 0.21) |

**Supplementary Table 2: Meta2d estimates from previously published circadian datasets.** Estimates are based on gene-level TPM normalised transcripts from a data window of 24-68h relative to transfer to constant light for each species. Rhythmic genes are defined as having a q-value < 0.01 and period and rAMP means are for rhythmic genes only.

| Species | Expressed genes | rhythmic genes | rhythmic genes % | Mean BH. *q*-value | SD BH. *q*-value | Mean Period | SD Period | Mean rAMP | SD rAMP |
|---|---|---|---|---|---|---|---|---|---|
| *Brassica rapa* | 42,186 | 14,783 | 35.04 | 0.0007 | 0.0018 | 23.54 | 1.42 | 0.43 | 0.25 |
| *Arabidopsis thaliana* | 26,392 | 10,317 | 39.09 | 0.0017 | 0.0024 | 23.62 | 2.04 | 0.31 | 0.21 |
| *Glycine max* | 43,165 | 20,347 | 47.14 | 0.0012 | 0.0022 | 24.65 | 1.49 | 0.35 | 0.25 |
| *Brachypodium distachyon* | 22,565 | 434 | 1.92 | 0.0038 | 0.0029 | 25.60 | 1.01 | 0.30 | 0.17 |
| *Triticum aestivum* | 86,567 | 18,633 | 21.52 | 0.0024 | 0.0026 | 26.82 | 3.21 | 0.28 | 0.20 |

**Supplementary Table 3: GO-slim terms for biological processes associated with similar patterns of circadian expression or different patterns of circadian expression in Brassica rapa paralogs.** Only enriched terms which were highly enriched (Fisher's exact test p<0.01) in one category and non-significantly expressed (p>0.05) in the other category are displayed.

| | GO.ID | Term | *p*-value in circadian similar paralogs | *p*-value in circadian different paralogs |
|---|---|---|---|---|
| Circadian similar patterns | GO:0015979 | photosynthesis | 0.00 | 0.98 |
| | GO:0006091 | generation of precursor metabolites and energy | 0.00 | 0.13 |
| | GO:0009056 | catabolic process | 0.00 | 0.37 |
| Circadian differential patterns | GO:0005975 | carbohydrate metabolic process | 0.07 | 0.00 |
| | GO:0006629 | lipid metabolic process | 0.45 | 0.00 |
| | GO:0006725 | cellular aromatic compound metabolic process | 0.51 | 0.00 |
| | GO:0046483 | heterocycle metabolic process | 0.51 | 0.00 |

**Supplementary Table 4: Expected and Observed runs of Rhythmic triad categories across the wheat genome.** Rhythmic unbalanced triads were classified as having one or two rhythmic homoeologs q <0.01 where the other homoeolog has a q >0.05. Triads with balanced rhythmicity have three rhythmic homoeologs q<0.05. In total, chromosome position data was found for 4225 of these triads. Expected maximum run lengths for each category were calculated as described in Supplementary Note 5. For Category test 3, Triads lacking rhythmicity on the A chromosome are either: triads with 1 rhythmic homeolog on the B sub-genome, triads with 1 rhythmic homoeolog on the D sub-genome or triads with two rhythmic homoeologs on the B and D subgenomes. The same logic applies to the other two categories. *P* -values for randomness in each category set were computed using a Wald–Wolfowitz test using publicly available code: https://github.com/psinger/RunsTest and implemented in Python.

| | Count | % | Expected max run length | Observed run length | Observed run length SD | Difference O-E | *p*-value Wald–Wolfowitz test: |
|---|---|---|---|---|---|---|---|
| Category test set 1: runs of numbers of rhythmic genes | | | | | | | |
| Triads with 1 rhythmic homoeolog | 2058 | 0.30 | 7.31 | 7 | 0.80 | -0.31 | 0.156 |
| Triads with 2 rhythmic homoeologs | 1392 | 0.20 | 5.52 | 5 | 0.58 | -0.52 | |
| Triads with 3 rhythmic homoeologs | 3448 | 0.50 | 12.75 | 11 | 1.41 | -1.75 | |
| Category test set 2: runs of numbers of rhythmic genes on specific subgenomes | | | | | | | |
| Triads with 1 rhythmic homoeolog on the A chromosome | 725 | 0.11 | 3.92 | 3 | 0.36 | -0.92 | 0.242 |
| Triads with 1 rhythmic homoeolog on the B chromosome | 716 | 0.10 | 3.90 | 4 | 0.38 | 0.10 | |

| | | | | | | | |
|---|---|---|---|---|---|---|---|
| Triads with 1 rhythmic homoeolog on the D chromosome | 617 | 0.09 | 3.66 | 3 | 0.34 | -0.66 | |
| Triads with 2 rhythmic homoeologs on the A and B chromosomes | 432 | 0.06 | 3.19 | 4 | 0.29 | 0.81 | |
| Triads with 2 rhythmic homoeologs on the A and D chromosomes | 490 | 0.07 | 3.34 | 3 | 0.29 | -0.34 | |
| Triads with 2 rhythmic homoeologs on the B and D chromosomes | 470 | 0.07 | 3.29 | 3 | 0.27 | -0.29 | |
| Triads with 3 rhythmic homoeologs on the A, B and D chromosomes | 3448 | 0.50 | 12.75 | 11 | 1.41 | -1.75 | |
| Category test set 3: runs of triads lacking rhythmicity on a particular sub-genome | | | | | | | |
| Triads lacking rhythmicity on the A chromosome | 1803 | 0.261 | 6.59 | 7 | 0.71 | 0.41 | 0.198 |
| Triads lacking rhythmicity on the B chromosome | 1832 | 0.266 | 6.67 | 6 | 0.75 | -0.67 | 0.056 |
| Triads lacking rhythmicity on the D chromosome | 1873 | 0.272 | 6.78 | 6 | 0.72 | -0.78 | 0.166 |

**Supplementary Table 5. Characteristics of circadian rhythms in wheat genes split into two hour period bins.** Comparison of rhythm *q*-values, baseline expression, relative amplitudes and relative amplitude error for each of the five period groups over a data window of 24-68h. Period groups were defined as having Metacycle q-values of <0.01, and period categories were confirmed with independent estimates from FFT-NLLS (Biodare2). Period categories are as follows: Group A: 21-23h, Group B: 23-25h, Group C: 25-27h, Group D 27-29, Group E 29-30. Statistics were calculated in Metacycle and RAE scores in Biodare2.

| Group.ID | N | Mean BH.Q | Sd BH.Q | Mean LOG10(base) | Sd LOG10(base) | Mean rAMP | Sd rAMP | Mean RAE | Sd RAE |
|---|---|---|---|---|---|---|---|---|---|
| Group_A | 808 | 0.00317 | 0.00282 | 0.255 | 0.809 | 0.224 | 0.142 | 0.669 | 0.15 |
| Group_B | 2229 | 0.00215 | 0.00248 | 0.627 | 0.784 | 0.3 | 0.194 | 0.667 | 0.16 |
| Group_C | 2506 | 0.00167 | 0.00221 | 0.712 | 0.781 | 0.385 | 0.257 | 0.667 | 0.15 |
| Group_D | 1940 | 0.00208 | 0.00241 | 0.677 | 0.8 | 0.285 | 0.166 | 0.69 | 0.13 |
| Group_E | 761 | 0.00232 | 0.0026 | 0.567 | 0.851 | 0.234 | 0.136 | 0.704 | 0.13 |

**Supplementary Table 6. Circadian characteristics of module eigengenes.** Expression data (TPM) over a time window of 24-68h after transfer to constant light (L:L) was filtered for rhythmicity (BH $q < 0.01$) before clustering in WGCNA. Eigengene expression values were analysed using Metacycle to approximate phase and period values.

| Species | Module ID | Number of genes | Absolute phase (ZT) from Metacycle (h) | Circadian phase (CT) from Metacycle (h) | Metacycle Period length (h) |
|---|---|---|---|---|---|
| *Arabidopsis* | A1 | 295 | 1.93 | 1.94 | 23.93 |
| *Arabidopsis* | A2 | 133 | 5.90 | 6.19 | 22.87 |
| *Arabidopsis* | A3 | 388 | 7.23 | 6.87 | 25.28 |
| *Arabidopsis* | A4 | 697 | 10.95 | 11.06 | 23.75 |
| *Arabidopsis* | A5 | 4,302 | 14.16 | 14.46 | 23.50 |
| *Arabidopsis* | A6 | 2,221 | 17.52 | 18.43 | 22.82 |
| *Arabidopsis* | A7 | 696 | 19.86 | 20.41 | 23.35 |
| *Arabidopsis* | A8 | 1,020 | 21.51 | 22.64 | 22.80 |
| *Arabidopsis* | A9 | 358 | 0.32 | 0.33 | 23.31 |
| Wheat | W1 | 806 | 2.12 | 1.74 | 29.18 |
| Wheat | W2 | 720 | 2.29 | 2.24 | 24.55 |
| Wheat | W3 | 351 | 7.44 | 6.34 | 28.18 |
| Wheat | W4 | 3,020 | 11.35 | 9.96 | 27.37 |
| Wheat | W5 | 3,369 | 15.41 | 12.69 | 29.13 |
| Wheat | W6 | 373 | 15.89 | 11.81 | 32.27 |
| Wheat | W7 | 2,634 | 19.90 | 17.95 | 26.62 |
| Wheat | W8 | 945 | 20.89 | 22.17 | 22.62 |
| Wheat | W9 | 4,106 | 22.95 | 22.15 | 24.87 |

**Supplementary Table 8: Gene IDs for core circadian genes and corresponding *Arabidopsis* orthologs**. Protein family trees correspond to Supplementary Figures 12-16.

| Wheat Gene ID RefSeqv1 | Suggested wheat gene name | Protein family tree | Most like *Arabidopsis* gene ID(s) | |
|---|---|---|---|---|
| TraesCS7A02G299400 | TaLHY_7A | Tree 1: MYB (LHY-like) | AT1G01060/AT2G46830 | LHY/CCA1 |
| TraesCS7D02G295400 | TaLHY_7D | Tree 1: MYB (LHY-like) | AT1G01060/AT2G46830 | LHY/CCA1 |
| TraesCS7B02G188000 | TaLHY_7B | Tree 1: MYB (LHY-like) | AT1G01060/AT2G46830 | LHY/CCA1 |
| TraesCS6B02G253900 | TaTOC1_6B | Tree 2: PRR (TOC1-like) | AT5G61380 | TOC1 |
| TraesCS6A02G227900 | TaTOC1_6A | Tree 2: PRR (TOC1-like) | AT5G61380 | TOC1 |
| TraesCS6D02G207100 | TaTOC1_6D | Tree 2: PRR (TOC1-like) | AT5G61380 | TOC1 |
| TraesCS5D02G326200 | TaPRR59_5D | Tree 2: PRR (TOC1-like) | AT2G46790/AT5G24470 | PRR5/PRR9 |

| | | | | |
|---|---|---|---|---|
| TraesCS5A02G320300 | TaPRR59_5A | Tree 2: PRR (TOC1-like) | AT2G46790/AT5G24470 | PRR5/PRR9 |
| TraesCS5B02G320500 | TaPRR59_5B | Tree 2: PRR (TOC1-like) | AT2G46790/AT5G24470 | PRR5/PRR9 |
| TraesCS4B02G115100 | TaPRR95_4B | Tree 2: PRR (TOC1-like) | AT2G46790/AT5G24470 | PRR5/PRR9 |
| TraesCS4D02G112800 | TaPRR95_4D | Tree 2: PRR (TOC1-like) | AT2G46790/AT5G24470 | PRR5/PRR9 |
| TraesCS4B02G198700 | TaPRR73_4B | Tree 2: PRR (TOC1-like) | AT5G60100/AT5G02810 | PRR3/PRR7 |
| TraesCS4A02G105300 | TaPRR73_4A | Tree 2: PRR (TOC1-like) | AT5G60100/AT5G02810 | PRR3/PRR7 |
| TraesCS4D02G199600 | TaPRR73_4D | Tree 2: PRR (TOC1-like) | AT5G60100/AT5G02810 | PRR3/PRR7 |
| TraesCS2A02G081900 | TaPRR37_2A (Ppd) | Tree 2: PRR (TOC1-like) | AT5G60100/AT5G02810 | PRR3/PRR7 |
| TraesCS2D02G079600 | TaPRR37_2D (Ppd) | Tree 2: PRR (TOC1-like) | AT5G60100/AT5G02810 | PRR3/PRR7 |
| TraesCSU02G221500 | TaPRR37_2B (Ppd) | Tree 2: PRR (TOC1-like) | AT5G60100/AT5G02810 | PRR3/PRR7 |

| | | | | |
|---|---|---|---|---|
| TraesCSU02G196100* | TaPRR37_2B (Ppd) | Tree 2: PRR (TOC1-like) | AT5G60100/AT5G02810 | PRR3/PRR7 |
| TraesCSU02G199500 | TaPRR37_2B (Ppd) | Tree 2: PRR (TOC1-like) | AT5G60100/AT5G02810 | PRR3/PRR7 |
| TraesCS6B02G426300 | TaADOa_6B1 | Tree 3: PAS/LOV (ZTL-like) | AT5G57360/AT2G18915 | ZTL/LKP2 |
| TraesCS7A02G431600 | TaADOa_7A | Tree 3: PAS/LOV (ZTL-like) | AT5G57360/AT2G18915 | ZTL/LKP2 |
| TraesCS7D02G423400 | TaADOa_7D | Tree 3: PAS/LOV (ZTL-like) | AT5G57360/AT2G18915 | ZTL/LKP2 |
| TraesCS6B02G149800 | TaADOb_6B2 | Tree 3: PAS/LOV (ZTL-like) | AT5G57360/AT2G18915 | ZTL/LKP2 |
| TraesCS6D02G111600 | TaADOb_6D | Tree 3: PAS/LOV (ZTL-like) | AT5G57360/AT2G18915 | ZTL/LKP2 |
| TraesCS6A02G121500 | TaADOb_6A | Tree 3: PAS/LOV (ZTL-like) | AT5G57360/AT2G18915 | ZTL/LKP2 |
| TraesCS4A02G164000 | TaFKF1_4A | Tree 3: PAS/LOV (ZTL-like) | AT1G68050 | FKF1 |
| TraesCS4B02G157500 | TaFKF1_4B | Tree 3: PAS/LOV (ZTL-like) | AT1G68050 | FKF1 |

| | | | | |
|---|---|---|---|---|
| TraesCS3B02G394200 | TaLUX-La_3B | Tree 4: MYB (LUX-like) | AT3G46640/AT5G59570 | LUX/BOA |
| TraesCS3D02G355500 | TaLUX-La_3D | Tree 4: MYB (LUX-like) | AT3G46640/AT5G59570 | LUX/BOA |
| TraesCS3A02G361800 | TaLUX-La_3A | Tree 4: MYB (LUX-like) | AT3G46640/AT5G59570 | LUX/BOA |
| TraesCS3A02G526600 | TaLUX/BOA_3A | Tree 4: MYB (LUX-like) | AT3G46640/AT5G59570 | LUX/BOA |
| TraesCS3D02G531900 | TaLUX/BOA_3D | Tree 4: MYB (LUX-like) | AT3G46640/AT5G59570 | LUX/BOA |
| TraesCS3B02G594300 | TaLUX/BOA_3B | Tree 4: MYB (LUX-like) | AT3G46640/AT5G59570 | LUX/BOA |
| TraesCS1A02G258400 | TaLUX-Lb_1A | Tree 4: MYB (LUX-like) | AT3G46640/AT5G59570 | LUX/BOA |
| TraesCS1B02G268900 | TaLUX-Lb_1B | Tree 4: MYB (LUX-like) | AT3G46640/AT5G59570 | LUX/BOA |
| TraesCS1D02G257700 | TaLUX-Lb_1D | Tree 4: MYB (LUX-like) | AT3G46640/AT5G59570 | LUX/BOA |
| TraesCS5B02G200500 | TaELF4-L_5B | Tree 5: ELF4-like | AT2G40080/AT1G72630/AT1G17455 | ELF4/ELF4-L2/ELF4-L4 |
| TraesCS5A02G201900 | TaELF4-L_5A | Tree 5: ELF4-like | AT2G40080/AT1G72630/AT1G17455 | ELF4/ELF4-L2/ELF4-L4 |
| TraesCS5D02G208200 | TaELF4-L_5D | Tree 5: ELF4-like | AT2G40080/AT1G72630/AT1G17455 | ELF4/ELF4-L2/ELF4-L4 |

| | | | | |
|---|---|---|---|---|
| TraesCS4B02G149300 | TaELF4-L_4B | Tree 5: ELF4-like | AT2G40080/AT1G72630/AT1G17455 | ELF4/ELF4-L2/ELF4-L4 |
| TraesCS4D02G149000 | TaELF4-L_4D | Tree 5: ELF4-like | AT2G40080/AT1G72630/AT1G17455 | ELF4/ELF4-L2/ELF4-L4 |
| TraesCS1D02G451200 | TaELF3_1D (Eps-D1) | BLASTP for ABL11477 | AT2G25930 | ELF3 |
| TraesCS1A02G443200 | TaELF3_1A | BLASTP for ABL11477 | AT2G25930 | ELF3 |
| TraesCS1B02G477400 | TaELF3_1B | BLASTP for ABL11477 | AT2G25930 | ELF3 |
| TraesCS3B02G135400 | TaGI_3B | BLASTP for AAQ11738 | AT1G22770 | GI |
| TraesCS3A02G116300 | TaGI_3A | BLASTP for AAQ11738 | AT1G22770 | GI |
| TraesCS3D02G118200 | TaGI_3D | BLASTP for AAQ11738 | AT1G22770 | GI |
| TraesCS2B02G478700 | TaRVE27a_2B | Tree 1: MYB (LHY-like) | AT5G17300/AT5G37260/AT1G18330 | RVE1/RVE2/RVE7 |
| TraesCS2A02G456600 | TaRVE27a_2A | Tree 1: MYB (LHY-like) | AT5G17300/AT5G37260/AT1G18330 | RVE1/RVE2/RVE7 |
| TraesCS2D02G456900 | TaRVE27a_2D | Tree 1: MYB (LHY-like) | AT5G17300/AT5G37260/AT1G18330 | RVE1/RVE2/RVE7 |

| | | | | |
|---|---|---|---|---|
| TraesCS7A02G553800 | TaRVE27b_7A | Tree 1: MYB (LHY-like) | AT5G17300/AT5G37260/AT1G18330 | RVE1/RVE2/RVE7 |
| TraesCS7B02G478200 | TaRVE27b_7B | Tree 1: MYB (LHY-like) | AT5G17300/AT5G37260/AT1G18330 | RVE1/RVE2/RVE7 |
| TraesCS7D02G540700 | TaRVE27b_7D | Tree 1: MYB (LHY-like) | AT5G17300/AT5G37260/AT1G18330 | RVE1/RVE2/RVE7 |
| TraesCS6D02G241900 | TaRVE27c_6D | Tree 1: MYB (LHY-like) | AT5G17300/AT5G37260/AT1G18330 | RVE1/RVE2/RVE7 |
| TraesCS6B02G288500 | TaRVE27c_6B | Tree 1: MYB (LHY-like) | AT5G17300/AT5G37260/AT1G18330 | RVE1/RVE2/RVE7 |
| TraesCS6A02G261100 | TaRVE27c_6A | Tree 1: MYB (LHY-like) | AT5G17300/AT5G37260/AT1G18330 | RVE1/RVE2/RVE7 |
| TraesCS6B02G266800 | TaRVE68_6B | Tree 1: MYB (LHY-like) | AT5G52660/AT3G09600 | RVE6/RVE3/RVE5/RVE4/RVE8 |
| TraesCS6D02G239800 | TaRVE68_6D | Tree 1: MYB (LHY-like) | AT5G52660/AT3G09600 | RVE6/RVE3/RVE5/RVE4/RVE8 |
| TraesCS6A02G258600 | TaRVE68_6A | Tree 1: MYB (LHY-like) | AT5G52660/AT3G09600 | RVE6/RVE3/RVE5/RVE4/RVE8 |
| TraesCS4A02G474100 | TaRVE86a_4A | Tree 1: MYB (LHY-like) | AT5G52660/AT3G09600 | RVE6/RVE3/RVE5/RVE4/RVE8 |

| | | | | |
|---|---|---|---|---|
| TraesCS7D02G014900 | TaRVE86a_7D | Tree 1: MYB (LHY-like) | AT5G52660/AT3G09600 | RVE6/RVE3/RVE5/RVE4/RVE8 |
| TraesCS7A02G017600 | TaRVE86a_7A | Tree 1: MYB (LHY-like) | AT5G52660/AT3G09600 | RVE6/RVE3/RVE5/RVE4/RVE8 |
| TraesCS7D02G458000 | TaRVE86b_7D | Tree 1: MYB (LHY-like) | AT5G52660/AT3G09600 | RVE6/RVE3/RVE5/RVE4/RVE8 |
| TraesCS7A02G470700 | TaRVE86b_7A | Tree 1: MYB (LHY-like) | AT5G52660/AT3G09600 | RVE6/RVE3/RVE5/RVE4/RVE8 |
| TraesCS7B01G372500 | TaRVE86b_7B | Tree 1: MYB (LHY-like) | AT5G52660/AT3G09600 | RVE6/RVE3/RVE5/RVE4/RVE8 |
| TraesCS4D02G166000 | TaLNK1_4D | BLASTP for Bradi1g60846 and AT5G64170 | AT5G64170 | LNK1 |
| TraesCS4A02G162200 | TaLNK1_4A | BLASTP for Bradi1g60846 and AT5G64170 | AT5G64170 | LNK1 |
| TraesCS4B02G153700 | TaLNK1_4B | BLASTP for Bradi1g60846 and AT5G64170 | AT5G64170 | LNK1 |

| | | | | |
|---|---|---|---|---|
| TraesCS3A02G195700 | TaLNK2_3A | BLASTP for Bradi2g13610 and AT3G54500 | AT3G54500 | LNK2 |
| TraesCS3B02G227900 | TaLNK2_3B | BLASTP for Bradi2g13610 and AT3G54500 | AT3G54500 | LNK2 |
| TraesCS6D02G397000 | TaCHEa_6D | BLASTP for Braidi3g60352 and At5g08330 | At5g08330 (che)/ AT5G23280 | CHE/TCP7 |
| TraesCS6A02G412800 | TaCHEa_6A | BLASTP for Braidi3g60352 and At5g08330 | At5g08330 (che)/ AT5G23280 | CHE/TCP7 |
| TraesCS6B02G462100 | TaCHEa_6B | BLASTP for Braidi3g60352 and At5g08330 | At5g08330 (che)/ AT5G23280 | CHE/TCP7 |
| TraesCS6B02G262600 | TaCHEb_6B | BLASTP for Braidi3g60352 and At5g08330 | At5g08330 (che)/ AT5G23280 | CHE/TCP7 |
| TraesCS6A02G233800 | TaCHEb_6A | BLASTP for Braidi3g60352 and At5g08330 | At5g08330 (che)/ AT5G23280 | CHE/TCP7 |

| | | | | |
|---|---|---|---|---|
| TraesCS6D02G216100 | TaCHEb_6D | BLASTP for Braidi3g60352 and At5g08330 | At5g08330 (che)/ AT5G23280 | CHE/TCP7 |
| TraesCS6A02G193300 | TaLWD_6A | BLASTP for Bradi3g44580, At1g12910 and AT3G26640 | At1g12910 (LWD1)/ AT3G26640 (LWD2) | LWD1/LWD2 |
| TraesCS6B02G221400 | TaLWD_6B | BLASTP for Bradi3g44580, At1g12910 and AT3G26640 | At1g12910 (LWD1)/ AT3G26640 (LWD2) | LWD1/LWD2 |
